# Robust chromatin state annotation

**DOI:** 10.1101/2023.07.15.549175

**Authors:** Mehdi Foroozandeh Shahraki, Marjan Farahbod, Maxwell Libbrecht

## Abstract

**Background:** Segmentation and genome annotations (SAGA) methods such as ChromHMM and Segway are widely to annotate chromatin states in the genome. These algorithms take as input a collection of genomics datasets, partition the genome, and assign a label to each segment such that positions with the same label have similar patterns in the input data. SAGA methods output an human-interpretable summary of the genome by labeling every genomic position with its annotated activity such as Enhancer, Transcribed, etc. Chromatin state annotations are essential for many genomic tasks, including identifying active regulatory elements and interpreting disease-associated genetic variation. However, despite the widespread applications of SAGA methods, no principled approach exists to evaluate the statistical significance of SAGA state assignments.

**Results:** Towards the goal of producing robust chromatin state annotations, we performed a comprehensive evaluation of the reproducibility of SAGA methods. We show that SAGA annotations exhibit a large degree of disagreement, even when run with the same method on replicated data sets. This finding suggests that there is significant risk to using SAGA chromatin state annotations.

To remedy this problem, we introduce SAGAconf, a method for assigning a measure of confidence (r-value) to SAGA annotations. This r-value is assigned to each genomic bin of a SAGA annotation and represents the probability that the label of this bin will be reproduced in a replicated experiment. This process is analogous to irreproducible discovery rate (IDR) analysis that is commonly used for ChIP-seq peak calling and related tasks. Thus SAGAconf allows a researcher to select only the reliable parts of a SAGA annotation for use in downstream analyses.

SAGAconf r-values provide accurate confidence estimates of SAGA annotations, allowing researchers to filter out unreliable elements and remove doubt in those that stand up to this scrutiny.

## 1 Background

Annotating regulatory elements in the genome is fundamental to answering key questions including the molecular basis of disease, evolution, cellular differentiation, and development. To this end, international mapping projects have recently measured epigenetic activity in hundreds of cell and tissue types using genome-wide assays such as ChIP-seq [1, 2].

Chromatin state annotations produced by segmentation and genome annotation (SAGA) methods have emerged as the predominant way to summarize epigenomic data sets in order to annotate the genome. SAGA methods include Segway [3] and ChromHMM [4] and others [5, 6, 7, 8] (reviewed in [9]). They take a collection epigenomic assay data sets from a given cell type or tissue as input and partition the genome into segments with similar patterns in the input data sets. The output is an annotation that assigns a label to each genomic position. These algorithms use probabilistic graphical models such as Hidden Markov Models. They are unsupervised in the sense that the model identifies patterns in the data and the researcher later maps each patterns to a putative biological functions such as enhancer, promoter or transcribed gene. Thus, SAGA algorithms are used to distill complex data into an interpretable summary of genomic activity. SAGA algorithms have been broadly applied. Large-scale epigenome mapping projects such as ENCODE [1] and Roadmap [2] produced them as their primary output, and researchers have now produced reference SAGA annotations for hundreds of cell types [10, 11, 12, 13]

Previous efforts to evaluate the reliability of SAGA chromatin state annotations have found mixed results. On one hand, SAGA annotations recapitulate known genome biology and the annotations accurately represent many genomic phenomena including transcription [7, 10, 9]. However, results are often dissimilar between SAGA methods and hyper-parameter settings of a given method [14, 10, 7, 9]. This suggests that some aspects of SAGA annotations do not reflect true biology.

Thus, there is a great need for a way to produce robust chromatin state annotations. Unfortunately, currently there is no principled way to evaluate the statistical significance of SAGA label assignments. SAGA annotations are not the result of a statistical test, so they do not carry a p-value. SAGA methods use probabilistic graphical models which output posterior probabilities; however, in practice these posterior probabilities are vastly overconfident, resulting in most positions receiving posterior probability *>*99% [15, 9].

Here, we propose the first method for assigning reliable confidence scores to SAGA annotations. We base our approach on evaluating the reproducibility of annotations across replicates [16]. We are motivated by methods for ChIP-seq peak calling, for which researchers generally use irreproducible discovery rate (IDR) analysis to assign confidence scores to peaks [17, 18, 19, 20]. In IDR analysis, researchers score putative peaks according to their reproducibility in two or more experimental replicates, with the expectation that peaks findings are expected to consistently rank highly in both experiments. Note that, although several methods have been developed for the related task of comparing sets of SAGA annotations to one another, to our knowledge no existing method can assign confidence estimates to a single chromatin state annotation. In particular, ChromDiff [21], SCIDDO [22] and Epi-Compare [23] perform the task of group-wise comparative analysis, in which they take as input two sets of annotations and identify differential labels between the two sets. Related method EpiAlign [24] and Chromswitch [25] similarly perform group-wise comparative analysis but are primarily designed to compare chromatin state patterns within a query region. CSREP [26] performs the task of group summarization, taking annotations for a group of samples as input and probabilistically estimating the state at each genomic position to derive a representative chromatin state map for the group. Because all of these existing studies analyze patterns across annotations of many cell types, they cannot estimate the confidence in the annotation of a single target cell type.

Towards the goal of understanding robustness of SAGA annotations, here we perform a comprehensive evaluation of the reproducibility of SAGA methods. As described below, we show that SAGA annotations exhibit a large degree of disagreement, even when run with the same method on replicated data sets. Much of this disagreement can be attributed to unimportant factors such as the granularity of label definitions and mismatch in the boundaries of annotated elements. Yet, much disagreement remains, suggesting that a substantial fraction of element annotations produced by SAGA cannot be relied upon as they are not reproduced by a replicate analysis.

To remedy this problem, we introduce SAGAconf, a method for assigning a measure of confidence (*r*-value) to SAGA annotations. This *r*-value is assigned to each genomic bin of a SAGA annotation and represents the probability that the label of this bin will be reproduced in a replicated experiment. Thus, SAGAconf allows a researcher to select only the reliable parts of a SAGA annotation for use in downstream analyses.

## 2 Results

### 2.1 Comprehensive evaluation of chromatin state reproducibility

We performed comprehensive evaluation of reproducibility of SAGA annotations (Methods). We collected replicated pairs of epigenomic data sets in five cell types. In each replicate pair, we applied a SAGA pipeline to produce two replicate annotations, which we term the “base” and “verification” annotation, respectively. For the purposes of exposition, we focus our analysis on a running example: ChromHMM run on GM12878 (under setting 1 described below: different data, different models). A SAGA model assigns an integer ID (1..*k*) to each genomic bin. These IDs represent different chromatin states, also known as “states” (Figure 1A). The relationship of a pair of annotations can be characterized by a *k × k* matrix representing the frequency of overlap of each base state with each verification state (Figure 1B). Since SAGA methods are unsupervised, the output integer state IDs do not directly correspond across two SAGA models. Thus, we must first detect corresponding states across annotations to enable comparison (Figure 1C and Section 4.4). To establish a correspondence between chromatin states while accounting for varying genomic coverages, we calculate intersection over union (IoU) of overlap (Methods 4.4, Figure 1C) and, for each base state, we define the corresponding verification state to be the one with highest IoU (Figure 1A,C).

**Figure 1.**
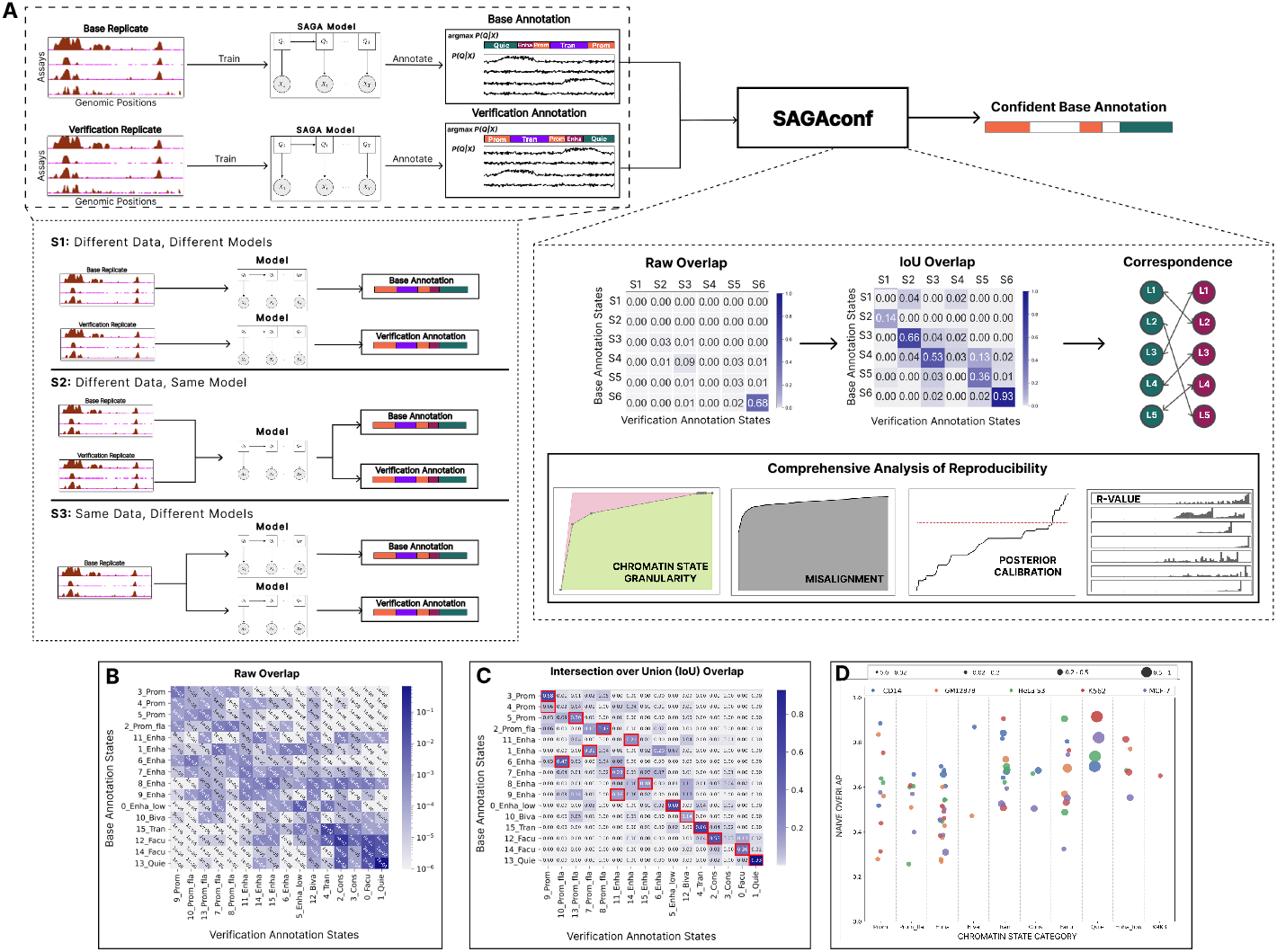
Schematic workflow. (A) We obtained histone modification assays from biological replicates via ENCODE DCC [1] and used these data sets to train SAGA models (Segway or ChromHMM) to generate chromatin state annotations. The SAGA model outputs a matrix representing the posterior probability *P* (*Q* |*X*) values of assigning each chromatin state to each genomic position and a vector of state labels assigned to the position with the highest posterior probability argmax*P* (*Q* |*X*) (Methods). One set of replicated data is chosen as the base and the other as the verification. SAGA training and genome annotation are performed according to three settings of variability: S1 (different Data, different models), where two separate SAGA models are trained independently using data from each biological replicate; S2 (different Data, same model), where data from both replicates are concatenated to train a single SAGA model that provides separate annotations for each replicate; and S3 (same Data, different models), where the same dataset (base replicate) is used to train two different SAGA models with different parameter initializations. Both the base and verification annotations, generated by any variability setting, are inputs to SAGAconf. The SAGAconf evaluation pipeline begins by forming a pairwise overlap frequency distribution matrix between the two annotations and calculating the intersection over union (IoU) overlap to determine correspondence between state pairs across the annotations. SAGAconf performs a comprehensive reproducibility analysis and outputs a subset of the base annotation that it identifies as confident (Methods). (B) The raw overlap frequency distribution from our running example annotation (setting 1, ChromHMM, GM12878). Rows and columns correspond to states in base and verification annotations, respectively. Color indicates frequency of overlap (log scale). (C) Same as (B), but color indicates the intersection over Union (IoU) of overlap is derived from raw overlap matrix (Methods, linear scale). For each chromatin state of the base annotation, its corresponding state in the verification annotation is defined as the one with the maximum IoU (marked with red square). (D) Fraction of overlap (Naive overlap) of various chromatin states categories identified in the ChromHMM annotations according to S1 (different Data, different models) for all five cell types. Each dot represents a chromatin state, with color denoting cell type and size proportional to genome coverage.

Irreproducibility between annotations may be caused either by differences in input data replicates or differences in model training. To delineate among these sources of irreproducibility, we assess pairs of experiments in three different settings of variability (Figure 1A, Section 4.11). In setting 1 (different data, different models) we trained two separate SAGA models using data from separate biological replicates, simulating the case where independent researchers each perform a SAGA analysis. In setting 2 (different data, same model), we train a single SAGA model and use it to annotate each replicate (known as “concatenated” annotation). This setting isolates irreproducibility due to differences in input data replicates. In setting 3 (same data, different models), the same dataset from one of the biological replicates is used to train two different SAGA models, but with a different random parameter initialization in each model. This setting isolates irreproducibility due to model training.

Since SAGA methods are unsupervised, each integer state label must be assigned a putative biological interpretation such as “Promoter” or “Enhancer”. To avoid bias stemming from manual interpretation, we used a previously-described automatic process to assign a vocabulary of human-readable chromatin state categories [10] (Supplementary Section 1.3). Note that interpretations are for exposition only and all analysis was performed at the level of states, not interpreted chromatin state categories. Because it is automated, the interpretation process may be imperfect; for example, in our running example, base state 6_Enha overlaps with verification state 10_Prom_fla; the mismatch in chromatin state category likely results from an error in the interpreter, not the annotations themselves.

### 2.2 Pairwise overlap does not fully capture the reproducibility profile of SAGA annotations

We found substantial differences between annotations of the two replicates (Figure 1 B,C,D). Overall, in our running example, only *∼*80% of genomic bins are annotated with the corresponding label in the verification annotation (Figure 3 H).

Overlap is poorest for punctate chromatin states such as Promoter, Enhancer and Bivalent (*∼*50% overlap, Figure 1 D). Overlap for broad chromatin states such as Transcribed, Facultative and Constitutive Heterochromatin is higher (*∼*70% overlap, Figure 1 D) and best for Quiescent (*∼*80% overlap, Figure 1 D). These results are generally consistent across the five cell types we tested (Figure 3 H). Overlap is significantly lower for Segway, which achieves just 30-40% average overlap (Figure 3 H). These results are concerning for the application of SAGA annotations, as reproducibility is far lower than the 95% standard used for most genomic predictions. Results are similar in other cell types and settings (Supplementary Section 2). In setting 2 (different data, same model), we observe a slight improvement in the naive overlap, over setting 1 (different model, different data), for ChromHMM annotations, but not for Segway (Figure 3H). However, in setting 3, where we used the same data with different models, we observed a significant improvement in the average overlap, especially for Segway annotations (Figure 3H). This suggests that the quality of data plays a more significant role in the poor naive overlap observed in settings 1 and 2 than the model training.

However, as we describe below, measuring reproducibility according to the naive overlap with the corresponding states may be too conservative, as doing so counts two types of variation that may not be important to practitioners.

### 2.3 Granularity of chromatin states affect reproducibility

Each base state generally overlaps with multiple verification states, yet this overlap usually occurs among a small number of related states (Figure 1C). For example, in our running example, base state 7_Enha frequently overlaps verification states 6_Enha, 11_Enha and 15_Enha. A similar issue occurs across related chromatin state types: base state 10 Biva overlaps a mixture of verification states 12_Biva and 13_Prom_fla. This suggests that irreproducibility may stem from excessive granularity of states. That is, dividing the genome among 16 labels as done by existing SAGA pipelines (section 4.2) requires a resolution in chromatin states that is too fine to achieve robust results. There is a trade-off between reproducibility and the number of states; a two-state annotation is likely to have near-perfect reproducibility, but carries little information (Figure 2 F,G).

**Figure 2.**
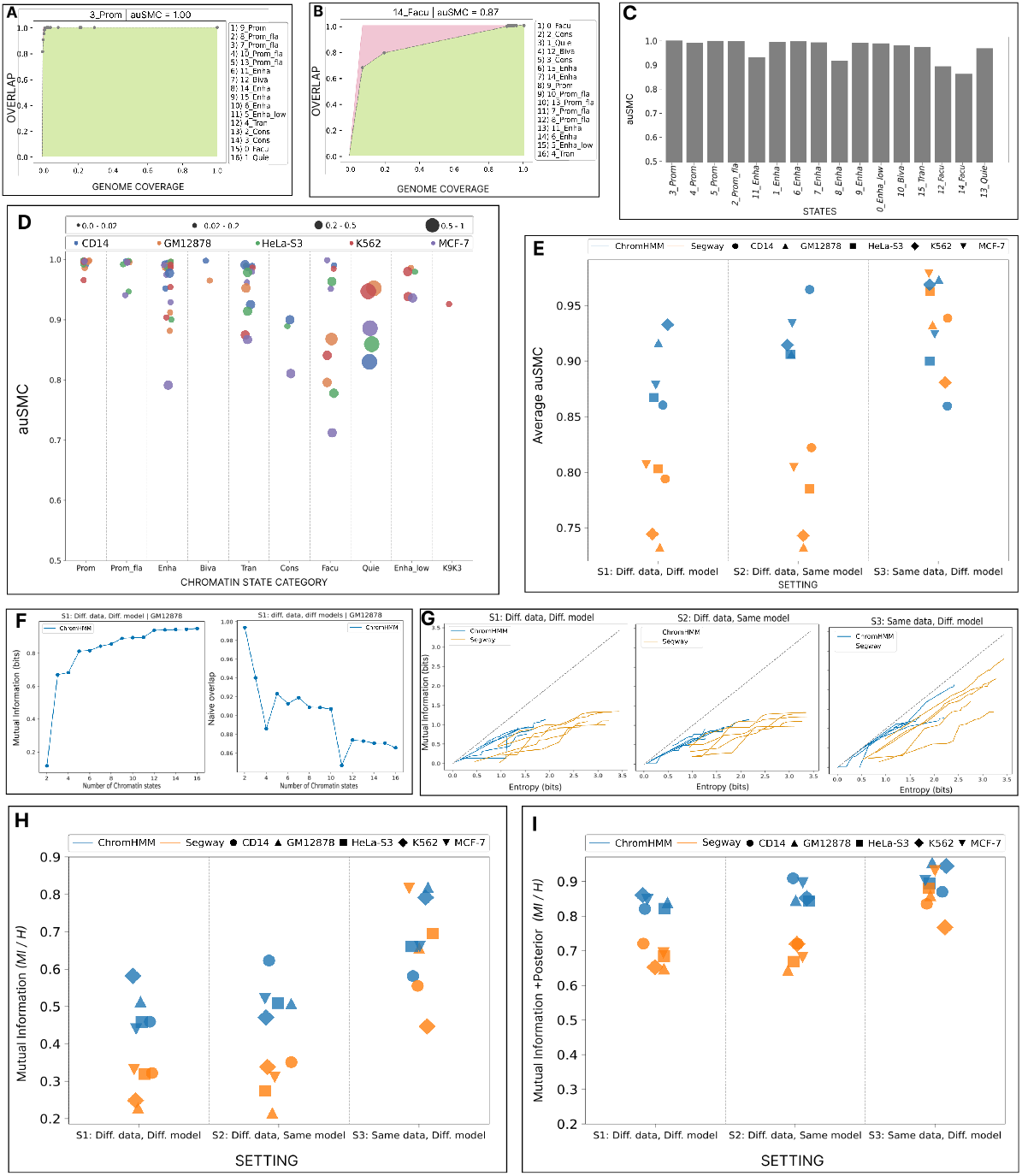
Evaluation of reproducibility as a function of granularity of chromatin state. **(A)** For a given chromatin state in the base annotation *B*_*i*_, we ordered the chromatin states in the verification annotation *V* according to their intersection over union overlap. The leftmost dot indicates the most-overlapped verification chromatin state *V*_*j*_ ; the horizontal axis indicates *V*_*j*_ ’s genomic coverage and vertical axis indicates the fraction of *B*_*i*_ overlapping with *V*_*j*_. The second point corresponds the the union of *V*_*j*_ with the second-most enriched verification chromatin state, and so on for subsequent points. This forms an ROC-like curve, with the green area representing the AUC for the observed overlap versus genomic coverage of each state, and the red area indicating deviation from perfect reproducibility where the first verification chromatin state covers all positions of target chromatin state in base annotation *B*_*i*_. The area under the state merging curve (auSMC) ratio is a numerical representation of chromatin state reproducibility as a function of chromatin state granularity which is calculated by dividing the observed area under the curve by the area under the curve pertaining to the perfect reproducibility case. In other words, larger red area corresponds to lower auSMC. Results are shown for the Promoter (3 Prom) chromatin state in the base annotation obtained from our running example (setting 1, ChromHMM, GM12878) Lists of chromatin state names on the right of A and B represent chromatin states in the verification annotation sorted according to their intersection over union (IoU) overlap with the target chromatin state in the base annotation. **(B)** Same as (A), but for Facultative Heterochromatin (14 Facu) **(C)** auSMC ratio of chromatin states in the base ChromHMM annotation according to S1 (different Data, different models) from GM12878 cell type. **(D)** auSMC ratio of various chromatin states categories identified in the ChromHMM annotation according to S1 (different Data, different models) for five cell types. Each dot represents a chromatin state, with color denoting cell type and size proportional to genome coverage. **(E)** Average auSMC ratio (weighted by the genome coverage) across two SAGA models, five cell types, and three settings. Color denotes the SAGA model and shapes represent cell types. **(F)** Mutual information (left) and naive overlap (right) as a function of the number of chromatin states, for our running example (ChromHMM, GM12878, setting 1). Horizontal axis indicates the number of states; the default 16-label model is on the far right, and each dot to the left represents an annotation after merging two labels in the annotation to its right. Mutual information indicates the number of bits of information about the verification state that is gained by observing the base state (Methods). **(G)** Mutual information between base and verification replicates after merging labels, as a function of the entropy of the base annotation. In a perfectly-reproducible cases, the amount of mutual information would the entropy. Color denotes SAGA model (Segway or ChromHMM). **(H)** Mutual information between base and verification annotations across two SAGA models, five cell types, and three settings as a fraction of the base annotation entropy. **(I)** Same as (H), but evaluating the mutual information when observing both the label and posterior probability (Methods).

**Figure 3.**
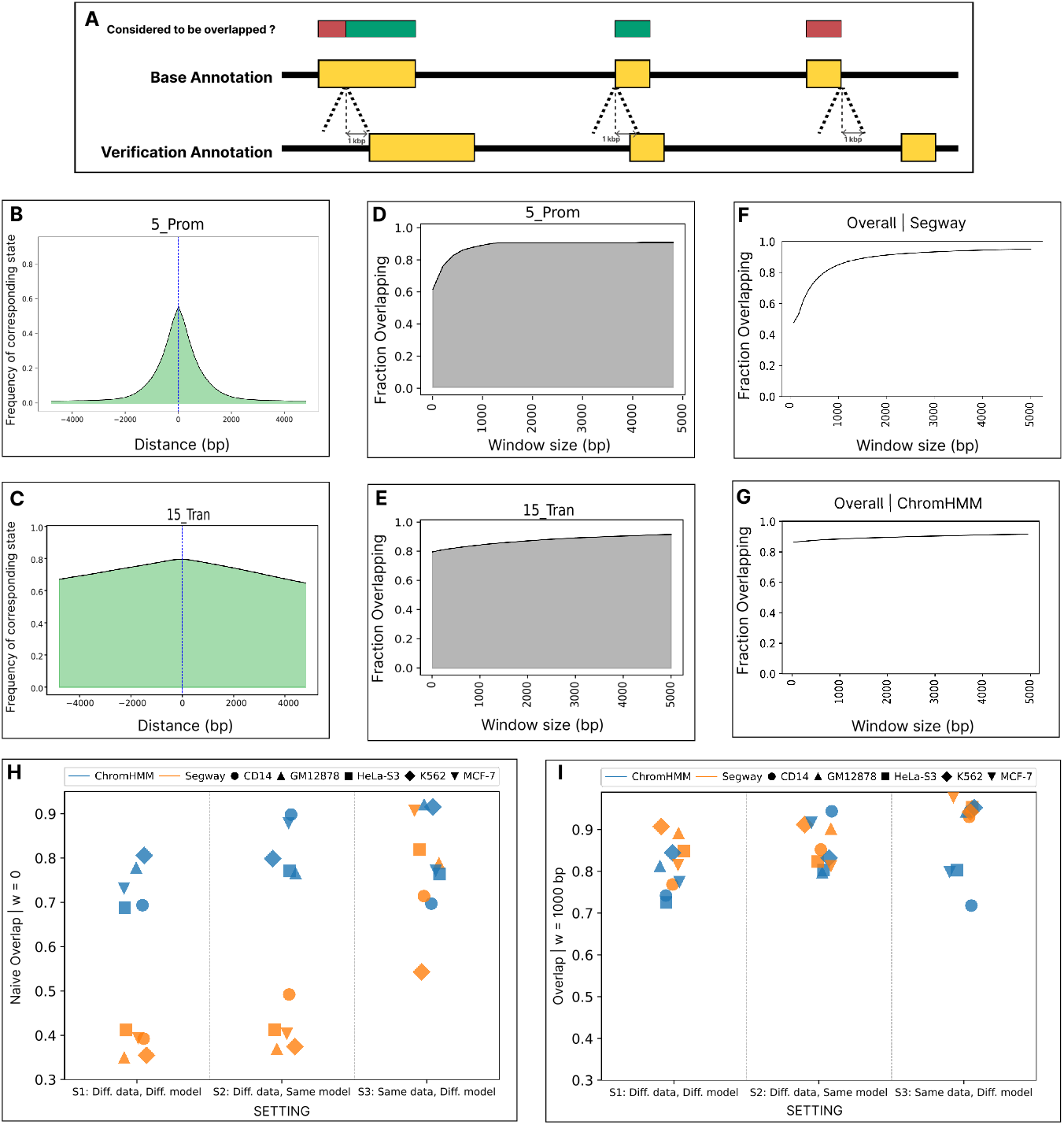
Spatial misalignment of corresponding chromatin states. **(A)** Schematic depicting the types of state misalignment and the influence of the tolerance window *w*. The horizontal axis depicts a genomic window. Yellow rectangles indicate a base state and its corresponding verification state. Red/green rectangles indicate whether the given position in the base annotation would be counted as overlapping the verification replicate. **(B)** Given that a position *g* is annotated as base state 5_Prom, the probability that a nearby position is annotated as the corresponding verification state (13_Prom_fla) as a function of distance from *g*. **(C)** Same as (B), but for 15_Trans. **(D)** Given that a position *g* is annotated with base label 5_Prom, the probability (vertical axis) that the corresponding label occurs within a window *g* ± *w*, as a function of *w* (horizontal axis). **(E)** Same as (D), but for 15_Trans. **(F**,**G)** The overall overlap of the base annotation from Segway and ChromHMM, respectively as a function of *w* (GM12878, setting 1). **(H)** Naive overlap across two SAGA models, five cell types, and three settings. Color denotes the SAGA model and shape represents cell type. **(I)** Same as (H), but allowing for a spatial tolerance window of *w* = 1000.

Thus, we evaluated whether each base annotation state can be recovered using multiple verification annotation states. To accomplish this, for each base state, we ordered verification states according to their intersection over union (IoU) overlap then iteratively merged the top two states in terms of intersection over union (IoU) overlap to create a “super-state” in the verification annotation that eventually covers the entire genome. This process produces an ROC-like (receiver-operating characteristic-like) curve showing the fraction of overlap versus genomic coverage of each state. The area under this state merging curve (auSMC) is a measure of a state’s reproducibility when taking into account such merges, calculated according the ratio of the observed area under the curve (shaded green in Figure 2A, B) to the the area under the curve pertaining to the perfect reproducibility (shaded red). We found that, indeed, base chromatin states can often be recovered using a combination of verification states. For example, although only 80% of base state 3 Prom is overlapped by its corresponding verification state 9 Prom, a further 10% overlaps related label 8 Prom fla, resulting in auSMC*>*0.99 (Figure 2A). Both SAGA models are almost always able to identify Enhancer chromatin states with auSMC *>* 0.8 (Supplementary Section 3.2). However, Promoter chromatin states identified by ChromHMM generally show strong auSMC (Figure 2D), while for Segway they often show a poor auSMC, even in S3 (same Data, different models) (Supplementary Section 3.2) However, some base states are not reproducible even when accounting for multiple verification states. For example, although 68% of base state 14 Facu is overlapped by its corresponding verification state 0 Facu, one must merge both other repressive states (2 Cons and 1 Quies), covering most of the rest of the genome, in order for the merged state to cover more than 90% of 14 Facu, resulting in aucSMC = 0.87. Facultative heterochromatin has the largest variation in terms of auSMC, likely due to differing thresholds dividing Quiescent and Facultative Heterochromatin states (Supplementary Section 3.3.3). As with naive overlap, settings 1 and 2 (different data, different models and different data, same model, respectively) have similar performance but setting 3 (same data, different model) performs much better, especially for Segway, indicating that most differences are cause by differences in the input data (Figure 2 E).

Information theory provides a more general way of evaluating reproducibility while accounting for the lack of one-to-one correspondence (Section 4.5). We found that, for our running example, the base annotation conveys 0.95 bits of mutual information about the verification annotation, out of 1.56 total bits of entropy replicate (Fig 2 H). In general, for both ChromHMM and Segway annotations, the base annotation usually conveys about 1 bit of information about the verification annotation (Figs 2 G,H).

These results suggest that reducing the number of states may give more robust annotations. Running SAGA with all possible numbers of states would be computationally infeasible and analysis likely would suffer from noise due to model training, so instead we simulated annotations with fewer states by merging states in the base and verification replicates, respectively. Thus, we created annotations with fewer states by merging states in the base and verification annotation, respectively. We iteratively merged pairs of state in each annotation base and validation until we were left with two states in each annotation, where at each iteration we merged the pairs resulting in the smallest loss of mutual information (Section 4.10, Figure 2 F,G,H)). As merging low-coverage labels has a smaller impact than merging high-coverage labels, throughout our analysis below we use the bits of entropy to estimate the complexity of an annotation instead of the raw number of labels (Figure 2 G). We found that SAGA annotations with fewer labels have greatly improved reproducibility. In our running example, moving from a 16-state to a 10-state annotation improves naive overlap from 87% to 91% while only reducing the entropy of the annotation from 1.56 to 1.37 bits (Figure 2 F).

### 2.4 Spatial misalignment of corresponding chromatin states leads to irreproducibility

We found that a substantial fraction of mismatch between replicates results from spatial misalignment of segment boundaries. This may occur when two segments with corresponding chromatin states from base and verification annotations partially overlap, but their borders do not fully align, or when two non-overlapping segments with corresponding states occur in close proximity (Fig. 3A). Such imprecision in segment boundaries may be unimportant to practitioners since it may not meaningfully hinder downstream applications, such as in identifying putative regulatory elements or localizing disease-associated variation.

Therefore, to evaluate the reproducibility of annotations, we investigated the overlap between analogous chromatin states while considering a window of size *w*=1000 base-pairs upstream and downstream of any given genomic positions. In other word, in this refined definition of overlap, each annotation at position *g* in base annotation is considered overlapped if its corresponding state is observed within the window of *g ± w* in the verification annotation.

We found that a large fraction of irreproducibility in SAGA annotations can be be attributed to spatial misalignment (Figure 3). Corresponding states frequently occur upstream and downstream of a target position (Figure 3 B, C). When measuring overlap while considering a window around each position, we observe that for the majority of chromatin state types, a window of size *w*=1000 bp significantly increases the overlap by capturing most of the misalignments, especially for chromatin states with short segments. For instance, in our running example, *∼*0.6 of regions labeled as 5 Prom exactly overlap (*w* = 0) their corresponding state in verification annotation (Figure 3 D). However, by slightly increasing the window size, the fraction of overlapping regions increases to *∼*0.9. This implies that for roughly *∼*0.3 of positions that are labelled as 5 Prom in base annotation, we can find a corresponding state in verification annotation within close proximity (1kbp). However, states such as 15 Tran are less affected by spatial misalignment.

Segway annotations are particularly susceptible to misalignment. Allowing for misalignment up to *w*=1000 increases overlap in Segway annotations from 47% to 85% (Figure 3F). In contrast, we do not observe a similar pattern for ChromHMM (Figure 3G). This pattern occurs because Segway uses signal values rather than binarized data and thus is sensitive to variation in the scale of input signals across replicates. This can be attributed to hyper-segmentation in Segway annotations due to its sensitivity to variation in input signals [3, 15]. Segway also tends to hyper-segment the genome into small segments that are inherently more prone to misalignment than longer segments.

Notably, without accounting for misalignments, in settings S1 (different data, different models) and S2 (different data, same model), we can observe a distinct gap between ChromHMM and Segway in terms of overall overlap (Figure 3H). However, S3 (same data, different models) essentially removes this gap and results in overall overlap values that are roughly in the same range for both ChromHMM and Segway. In S3 (same data, different models), we do not observe severe misalignment issues for Segway, which confirms Segway’s sensitivity to noise in the input data. Moreover, ChromHMM’s overall overlap is significantly less dependent on windowsize, suggesting less susceptibility to misalignment due to data binarization and removal of fine details in the input data. After accounting for misalignment, Segway annotations significantly improve in terms of reproducibility and slightly outperform ChromHMM annotations(Fig. 3 I).

### 2.5 Posterior probability indicates robustness of annotations

SAGA methods generate posterior probabilities that indicate the model’s confidence about the state assignment at every genomic position. We hypothesized that these posterior probabilities indicate reliability, such that confidently-annotated positions are robust. Unfortunately, these probability values are often vastly too confident; most positions receive *>* 99% posterior [9].

We found that, although rates of reproducibility are much lower than model posterior probabilities, there is a strong increasing relationship between posterior probability and reproducibility. Specifically, for each chromatin state in the base annotation, we evaluated the frequency with which it overlaps its corresponding state in verification annotation as a function of the model’s posterior probability, using an isotonic regression (Section 4.8). For Segway, doing this analysis required modifying the underlying model to artificially weaken model predictions to avoid posteriors being rounded up to exactly 1.0 due to floating point under/overflow (Supplementary Section 1).

The results showed a positive trend between posterior probability and reproducibility, indicating that higher posterior probabilities are associated with higher levels of reproducibility (Figure 4A). This means that despite being over-confident, posterior probability of SAGA methods contain information about their reproducibility. Posterior probabilities also contains biologically-relevant information about gene expression and transcription start sites, further confirming their usefulness. We found that there is a strong correlation between posterior probability of the Transcribed state within a gene body and expression of that gene (Figure 4B). That is, as the posterior of transcribed chromatin state increases, the average expression as measured by RNA-seq increases as well. Conversely, the posterior of the Quiescent state is negatively correlated (Figure 4C). Notably, even for positions that did not ultimately receive a label of Quiescent, a small but non-zero posterior probability of Quiescent at the gene body is associated with low expression.

**Figure 4.**
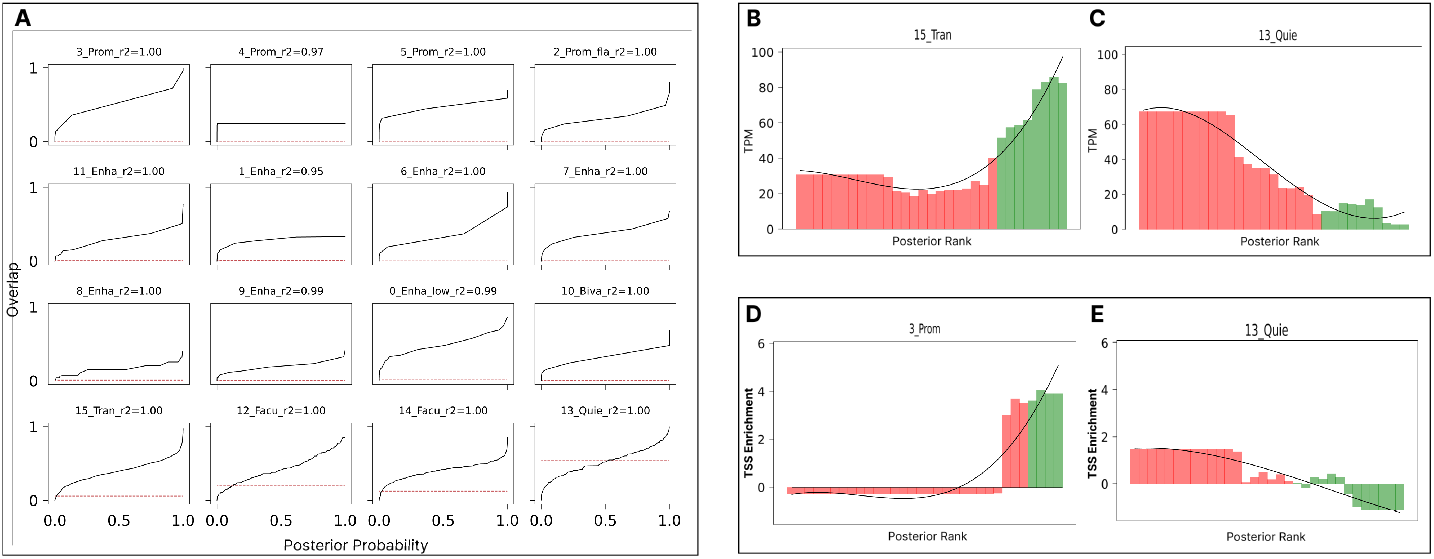
Posterior probability is associated with reproducibility. **(A)** The frequency of overlap with a corresponding state as a function of posterior probability for all states in the base annotation in running example (ChromHMM, GM12878, setting 1). In each sub-panel of **(A)**, the horizontal axis corresponds to the posterior probability and the vertical axis corresponds to the observed overlap. The red dotted horizontal line in each sub-panel represents the genome coverage of the corresponding state in the verification state, indicating the overlap expected from random overlap. The intersection of the calibration curve and the red dotted-line corresponds to the posterior threshold from which any larger probability would exceed the probability of random overlap. *R*^2^ score represents how well the isotonic regression curve fits the data. **(B)** shows the mean expression level (TPM) as a function of posterior of 15_Tran. The horizontal axis represents rank of posterior probability, while the vertical axis shows TPM. **(C)** Same as (B) but for 13_Quie. Similarly, **(D)** shows the enrichment around TSS regions as a function of posterior of a 3_Prom. The horizontal axis shows the rank of posterior probability and the vertical axis shows enrichment around TSS. **(E)** Same as (D) but for 13_Quie. confirming that the posterior contains biologically-relevant information about transcription start sites. In **(B-E)**, green bars represent genomic loci for which the maximum-a-posterior is the target chromatin state in the base annotation and red bars correspond to the loci where the maximum-a-posterior is another chromatin state.

Similarly to the result on gene bodies, posterior of Promoter states is positively correlated with occurrence of annotated transcription start sites (TSSs), and the converse is true of Quiescent states (Figure 4D,E).

We found that the posterior of the state assignment to the base annotation conveys a great deal of information about what state is assigned to the verification annotation (Methods). As mentioned above, knowing only the identity of the state assigned to the base annotation removes 51% of the uncertainty in the verification state, as measured by the mutual information/entropy ratio (Figure 2H). However, knowing both the state and its posterior probability increases the mutual information to 85%, suggesting that the posterior probability strongly indicates whether the annotation of a given genomic bin is reliable. This increase is even more dramatic for Segway, increasing the fraction of mutual information from 0.20-0.35 to 0.64-0.72 (Figure 2I).

### 2.6 SAGAconf yields robust annotations

As described above, even when allowing for variability in granularity of state definitions and in spatial positioning, the reproducibility of SAGA annotations falls below the 90-95% threshold sought in most applications. Fortunately, as describe in the previous section, the posterior probability output by the SAGA probabilistic model indicates the reliability of the annotation of each genomic position.

In order to produce reliable chromatin state annotations, we propose SAGAconf, a method for producing robust chromatin state annotations. SAGAconf infers the probability that the annotation to a given genomic position will be reproduced in a replicate experiment (within a spatial tolerance of *w* = 1000 bp), according to the SAGA model’s output state and posterior probability, which we term the r-value (Section 4.12). SAGAconf outputs the annotation to a subset of the genome passing a user-defined r-value threshold (usually 90% or 95%). Thus SAGAconf is analogous to the IDR pipeline used for ChIP-seq peak calling and related tasks [18]. SAGAconf is independent of SAGA methodology, as it takes as input only a SAGA annotation and its posterior probability matrix.

We found that *r*-values are distributed differently across chromatin states (Fig 5 A). In our running example, nearly all positions labeled with 3 Prom achieve *r >* 0.9, meaning that this state is reproduced independent of model posterior. Conversely, all positions with state 4 Prom have *r <* 0.9, indicating that this state can never be confidently annotated. However, most states show a range of *r*-values, indicating that the reliability depends on the posterior; for example, 52% of state 0 Enha low have sufficiently-high posterior to pass the *r >* 0.9 threshold. Thus applying SAGAconf is critical to knowing whether the annotation to any given locus can be relied upon.

**Figure 5.**
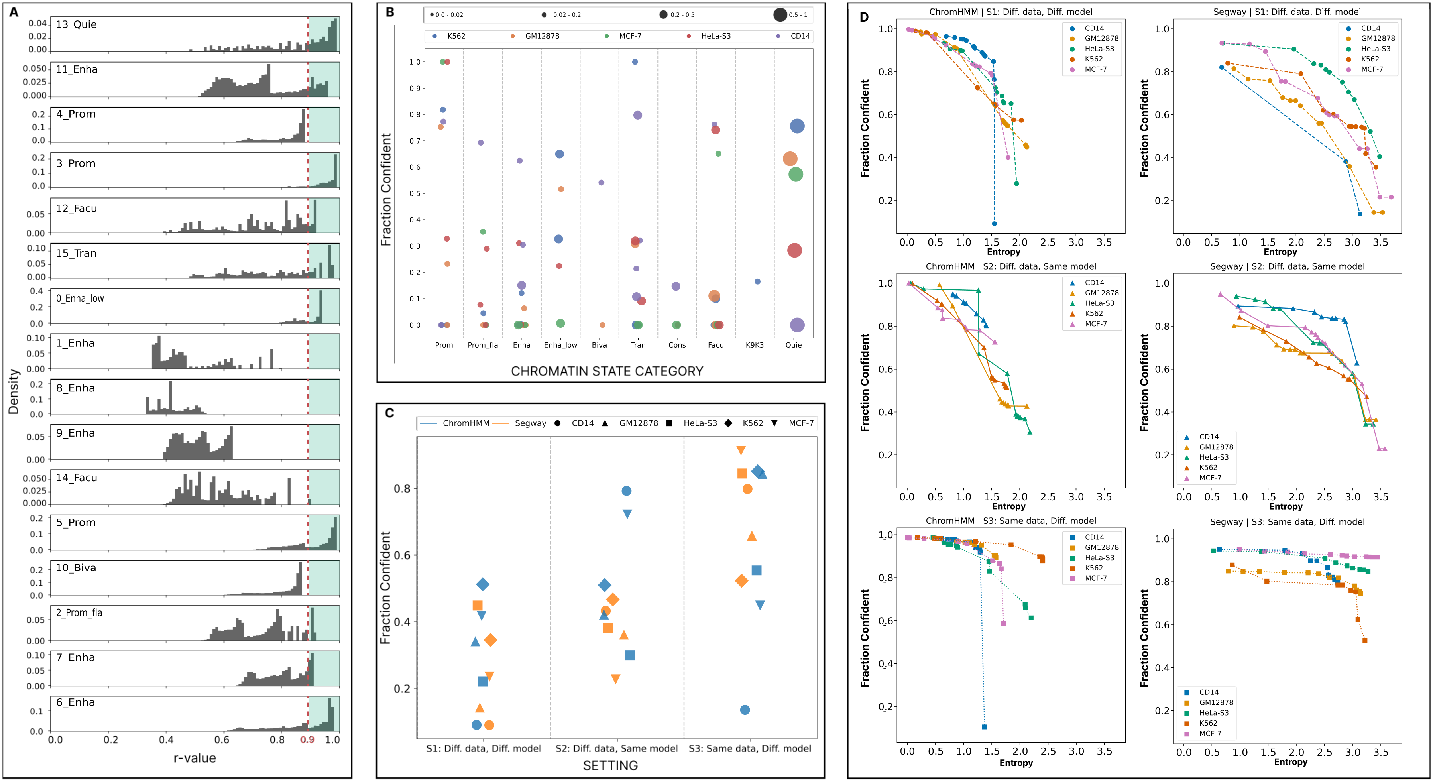
SAGAconf identifies confident state annotations. SAGAconf integrates sources of reproducibility information such as granularity, alignment, and posterior to derive an *r*-value for each genomic position. The r-value estimates the probability of reproducing the annotation at a specific location. **(A)** Density histogram of *r*-values for each chromatin state in the running example (setting 1, ChromHMM, GM12878). A threshold of *α* = 0.9 is applied to label positions with *r*-value *≥α* as reproducible and irreproducible otherwise. The horizontal axis represents *r*-value, the vertical axis represents density, the red dotted line represents the threshold, and the green-shaded area show confident annotations. **(B)** Fraction of chromatin states called confident by SAGAconf for ChromHMM annotations according to S1 (different Data, different models) in five cell types. Each dot represents a chromatin state, with color denoting cell type and size proportional to genome coverage. **(C)** Fraction of genome called confident across two SAGA models, five cell types, and three settings. Color denotes SAGA model and shapes represent cell types. **(E)** We measured the fraction of the genome called as confident by SAGAconf during the process of post-clustering; that is, we measure the fraction of confident positions in the genome as a function of the base annotation’s entropy as we merged similar chromatin states in the base annotation. The horizontal axis represents base annotation entropy, the vertical axis represents the fraction of genome identified as confident, sub-panels correspond to different SAGA models and settings, and colors represent cell types.

Overall, 10%-50% of positions are annotated confidently according to SAGAconf (Figure 5C). Most enhancer states from ChromHMM are irreproducible (Figure 5B), while Segway identifies more-reproducible enhancers (Supplementary Section 6). After applying SAGAconf, Segway and ChromHMM both show roughly similar reproducibility in terms of fraction of the genome which is identified as confident (Figure 5 C).

SAGA models with fewer states can confidently annotate a much larger fraction of the genome. In our running example, the default 16-label annotation has 2.1 bits of entropy and confidently annotates 43% of the genome, but merging down to an annotation with 1.7 bits (10 labels) yields 55% of the being being confidently annotated. This difference is more extreme for Segway. Although the default 16-label annotation can confidently annotate only a small fraction (*<* 10%), a 1.7 bit annotation confidently annotates 64% of the genome.

## 3 Discussion

Chromatin state annotations are essential for various downstream tasks, including identifying regulatory regions and cell type-specific activity patterns, interpreting disease-association studies, studying gene regulation, and analyzing cellular differentiation [1, 2, 14, 10, 9]. Therefore, it is paramount to ensure their reliability, else all subsequent analysis may be inaccurate. Yet, despite the fact that statistical guarantees (such as p-values, q-values or IDR) are used ubiquitously in genomics and in science in general, no such statistical guarantee previously existed for SAGA. Here, we provide a comprehensive evaluation of genome annotations in terms of their reproducibility and confidence. Using replicated data, we delineate different sources of irreproducibility stemming from the data and the SAGA models. We found that SAGA chromatin state annotations are frequently irreplicable, meaning that they often disagree when run on two replicated data sets. A substantial fraction of this disagreement remains after accounting for mismatch in chromatin states across models and for spatial misalignment between segments. This finding suggests that there is significant risk to using SAGA annotations without any filtering.

To mitigate this risk, we introduce a framework SAGAconf that identifies a confident subset of the genome that is annotated reliably. SAGAconf does so by leveraging the posterior probability of the underlying probabilistic graphical model used by the SAGA method, which we demonstrated to be informative of reproducibility. We showed that SAGAconf correctly distinguishes reliable versus unreliable SAGA annotations within the genome. Thus downstream applications of SAGA annotations would be improved by applying SAGAconf to filter out genomic positions with unreliable labels. This filtering step is analogous to the use of irreproducible discovery rate (IDR) analysis for ChIP-seq peak calling.

This study tackles a repeatedly-encountered problem in the field that is the lack of comprehensive and principled approaches for evaluation of SAGA genome annotations. Conventionally, SAGA genome annotations are evaluated using qualitative and quantitative methods [9]. Qualitative methods involve examining various statistics of an annotation to determine whether it reflects expected features of genome biology [27]. However, currently, there are no generally agreed upon statistics that hold for all high-quality annotations. Quantitative methods on the other hand, involve prediction problems such as predicting RNA-seq expression given only the annotation label at the gene’s promoter [28] [10], [7]. But, such prediction tasks are useful for the purpose of comparing different annotations but do not serve as a realistic evaluation of the annotations themselves. This study introduces SAGAconf as a comprehensive and principled alternative for this task that addresses various shortcomings of previous evaluation approaches.

The findings of this study suggest that reproducibility of chromatin states depends greatly on the quality of data; that is, estimations of confidence are affected by the replicate with inferior data quality. In addition to obtaining better replicated data, and effective pre-processing methods [29, 30, 19], future endeavors can tackle this problem by designing SAGA models that are more robust to data quality.

Future studies should enhance our understanding of genome annotation confidence by evaluating the effect on reproducibility of the choice of SAGA methods (e.g. Segway [3], ChromHMM [4], IDEAS [12] and others), model hyper-parameters, resolution (e.g., by using coarse resolution to annotate domains), and data types (e.g., conservation via ConsHMM) [31]. Additionally, factors such as data quality and the availability of epigenomic assays should be considered. These measurements can be potentially valuable for inferring developmental and lineage-specific changes in the epigenome.

Currently, SAGAconf treats the base and verification replicates separately. Combining these replicates to obtain a unified representation of reproducibility across experiments is a potential area for future research. A naive approach to combining replicates would be to run SAGAconf twice, with each of the two possible assignments to base and verification, then output the intersection of the confident regions.

Many epigenomic assays on public databases such as [1] are not performed on replicated experiments, so a version of SAGAconf that does not require replicated data would be valuable. One naive option would to use data from a similar cell type the produce a verification replicate. Such a confidence estimate will be conservative, meaning that a smaller fraction of the genome will be deemed reproducible by SAGAconf. There is a need for future work to develop a model that can generalize statistics from a replicated cell types to unreplicated cell types.

## 4. Methods

### 4.1 Data collection

We retrieved sets of replicated histone modification ChIP-seq data for five cell types from the ENCODE DCC [1]^[1]^. These cell types are all among ENCODE’s tier-1 and tier-2 cell types and have the most number of ChIP-seq assays with isogenic replicate data (Supplementary Section 1). Isogenic replication, also referred to as biological replication, is a process in which two biosamples are derived from the same human donor or model organism but treated separately ^[2]^.

Consequently, we collected pairs of replicated histone modification ChIP-seq data for the following cell types.

1. GM12878 with 11 histone modification assays
2. K562 with 11 histone modification assays
3. HeLa-S3 with 11 histone modification assays
4. CD14-positive-monocyte with 11 histone modification assays
5. MCF-7 with 13 histone modification assays

For each cell type, we assembled two data sets each consisting of all of the assays belonging to one replicate. These datasets consist of a feature vector for each genomic position in which each element corresponds to a particular histone modification measurement. With *d* histone modification assays, the dataset is a R*G×d* matrix, where *G* denotes genomic positions. For instance, for cell type GM12878, we have two replicated datasets, and in each dataset, we have an 11-dimensional feature vector per genomic position. These feature vectors are considered by SAGA models as observed events *X*_*g*_, and are used as training data. All of the ChIP-seq data are aligned with hg38 (GRCh38) reference human genome.

### 4.2 Model training and annotation

Using the collected data, we trained ChromHMM and Segway models with a fixed set of hyper-parameters and obtained chromatin state annotations from them. Specifically, for both methods, to specify the number of chromatin states, we used the formula 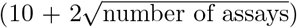 as suggested by [10] to scale with the amount of available data (11 histone modification assays for GM12878, K562, HeLa-S3, CD14-positive-monocyte, and 13 assays for MCF-7). For specifying hyper-parameter settings of both ChromHMM and Segway, we followed standard practice as previously performed by [14, 3, 32, 10, 4]. Details of hyper-parameter settings that were used for model training and annotation are explained in the Supplementary Section.

As a post-processing step, to enable comparison between two replicates, we divided the genome into resolution-sized positions and assigned the posterior *P* (*Q*_*g*_ = *q*|*X*) and annotation results to each genomic position. This additional parsing step provides us with a unified format which is readily comparable, independent of datasets, parameters, or even the SAGA algorithm that generated the annotations. Parsed annotation results are matrices of size *G × K* where *K* denotes the number of chromatin states, and *G* corresponds to the number of genomic positions; that is, the whole genome size divided by `--resolution` and each element in the matrix corresponds to the posterior probability of each state at each position.

### 4.3 Biological state interpretation

As the raw annotation results of SAGA algorithms are states named with the integer ID of their clusters, we need an additional interpretation step in which cluster IDs are translated to obtain human-readable results into functional biological roles (also known as “mnemonics”). To avoid bias from manual interpretation, we used an automated interpretation procedure introduced in [10]. Specifically, we used a pre-trained random forest classifier which, using the enrichment of states around conserved regions and enrichment of different histone modification marks for each state, assigns a predicted biological interpretation to each state. This classifier is trained on Segway annotations and might not be as accurate on ChromHMM annotations. These interpretation terms are solely for the interpretability of results, and every step of the reproducibility evaluation pipeline is independent of the interpretation terms [33].

### 4.4 Pairwise overlap of chromatin states

SAGA methods output integer state IDs for each identified chromatin state. Because they are unsupervised, the state IDs are not consistent across two different models. Therefore, to enable comparison among chromatin states of the two annotations, the first step is to identify states that correspond to related genomic functions across the two annotations by measuring the pairwise overlap between them.

Let *k* be a chromatin state in the base annotation *B* and *l* be a chromatin state in verification annotation *V*. Their joint frequency of overlap 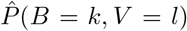 is defined as:

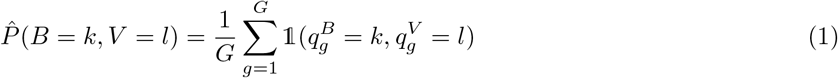

Where *g* corresponds to genomic positions and 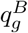 and 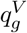 denote assigned states to the genomic position *g* in base and verification annotations, respectively. Similarly,

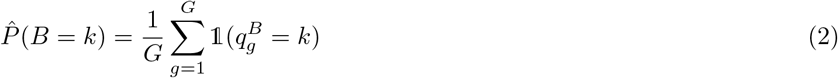

is the marginal frequency (i.e. genomic coverage) of state *k* in the base annotation *B*.

The joint distribution of overlap 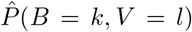 is greatly influenced by the states’ genome coverage. For example, pairs of Quiescent states that cover most of the genome, will have a very high probability while highly corresponding promoter pairs might get small overlap probabilities. To identify correspondence of states regardless of their genome coverage, we use intersection over union (IoU), also known as Jaccard’s similarity coefficient, which is formulated as follows.

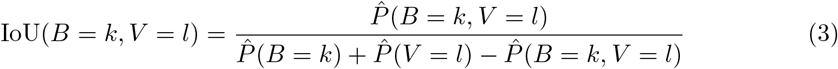

### 4.5 Mutual information between annotations

The mutual information of base and verification annotations *I*(*B*; *V*) can quantify the shared information of two annotations and it is calculated as follows:

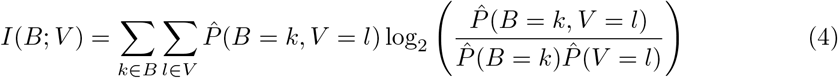

Mutual information can have values ranging from 0 for total independence and +*∞*. However, MI *I*(*B*; *V*) is always upper bounded by both entropies of the base annotation *H*(*B*) and the verification annotation *H*(*V*) which quantify the total amount information that can be contained within these annotations. Entropy of each set of annotation is calculated as follows:

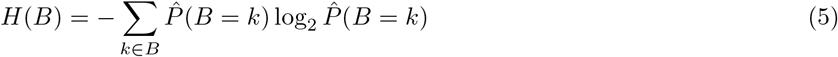

Therefore, we normalize the mutual information into values that reflect the fraction of information in the base annotation that is shared with the verification annotation.

The posterior of the base replicate provides information into the replicability of the annotation. We compute the information as follows. We make 10 equally distanced bins based on the posterior values of the base annotation chromatin states. Let *I* be a bin index, where 1 *≤ i ≤* 10, the mutual information between the base and verification annotations *I*(*B*; *V*) with respect to posterior probabilities of the base annotation is calculated as follows:

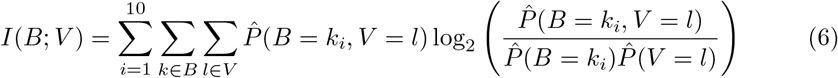

### 4.6 Overlap and spatial alignment

We hypothesized that due to a variety of noises, corresponding chromatin states might not locate at the exact same position across the two annotations. To investigate the impact of misalignment of corresponding segments, we relaxed the criteria for considering an annotation as overlapped by allowing a window of size *w* upstream and downstream of any given position *g* to look for corresponding states. Therefore, we consider proximal positions in the overlap calculations:

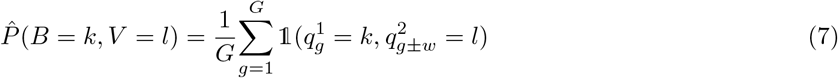

Where 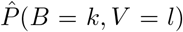 is the updated probability of overlap which considers a window of size *w* around segments of the verification annotation.

### 4.7 Granularity of genomic functions

In a perfect case of reproducibility, one would expect that for each state *k* in base annotation, one state should exist in the verification annotation that exactly covers the same genomic positions. However, in practice, positions with state *k* in *B* are distributed over two or more states in *V* and vice versa.

To understand the effect of genomic functional granularity (i.e. the number of chromatin states) on reproducibility, we measure the overlap while iteratively verification state. To that end, for a given target state of base annotation, we sorted all of the states of verification annotation according to their intersection over union (IoU) of overlap. Based on this order, we start merging the states in *V* iteratively until all of the genome is covered with one “super-state” in *V*. We evaluate what fraction of the genome needs to be covered by *V* states in order to cover most of the positions of the target state in *B*. We term this the “state merging curve” (SMC). The area under the SMC in case of perfect reproducibility, is when = the first *V* state covers all of the positions of target state in *B*. We define the ratio between these two curves as the area under the SMC (auSMC), which is a simple numerical representation of the state’s reproducibility as a function of its chromatin state granularity.

### 4.8 Calibration of posterior probability

The posterior probabilities produced by SAGA tend to be overly confident, with the majority of positions receiving a probability *>*99%, which does not accurately reflect the reliability of the annotations. However, we investigated if there is any correlation between reproducibility and posterior probability. By examining the relationship between posterior probability and reproducibility, as measured by pairwise overlap, we can improve confidence estimates by creating a calibration curve. This allows us to transform raw posteriors into more accurate and robust confidence scores that better represent the actual likelihood of reproducibility for a given annotation.

Initially, we establish pairs of corresponding chromatin states. For each state *k* in *B*, the state with the greatest IoU overlap score in *V* is considered as the matching state. To carry out calibration, we computed the ratio of overlap as a function of the model’s posterior. Specifically, let 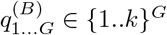 and 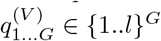 represent vectors of base and verification annotations respectively. In base annotation, for each state *k*, we first arrange the vector of posterior probabilities of all genomic positions 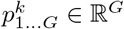 based on the posterior value. Then, we divide the sorted array into b sub-arrays (bins) of equal size such that *b*_*i*_ *⊂ {*1 … *G}*. For each bin, we compute the fraction of the target state in the base annotation that overlap with its corresponding state in the verification annotation. It is expected that for first bins with lower posterior value, the overlap is lower than that of high posterior bins. By performing the binning step, we can quantitatively compare two variables with different natures, namely the overlap and posterior probability.

We assume that the pairwise agreement and posterior probability of corresponding chromatin states are not negatively correlated. Thus, assuming a monotonic and non-decreasing trend, we fit an isotonic regression model to create a calibration curve [34, 35, 36]. Isotonic regression is a non-negative piece-wise regression model in which we aim to learn a curve ŷ to solve a problem formulated as follows:

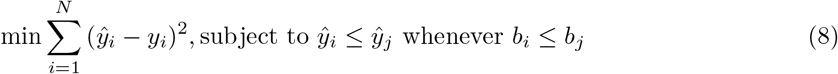

To further explain, the model attempts to fit a curve ŷ to the sequence of “posterior vs. overlap ratio” bins *b* such that for all *b*_*i*_ *≤ b*_*j*_, the curve ŷ_*i*_ *≤* ŷ_*j*_ will be produced, resulting in a non-negative trend. We selected the isotonic regression model for this calibration task because, unlike linear regressions that impose linearity, these models are not limited by any functional form and can fit any form in the observed data as long as it is monotonically increasing. The Pool-Adjacent-Violators Algorithm (PAVA) is commonly used to fit the isotonic regression model [34, 35, 36]. We utilized isotonic regression from Python’s Scikit-learn package [37].

### 4.9 Validation against known phenomena

In contrast to supervised learning problems, where the predictions of models can be validated against a labeled test set, chromatin states lack a gold standard. One common method to validate annotations is to analyze the enrichment of promoter-like states around conserved positions such as transcription start sites (TSS) from reference annotations [38, 10, 24, 12]. Another method is to check if the transcribed regions predicted by SAGA models overlap with experimentally-validated expressed regions obtained from RNA-seq data [28].

We used this approach to evaluate whether the posterior probability of annotations can confidently predict known regulatory regions. Firstly, similar to calibration of posterior probability, for each state *k*, we rank the posterior probabilities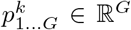 in ascending order. Then, we split the sorted array into *b* equal size sub-arrays (bins) such that *b*_*i*_ *⊂ {*1 … *G}*. For each bin of posterior probability, we calculate the enrichment of regions with the posterior of state *k* in this bin range around transcription start sites. Enrichment is calculated as 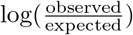 In this case, observed is the number of regions with the posterior of state *k* in a given bin range around TSS while expected is the number of positions with the posterior of state *k* in that bin range across the whole genome. Similarly, for each bin, we investigate the mean RNA-seq expression level (TPM or transcripts per million) of genomic positions within that posterior range [39].

### 4.10 Merging chromatin state to produce lower-granularity annotations

Determining the optimal number of chromatin states to specify as a hyper-parameter to the SAGA model is not straightforward. However, it is evident that increasing the number of chromatin states (i.e., increasing granularity) leads to decreased reproducibility, as it becomes more difficult for the model to distinguish between different states. In other words, more granularity in the annotations leads to higher entropy which naturally leads to irreproducibility. This presents a trade-off between the granularity of chromatin states learned from the data and the reproducibility of those annotations.

To investigate this trade-off, we iteratively merge pairs of the most similar chromatin states in both the base and validation annotations. To do this, we need to define a metric *S* that measures the pairwise similarity of different chromatin states within one annotation based on their overlap behavior with the other annotation. We measure this pairwise similarity as follows:

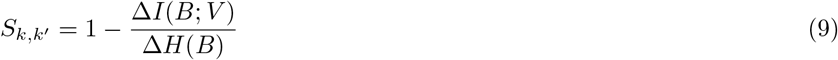

Where ∆*I*(*B*; *V*) and ∆*H*(*B*) represent the change in mutual information and entropy of base annotation, respectively, resulting from merging states *k* and *k*^*′*^ in the base annotation annotation. The verification annotation goes through the exact same process at each iteration.

After calculating metric *S* for both base and validation annotation, at each iteration we merge two pairs with maximum *S* as the most similar pairs for which merging results in the least change in mutual information and the greatest change in entropy. By merging any two chromatin states, both entropy and mutual information decrease however, merging similar chromatin states (with similar pattern of overlap) should ideally result in minimum change in mutual information. This process is repeated until all pairs of chromatin states have been considered for merging. By analyzing how the mutual information and entropy change as pairs of chromatin states are merged, we can determine the optimal number of chromatin states that balances granularity and reproducibility.

### 4.11 Delineating different sources of variability

To obtain a comprehensive insight into how the variability of models and data can affect reproducibility, we assessed the reproducibility of annotations in several settings and different levels of technical and biological variability. Therefore, we selected pairs of experiments in the three following scenarios to evaluate their reproducibility.

**Setting 1 (Different data, Different model)**: We trained two separate SAGA models using the collected data from each isogenic replicate. Two trained models were then used to annotate the genome. Finally, the annotations were compared to uncover the reproducibility from various sources, including the data, the model training process, parameter initialization, etc.

**Setting 2 (Different data, Same model)**: We concatenated data from both isogenic replicates to form a single extended dataset which we used to train a single model. The model provided separate annotations for each of the replicates. In contrast to setting 1, concatenated runs remove elements of variability in training and state matching and can uncover the irreproducible elements that are only attributed to the data from replicated experiments.

**Setting 3 (Same data, Different model)**: We also used the same dataset (from replicate 1) to separately train two different models while only changing the random seed used for parameter initialization. This setting uncovers the irreproducibility that is to be attributed to the initialization and training of models.

### 4.12 SAGAconf

Our analysis suggest that there are various aspects to the problem of reproducibility of SAGA annotations including the granularity of chromatin states, proximity and spatial alignment, and the information embedded within models’ posterior probability. Therefore, we designed SAGAconf, an integrative approach that combines these sources of information to derive an reproducibility score (r-value) at each genomic position which can then be used to filter out irreproducible state assignments and thus obtain robust chromatin state annotations.

The SAGAconf reproducibility assessment pipeline starts by defining corresponding states across the two annotations. We compute a IoU overlap matrix while considering a window of size *w* upstream and downstream of each position according to equations 3 and 7. Then, by setting a threshold *c* on the IoU of overlap, for each chromatin state *k* in the base annotation, its corresponding states in verification annotation should either have a IoU *≥ c* or have the maximum IoU with *k* among all verification annotation states. Note SAGAconf allows for more than one chromatin states in verification annotation to count as corresponding as long as they have IoU *≥ c* which can mitigate the issues associated with chromatin state granularity. Then, we calibrate posterior values into reproducibility score according to section 4.8. Here, at each bin *b*_*i*_ *⊂ {*1 … *G}*, we calculate the ratio of genomic position in base annotation within *b*_*i*_ that have one of the corresponding states in a window of size *w* around that position in verification annotation. Using the isogenic regression obtained from last step, we get a reproducibility score for each position in the genome. The resulting reproducibility score *r*-value ranges from 0 to 1 and it represents the probability of that position or its proximity being labeled with a related genomic function in the other annotation.

Lastly, using a hard threshold *α* on the *r*-value, we assign a Boolean label of “reproduced” for *r ≥ α* or “not-reproduced” *r < α* to every position in the genome. Using these Boolean labels of reproducibility, we can robustly identify a confident and reliable subset from genome annotation while removing the irreproducible predictions.

## Abbreviations

SAGA: Segmentation and genome annotation.
IDR: Irreproducible discovery rate.
IoU: Intersection over union.
auSMC: Area under the state merging curve.
TSS: Transcription start site.
TPM: Transcript per milion.
PAVA: Pool-Adjacent-Violators Algorithm.

## Availability of data and materials

The code used to generate the results presented in this paper is available in our GitHub repository https://github.com/mehdiforoozandeh/SAGAconf. Extended results and detailed information about the datasets used in this study can be found in the supplementary material.

## Footnotes

[1] http://encodeproject.org/

[2] https://www.encodeproject.org/data-standards/terms/

## References

1. Consortium, E.P., et al.: An integrated encyclopedia of dna elements in the human genome. Nature 489(7414), 57 (2012)

2. Kundaje, A., Meuleman, W., Ernst, J., Bilenky, M., Yen, A., Heravi-Moussavi, A., Kheradpour, P., Zhang, Z., Wang, J., Ziller, M.J., et al.: Integrative analysis of 111 reference human epigenomes. Nature 518(7539), 317–330 (2015)

3. Hoffman, M.M., Buske, O.J., Wang, J., Weng, Z., Bilmes, J.A., Noble, W.S.: Unsupervised pattern discovery in human chromatin structure through genomic segmentation. Nature methods 9(5), 473–476 (2012)

4. Ernst, J., Kellis, M.: Chromhmm: automating chromatin-state discovery and characterization. Nature methods 9(3), 215–216 (2012)

5. Day, N., Hemmaplardh, A., Thurman, R.E., Stamatoyannopoulos, J.A., Noble, W.S.: Unsupervised segmentation of continuous genomic data. Bioinformatics 23(11), 1424–1426 (2007)

6. Biesinger, J., Wang, Y., Xie, X.: Discovering and mapping chromatin states using a tree hidden markov model. In: BMC Bioinformatics, vol. 14, pp. 1–11 (2013). Springer

7. Zhang, Y., Hardison, R.C.: Accurate and reproducible functional maps in 127 human cell types via 2d genome segmentation. Nucleic acids research 45(17), 9823–9836 (2017)

8. Zhang, Y., An, L., Yue, F., Hardison, R.C.: Jointly characterizing epigenetic dynamics across multiple human cell types. Nucleic acids research 44(14), 6721–6731 (2016)

9. Libbrecht, M.W., Chan, R.C., Hoffman, M.M.: Segmentation and genome annotation algorithms for identifying chromatin state and other genomic patterns. PLoS Computational Biology 17(10), 1009423 (2021)

10. Libbrecht, M.W., Rodriguez, O.L., Weng, Z., Bilmes, J.A., Hoffman, M.M., Noble, W.S.: A unified encyclopedia of human functional dna elements through fully automated annotation of 164 human cell types. Genome biology 20(1), 1–14 (2019)

11. Vu, H., Ernst, J.: Universal annotation of the human genome through integration of over a thousand epigenomic datasets. Genome Biology 23, 1–37 (2022)

12. Zhang, Y., Mahony, S.: Direct prediction of regulatory elements from partial data without imputation. PLoS computational biology 15(11), 1007399 (2019)

13. Boix, C.A., James, B.T., Park, Y.P., Meuleman, W., Kellis, M.: Regulatory genomic circuitry of human disease loci by integrative epigenomics. Nature 590(7845), 300–307 (2021)

14. Hoffman, M.M., Ernst, J., Wilder, S.P., Kundaje, A., Harris, R.S., Libbrecht, M., Giardine, B., Ellenbogen, P.M., Bilmes, J.A., Birney, E., et al.: Integrative annotation of chromatin elements from encode data. Nucleic acids research 41(2), 827–841 (2013)

15. Chan, R.C., Libbrecht, M.W., Roberts, E.G., Bilmes, J.A., Noble, W.S., Hoffman, M.M.: Segway 2.0: Gaussian mixture models and minibatch training. Bioinformatics 34(4), 669–671 (2018)

16. Impagliazzo, R., Lei, R., Pitassi, T., Sorrell, J.: Reproducibility in learning. In: Proceedings of the 54th Annual ACM SIGACT Symposium on Theory of Computing, pp. 818–831 (2022)

17. Consortium, E.P.: A user’s guide to the encyclopedia of dna elements (encode). PLoS biology 9(4), 1001046 (2011)

18. Li, Q., Brown, J.B., Huang, H., Bickel, P.J.: Measuring reproducibility of high-throughput experiments. The Annals of Applied Statistics 5(3), 1752–1779 (2011). doi:10.1214/11-AOAS466

19. Landt, S.G., Marinov, G.K., Kundaje, A., Kheradpour, P., Pauli, F., Batzoglou, S., Bernstein, B.E., Bickel, P., Brown, J.B., Cayting, P., et al.: Chip-seq guidelines and practices of the encode and modencode consortia. Genome research 22(9), 1813–1831 (2012)

20. Bailey, T., Krajewski, P., Ladunga, I., Lefebvre, C., Li, Q., Liu, T., Madrigal, P., Taslim, C., Zhang, J.: Practical guidelines for the comprehensive analysis of chip-seq data. PLoS computational biology 9(11), 1003326 (2013)

21. Yen, A., Kellis, M.: Systematic chromatin state comparison of epigenomes associated with diverse properties including sex and tissue type. Nature communications 6(1), 1–13 (2015)

22. Ebert, P., Schulz, M.H.: Fast detection of differential chromatin domains with sciddo. Bioinformatics 37(9), 1198–1205 (2021)

23. He, Y., Wang, T.: Epicompare: an online tool to define and explore genomic regions with tissue or cell type-specific epigenomic features. Bioinformatics 33(20), 3268–3275 (2017)

24. Ge, X., Zhang, H., Xie, L., Li, W.V., Kwon, S.B., Li, J.J.: Epialign: an alignment-based bioinformatic tool for comparing chromatin state sequences. Nucleic acids research 47(13), 77–77 (2019)

25. Jessa, S., Kleinman, C.L.: Chromswitch: a flexible method to detect chromatin state switches. Bioinformatics 34(13), 2286–2288 (2018)

26. Vu, H., Koch, Z., Fiziev, P., Ernst, J.: A framework for group-wise summarization and comparison of chromatin state annotations. Bioinformatics 39(1), 722 (2023)

27. Buske, O.J., Hoffman, M.M., Ponts, N., Le Roch, K.G., Noble, W.S.: Exploratory analysis of genomic segmentations with segtools. BMC bioinformatics 12(1), 1–7 (2011)

28. Daneshpajouh, H., Chen, B., Shokraneh, N., Masoumi, S., Wiese, K.C., Libbrecht, M.W.: Continuous chromatin state feature annotation of the human epigenome. Bioinformatics 38(11), 3029–3036 (2022)

29. Bayat, F., Libbrecht, M.: Vss: variance-stabilized signals for sequencing-based genomic signals. Bioinformatics 37(23), 4383–4391 (2021)

30. Xiang, G., Keller, C.A., Giardine, B., An, L., Li, Q., Zhang, Y., Hardison, R.C.: S3norm: simultaneous normalization of sequencing depth and signal-to-noise ratio in epigenomic data. Nucleic acids research 48(8), 43–43 (2020)

31. Arneson, A., Felsheim, B., Chien, J., Ernst, J.: Conshmm atlas: conservation state annotations for major genomes and human genetic variation. NAR Genomics and Bioinformatics 2(4), 104 (2020)

32. Libbrecht, M.W., Ay, F., Hoffman, M.M., Gilbert, D.M., Bilmes, J.A., Noble, W.S.: Joint annotation of chromatin state and chromatin conformation reveals relationships among domain types and identifies domains of cell-type-specific expression. Genome research 25(4), 544–557 (2015)

33. Farahbod, e.a. Marjan: ntegrative chromatin state annotation of 234 human encode4 cell types using segway reveals disease driver. In preparation (2023)

34. De Leeuw, J.: Correctness of kruskal’s algorithms for monotone regression with ties. Psychometrika 42(1), 141–144 (1977)

35. De Leeuw, J., Hornik, K., Mair, P.: Isotone optimization in r: pool-adjacent-violators algorithm (pava) and active set methods. Journal of statistical software 32, 1–24 (2010)

36. Chakravarti, N.: Isotonic median regression: a linear programming approach. Mathematics of operations research 14(2), 303–308 (1989)

37. Pedregosa, F., Varoquaux, G., Gramfort, A., Michel, V., Thirion, B., Grisel, O., Blondel, M., Prettenhofer, P., Weiss, R., Dubourg, V., et al.: Scikit-learn: Machine learning in python. the Journal of machine Learning research 12, 2825–2830 (2011)

38. O’Leary, N.A., Wright, M.W., Brister, J.R., Ciufo, S., Haddad, D., McVeigh, R., Rajput, B., Robbertse, B., Smith-White, B., Ako-Adjei, D., et al.: Reference sequence (refseq) database at ncbi: current status, taxonomic expansion, and functional annotation. Nucleic acids research 44(D1), 733–745 (2016)

39. Wagner, G.P., Kin, K., Lynch, V.J.: Measurement of mrna abundance using rna-seq data: Rpkm measure is inconsistent among samples. Theory in biosciences 131, 281–285 (2012)

