## Supplementary Material for "Robust chromatin state annotation"

### 1 Model training and annotation

#### 1.1 Segway

Segway requires inputs to be loaded on a Genomedata file [1]. We loaded all the track files belonging to each replicate on a Genomedata file. As training Segway on whole-genome is computationally expensive, we trained the Segway model on mini-batches of 3 percent of the genome [2, 3, 4] (Table 1), and we obtained genome annotations and posterior probability values.

The hyper-parameter `--minibatch-fraction` specifies the fraction of the genome that is randomly selected by Segway at each training iteration and used for training [3].

With default parameter settings, Segway's posterior probabilities lack numerical precision due to underflow and overflow of floating points such that >90% of genomic positions receive either exactly zero or exactly one posterior value. To mitigate this issue and to enable downstream reproducibility analysis based on granular posterior probability values, we need to tune hyper-parameters. To tackle this issue and to obtain granular probability values, we artificially softened the model's posterior by raising the emission probabilities (`--track-weights`) to the power of $e = 0.01$ ($\frac{1}{\texttt{--resolution}}$) and by doing so, we uniformly decreased the model's confidence. The hyper-parameter `--track-weights` is the exponent of emission probability, and the modified posterior values are calculated as follows:

$$P(Q_{1,..,T} = q_{1,..,T}|X_{1,..,T}) = \prod_{t=1}^{T} P(Q_t = q_t|Q_{t-1})P(X_t|Q_t = q_t)^e \tag{1}$$

Where, $P(X_t|Q_t = q_t)$ and $P(Q_t = q_t|Q_{t-1})$ are emission and transition probabilities, respectively, and 0.01 which is the exponent of emission probability is specified to the model through the hyper-parameter `--track-weights`. This additional step mitigates the issue of lack of granularity by increasing the number of positions with granular posterior value. Moreover, to specify the number of labels, we used the formula $(10 + 2\sqrt{\text{number of assays}})$ as suggested by [5] to scale with the amount of available data. For all of our collected datasets with 10 to 13 histone modification assays, the number of labels is 16. Generally, having longer segments is desired for Segway [4, 3] therefore, this method uses hard and soft constraints for controlling the length distribution of segments. The hyper-parameters `--prior-strength` and `--segtransition-weight-scale` are the soft constraints and act as length prior and exponent of transition probability, respectively. For Segway, we mostly followed the hyper-parameter settings of [5].

| Hyper-parameter | Value |
|---|---|
| `--resolution` | 100 |
| `--num-labels` | $10 + 2\sqrt{\text{number of assays}}$ |
| `--track-weight` | $\frac{1}{100}$ |
| `--segtransition-weight-scale` | 1 |
| `--prior-strength` | 1 |
| `--minibatch-fraction` | 0.03 |

Table 1: Hyper-parameters used to train Segway models

#### 1.2 ChromHMM

For ChromHMM we followed standard practice and trained the models with default hyper-parameter setting and on the whole genome. For ChromHMM runs, the `--resolution = 200` and the model is trained on whole genome. As a pre-processing step, ChromHMM binarizes the signals throughout the genome, using a hard threshold on the Poisson distribution probability [6], with each position having an indicator of the presence or absence of a certain epigenetic mark. Lastly, trained ChromHMM models provided genome annotations and their corresponding posterior probabilities.

#### 1.3 Interpretation of Chromatin State Types

The table below provides an interpretation of chromatin state types in terms of their biological roles. These interpretations were generated using an automated method described in [5], which predicts the biological interpretation of each label based on its enrichment around conserved regions and its association with different histone modification marks. It is important to note that these interpretation terms are provided for the purpose of making the results more interpretable, and all steps of the reproducibility evaluation pipeline are independent of these terms.

| **Chromatin State Type** | **Short Name** | **Description** |
|---|---|---|
| Promoter | Prom | Presence of promoter-associated marks H3K4me3 and H3K9ac. Highly enriched at transcription start sites (TSSs). |
| PromoterFlanking | Prom_fla | Presence of promoter-associated marks H3K4me3 and H3K9ac, but at lower levels than Promoters. Tends to occur upstream or downstream of TSSs. |
| Enhancer | Enha | Presence of the enhancer-associated marks H3K27ac and H3K4me1. |
| EnhancerLow | Enha_low | Same as Enhancer, but with lower signal values. |
| Bivalent | Biva | Presence of both activating (H3K27ac) and repressive (H3K27me3) marks. Thought to mark regulatory elements “poised” for activation. |
| CTCF | | Presence of the transcription factor CTCF, thought to play a role in chromatin conformation. In samples without measured CTCF, this label marks positions with CTCF-associated marks. |
| Transcribed | Tran | Characterized by the transcription-associated mark H3K36me3. Highly enriched in annotated gene bodies. |
| K9K36 | K9K36 | Presence of the marks H3K9me3 and H3K36me3, a pattern associated with zinc finger genes. |
| FacultativeHet | Facu | Facultative (Polycomb) Heterochromatin, characterized by the histone modification H3K27me3. Thought to carry out cell type-specific repression. |
| ConstitutiveHet | Cons | Constitutive Heterochromatin, characterized by the histone modification H3K9me3. Marks permanently silent regions such as centromeres and telomeres. |
| Quiescent | Quie | Lack of any marks |

### 2 Comprehensive evaluation of chromatin state reproducibility

#### 2.1 Setting 1: Different Data, Different Models

##### 2.1.1 ChromHMM

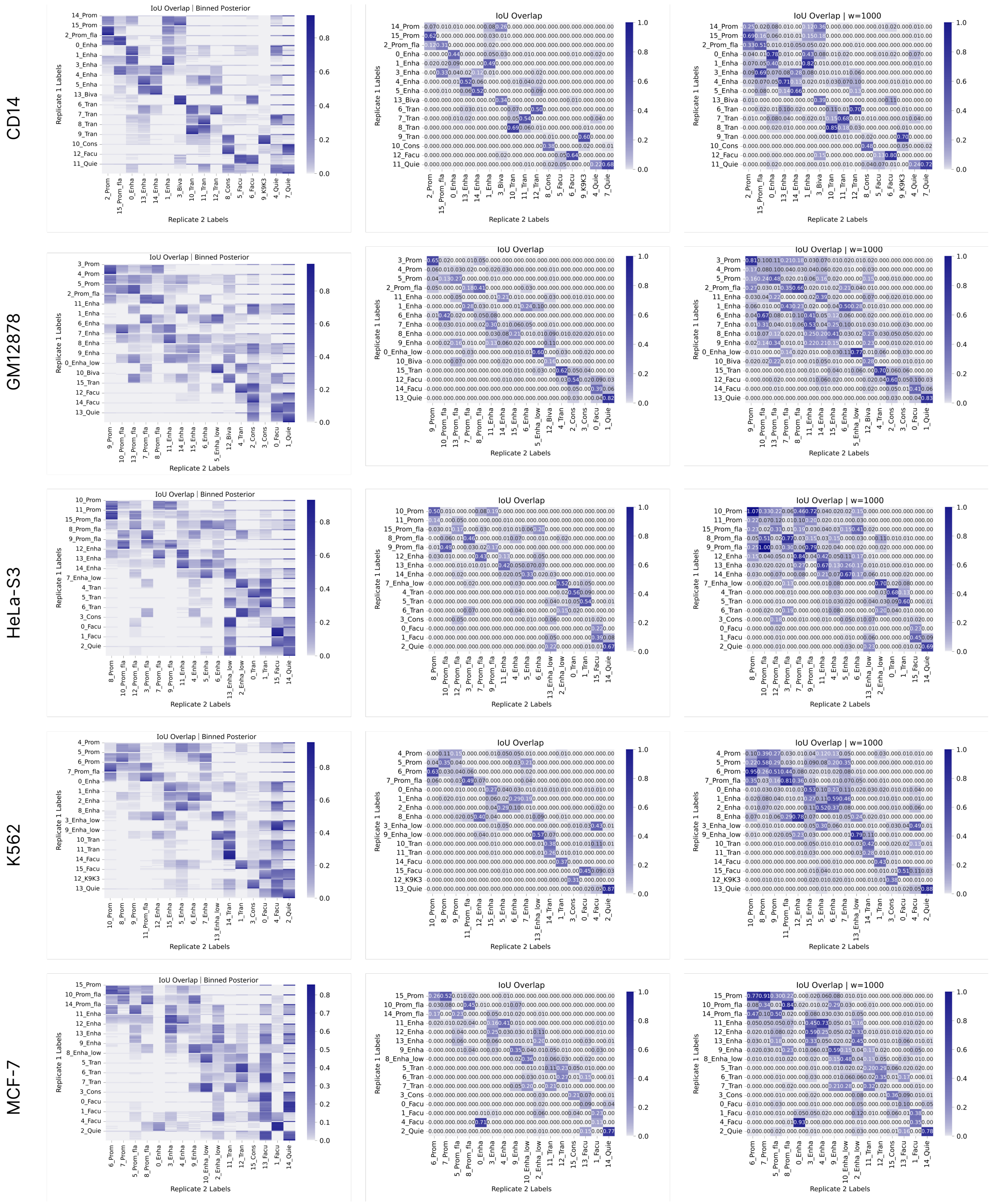


Figure 1: Intersection over union (IoU) overlap of ChromHMM according to setting 1 (different data, different models). Columns from left to right are binned posterior, IoU overlap with maximum-a-posterior state with $w = 0$, IoU overlap with maximum-a-posterior state with $w = 1000$. Color axis is linear and rows correspond to five cell types.

##### 2.1.2 Segway

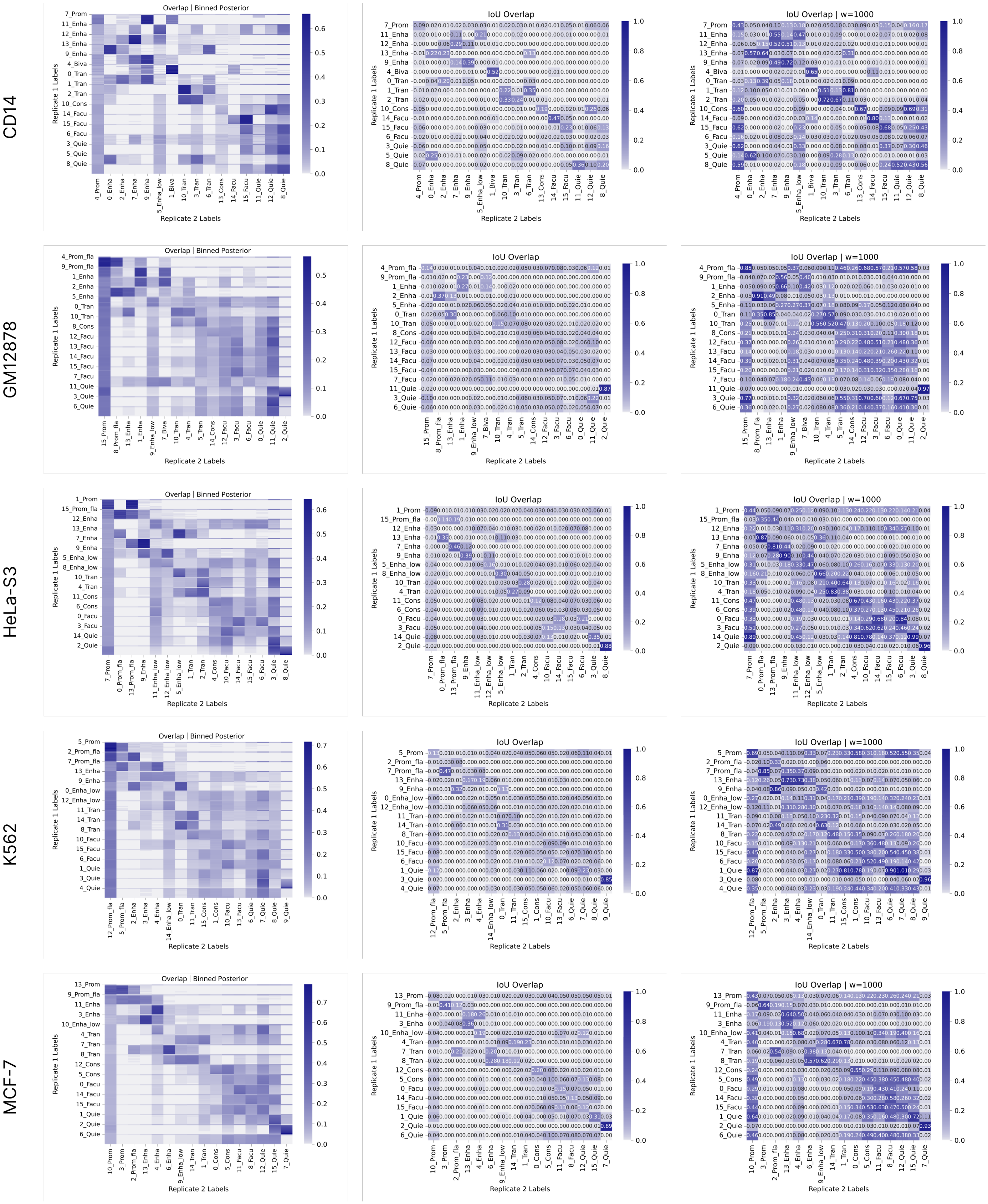


Figure 2: Intersection over union (IoU) overlap of Segway according to setting 1 (different data, different models). Columns from left to right are binned posterior, IoU overlap with maximum-a-posterior state with $w = 0$, IoU overlap with maximum-a-posterior state with $w = 1000$. Color axis is linear and rows correspond to five cell types.

#### 2.2 Setting 2: Different Data, Same Model

##### 2.2.1 ChromHMM

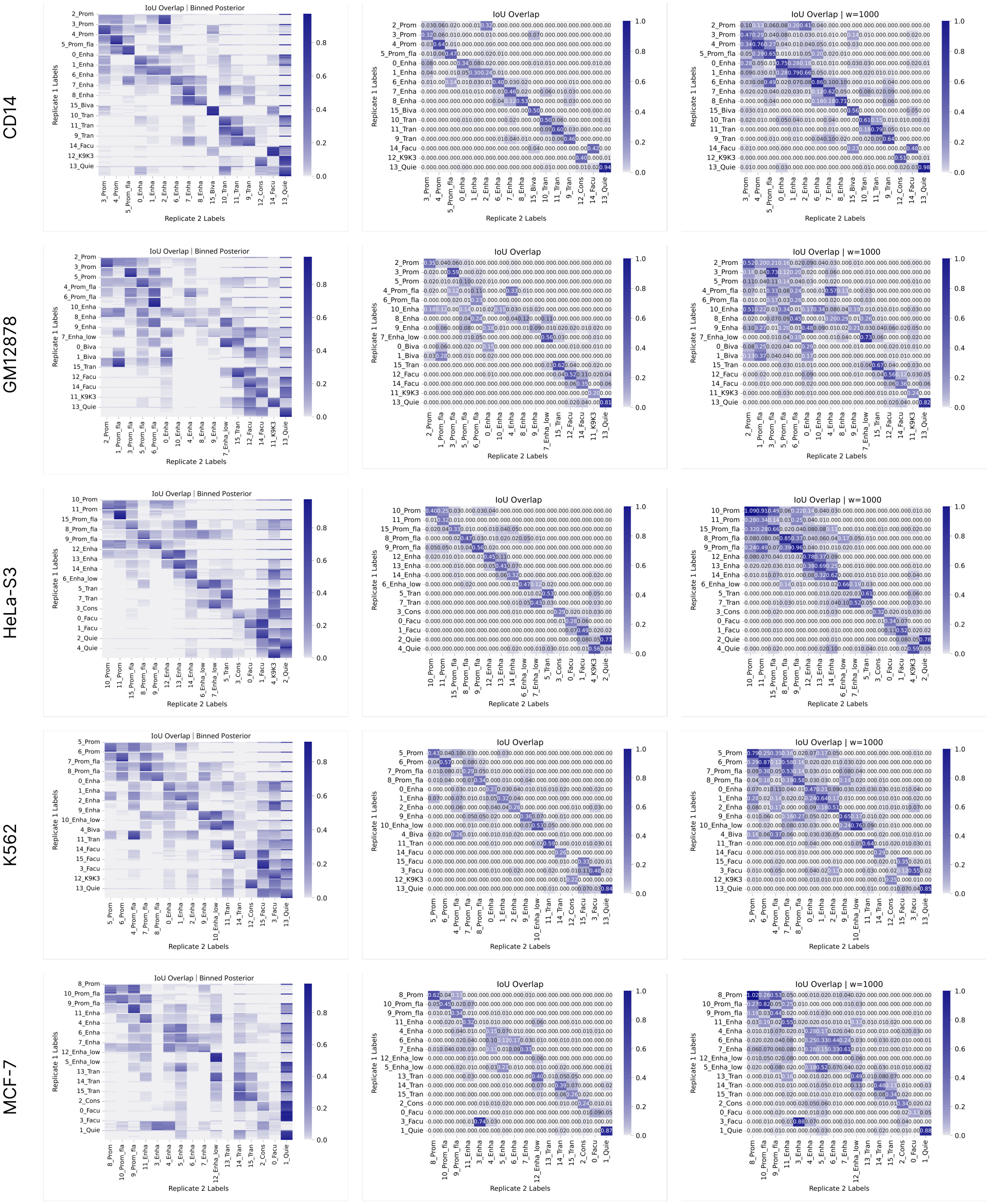


Figure 3: Intersection over union (IoU) overlap of ChromHMM according to (setting 2: different data, same model). Columns from left to right are binned posterior, IoU overlap with maximum-a-posterior state with $w = 0$, IoU overlap with maximum-a-posterior state with $w = 1000$. Color axis is linear and rows correspond to five cell types.

##### 2.2.2 Segway

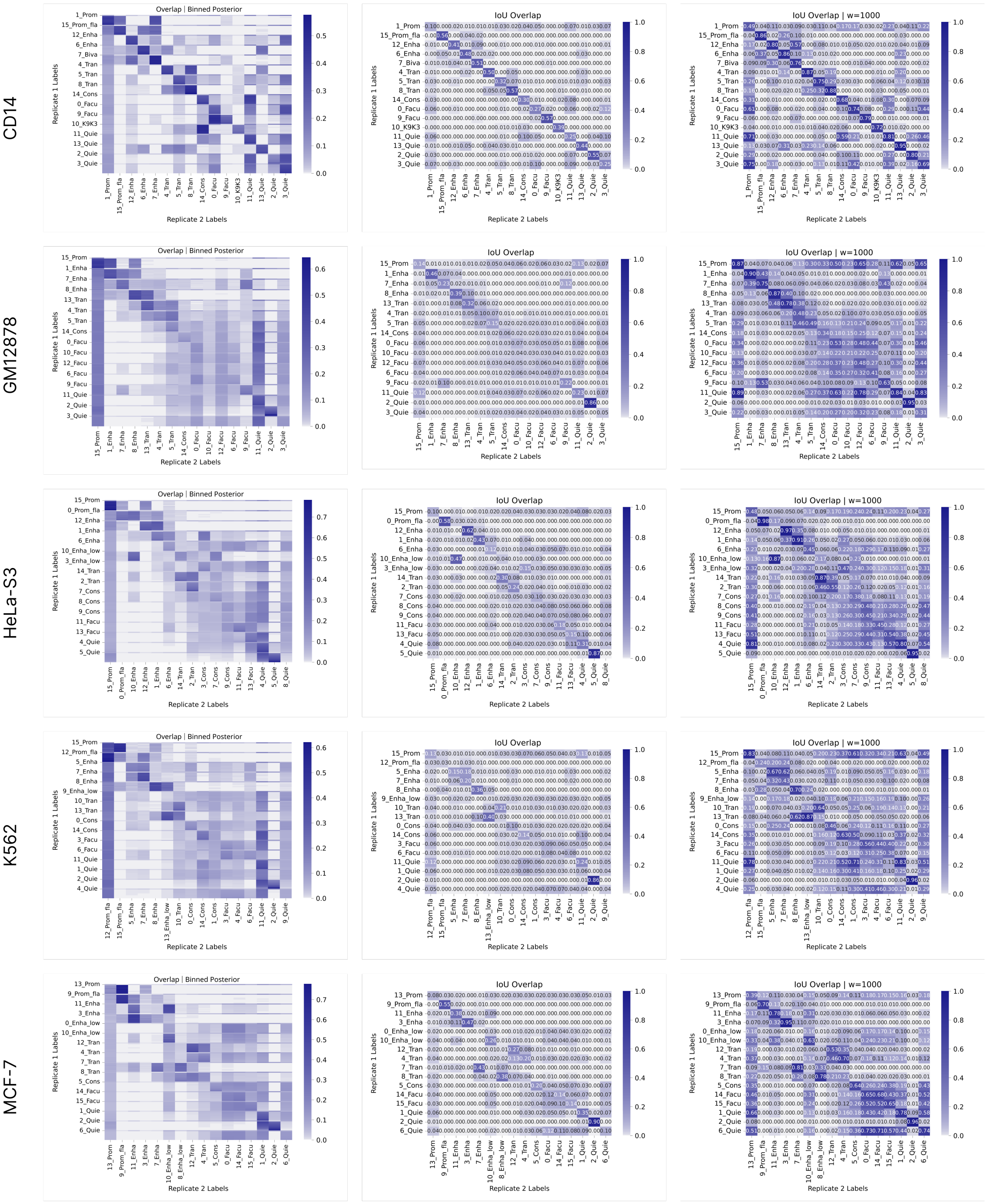


Figure 4: Intersection over union (IoU) overlap of Segway according to (setting 2: different data, same model). Columns from left to right are binned posterior, IoU overlap with maximum-a-posterior state with $w = 0$, IoU overlap with maximum-a-posterior state with $w = 1000$. Color axis is linear and rows correspond to five cell types.

#### 2.3 Setting 3: Same Data, Different Models

##### 2.3.1 ChromHMM

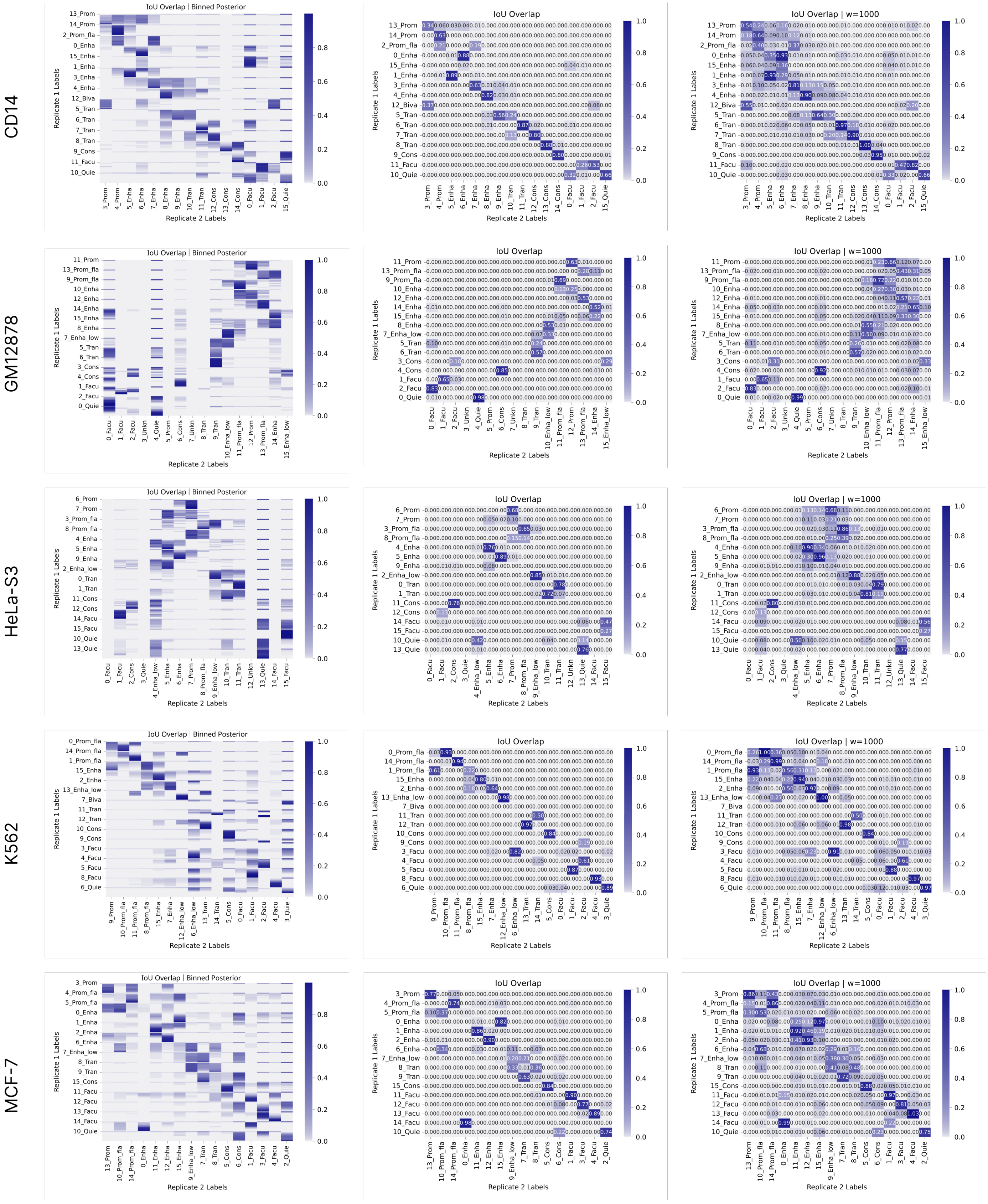


Figure 5: Intersection over union (IoU) overlap of ChromHMM according to (Setting 3: Same Data, Different Models). Columns from left to right are binned posterior, IoU overlap with maximum-a-posterior state with $w = 0$, IoU overlap with maximum-a-posterior state with $w = 1000$. Color axis is linear and rows correspond to five cell types.

##### 2.3.2 Segway

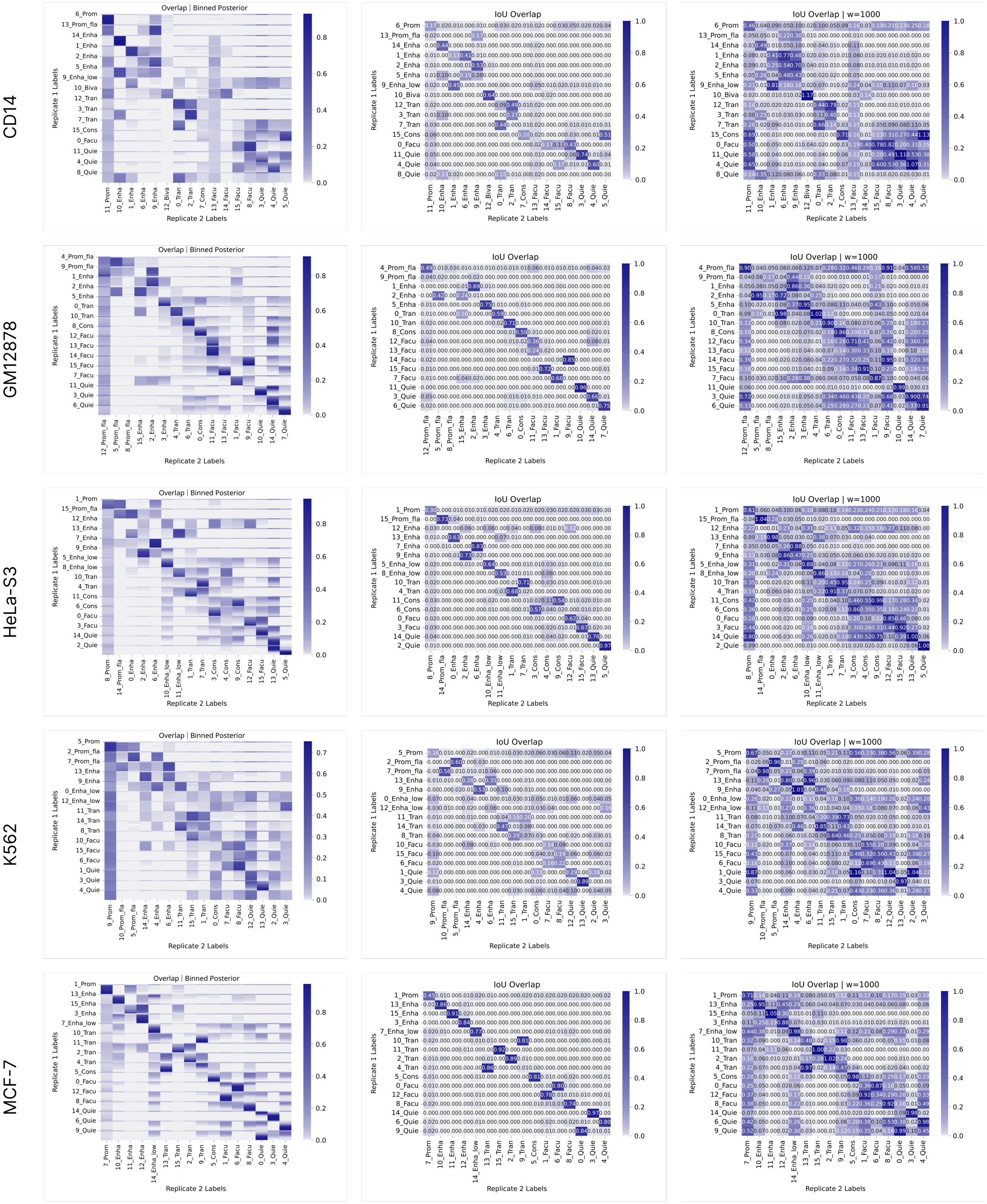


Figure 6: Intersection over union (IoU) overlap of Segway according to (Setting 3: Same Data, Different Models). Columns from left to right are binned posterior, IoU overlap with maximum-a-posterior state with $w = 0$, IoU overlap with maximum-a-posterior state with $w = 1000$. Color axis is linear and rows correspond to five cell types.

### 3 Chromatin states overlap differently across replicates

#### 3.1 Naive overlap

##### 3.1.1 ChromHMM

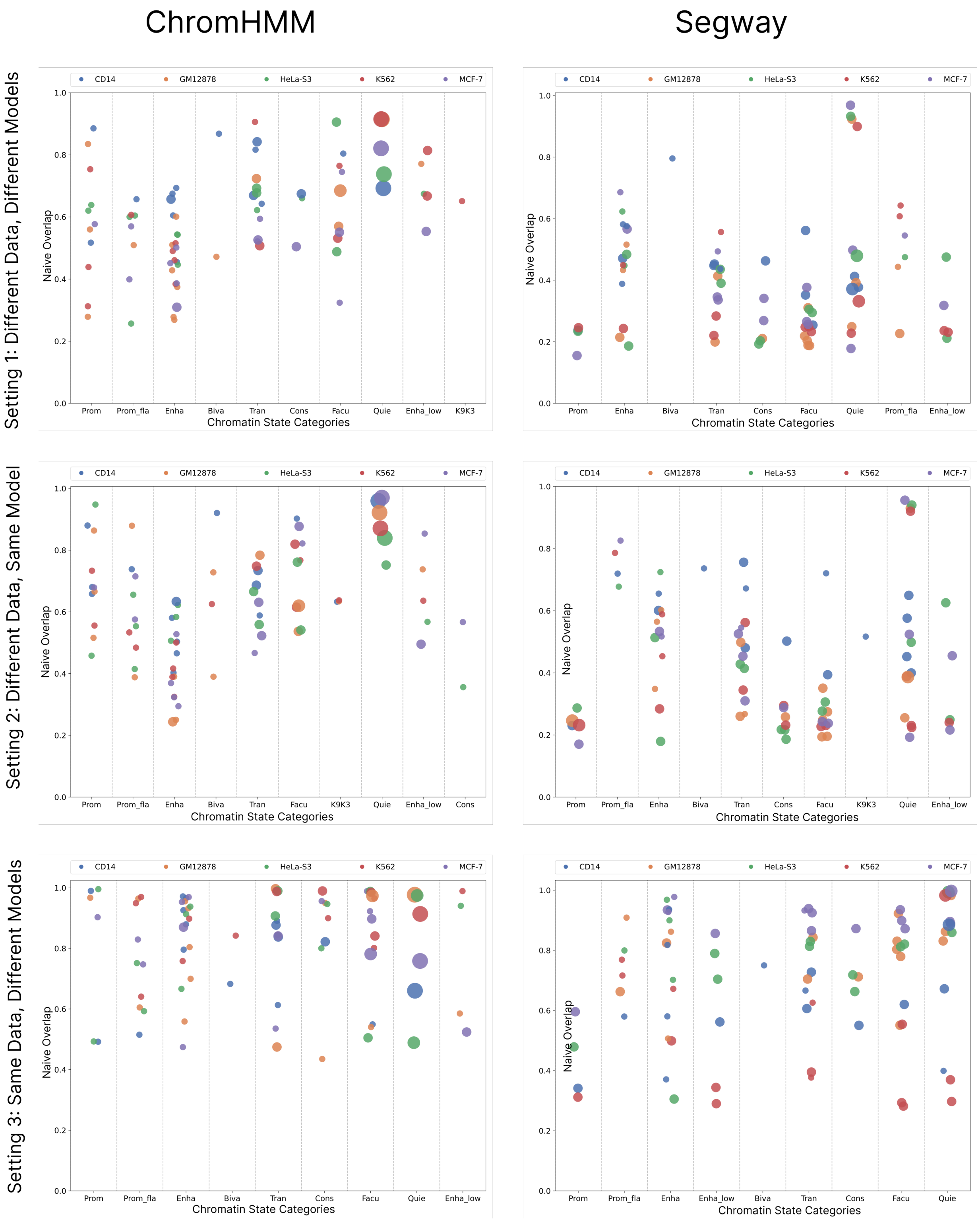


Figure 7: Fraction of overlap (Naive overlap) of various chromatin states categories identified in the ChromHMM and Segway annotation according to three settings for all five cell types. Each dot represents a chromatin state, with color denoting cell type and size proportional to genome coverage

#### 3.2 Granularity of Chromatin States

##### 3.2.1 ChromHMM

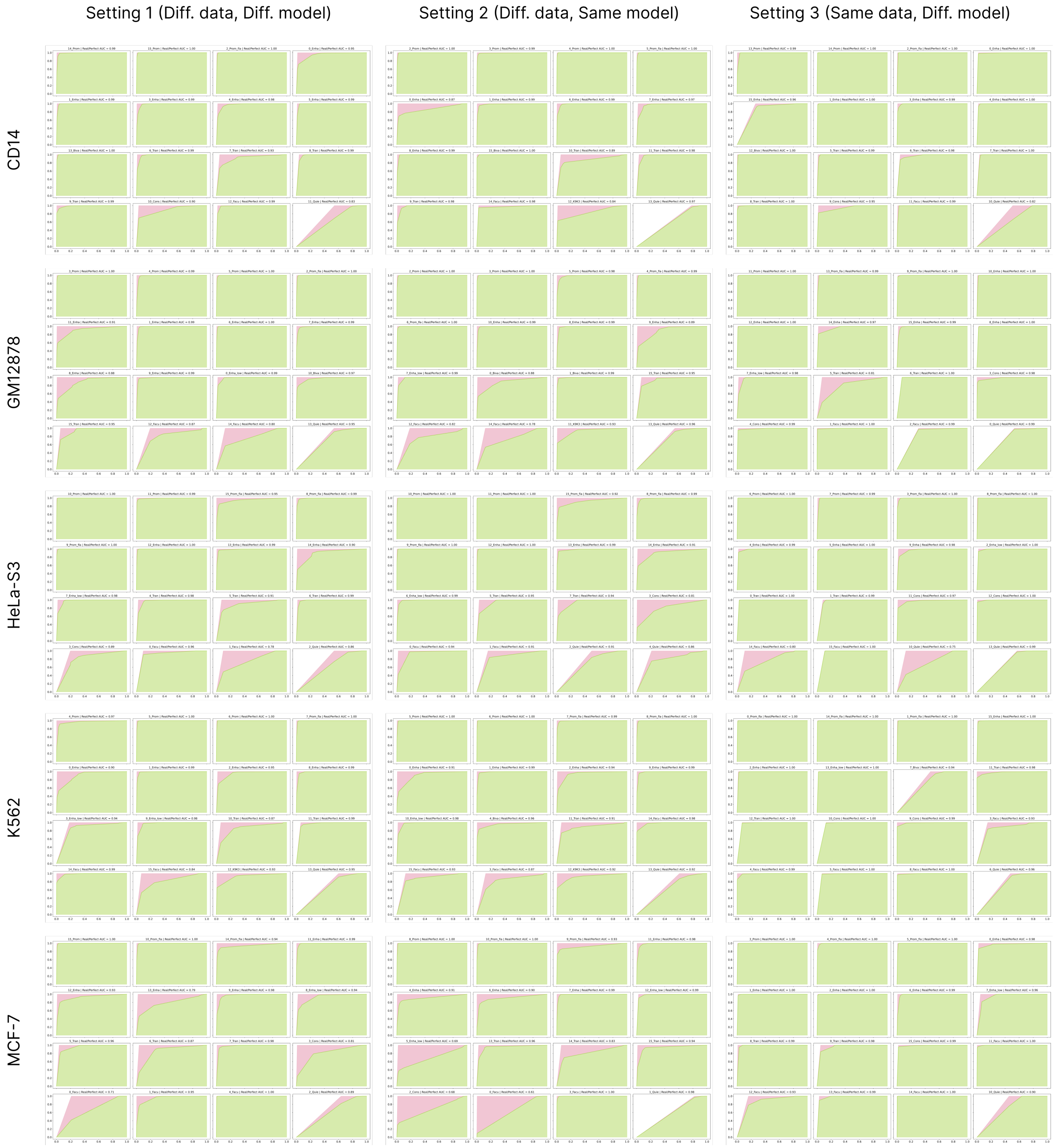


Figure 8: State merging curve for ChromHMM runs. Columns correspond to different settings and rows correspond to five cell types. The area under the state merging curve (auSMC) ratio is a numerical representation of chromatin state reproducibility as a function of chromatin state granularity which is calculated by dividing the observed AUC by the AUC pertaining to the perfect reproducibility case.

##### 3.2.2 Segway

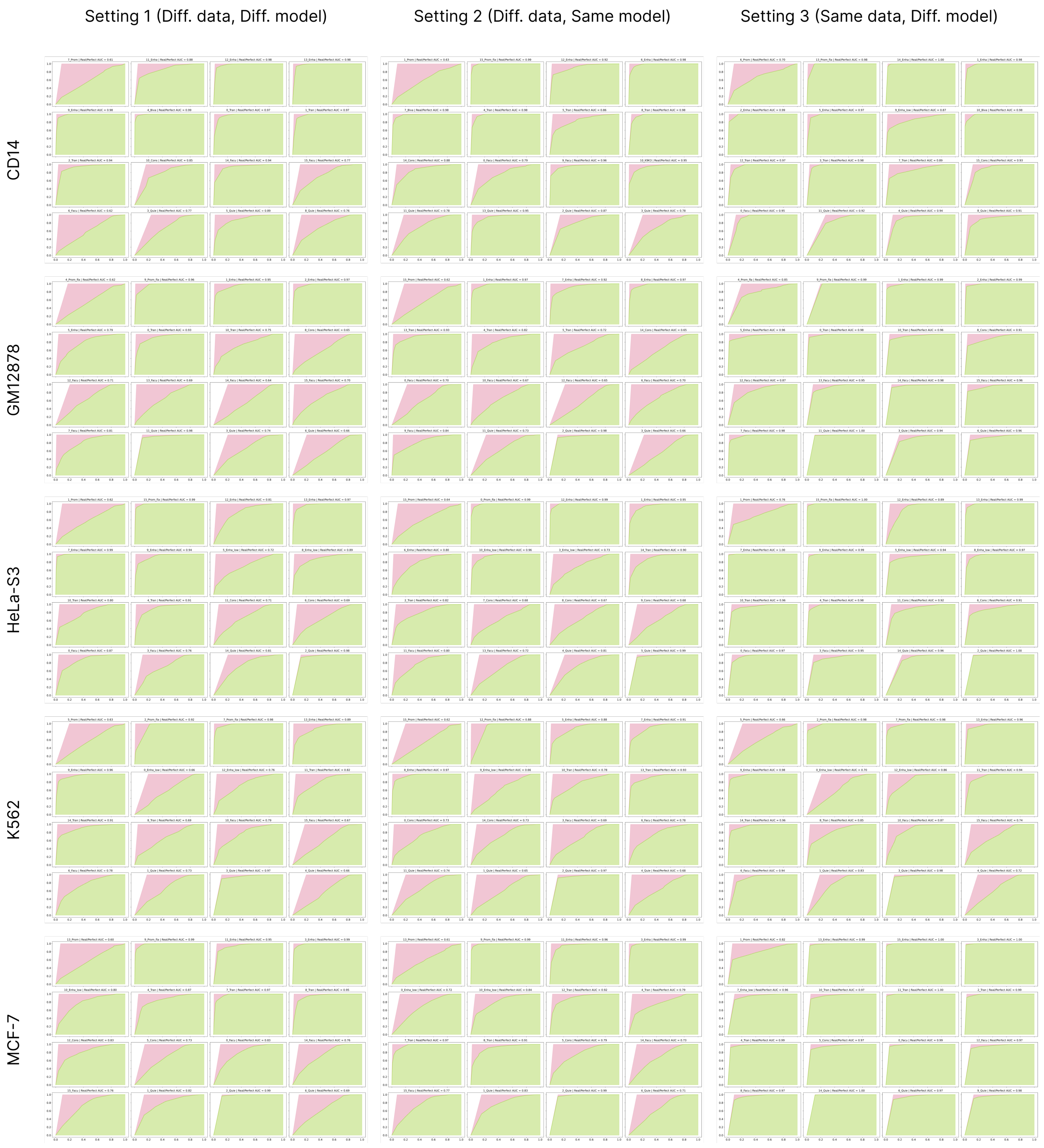


Figure 9: State merging curve for Segway runs. Columns correspond to different settings and rows correspond to five cell types. The area under the state merging curve (auSMC) ratio is a numerical representation of chromatin state reproducibility as a function of chromatin state granularity which is calculated by dividing the observed AUC by the AUC pertaining to the perfect reproducibility case.

#### 3.3 Area under state merging curve (auSMC) for various genomic functions

##### 3.3.1 ChromHMM

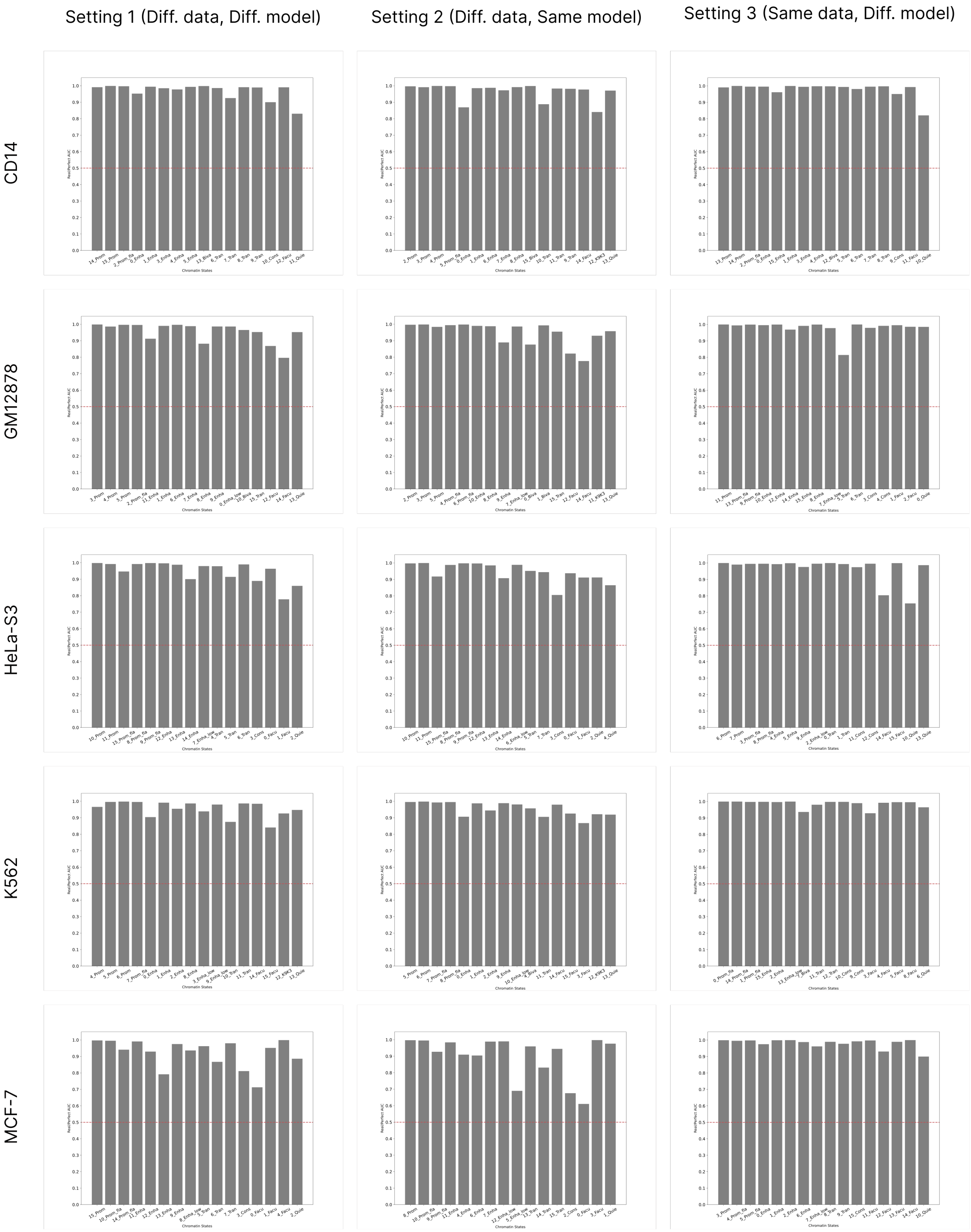


Figure 10: The auSMC ratio of chromatin states in the base ChromHMM annotations. Rows correspond to different cell types and columns correspond to different settings of variability.

##### 3.3.2 Segway

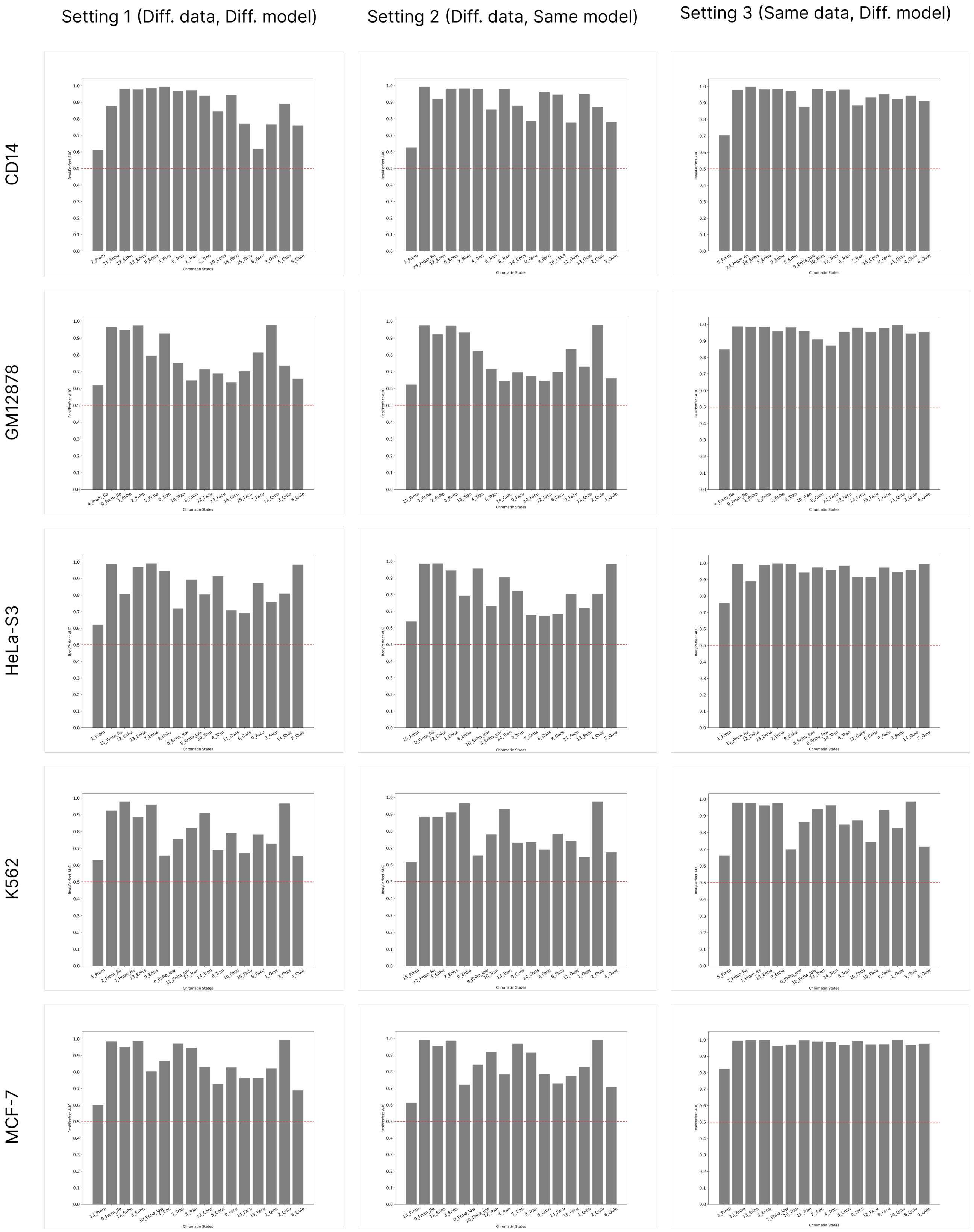


Figure 11: The auSMC ratio of chromatin states in the base Segway annotations. Rows correspond to different cell types and columns correspond to different settings of variability.

##### 3.3.3 Area under state merging curve (auSMC) chromatin state categories

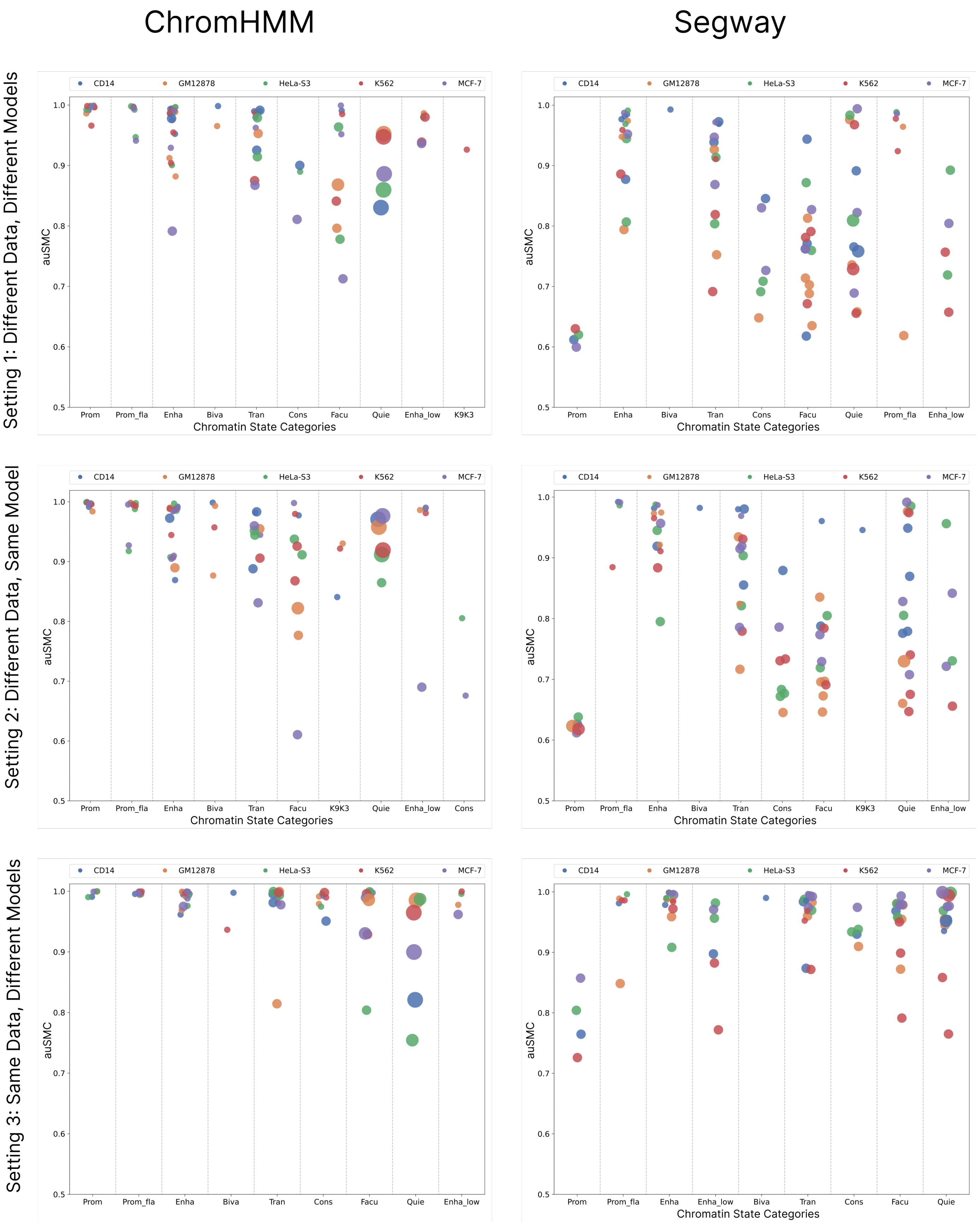


Figure 12: auSMC ratio of various chromatin states categories identified in the ChromHMM (left column) and Segway (right column) annotation according to three settings of variability (rows) for five cell types. Each dot represents a chromatin state, with color denoting cell type and size proportional to genome coverage.

#### 3.4 Post-Clustering

##### 3.4.1 Mutual Information

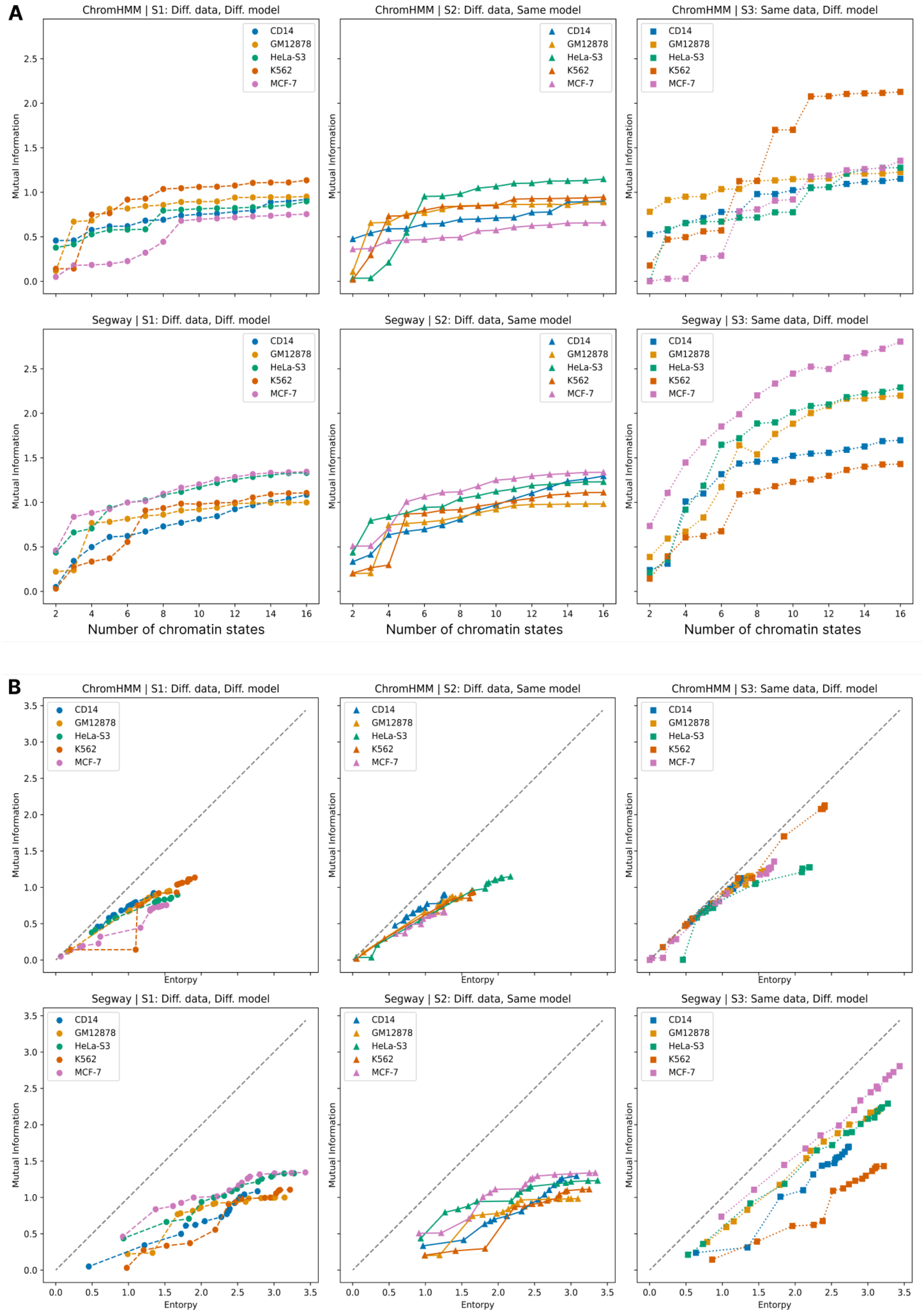


Figure 13: **(A)** Mutual information as a function of number of states of annotations measured during the post-clustering process. **(B)** Mutual information as a function of entropy of annotations measured during the post-clustering process.

##### 3.4.2 Average r-value

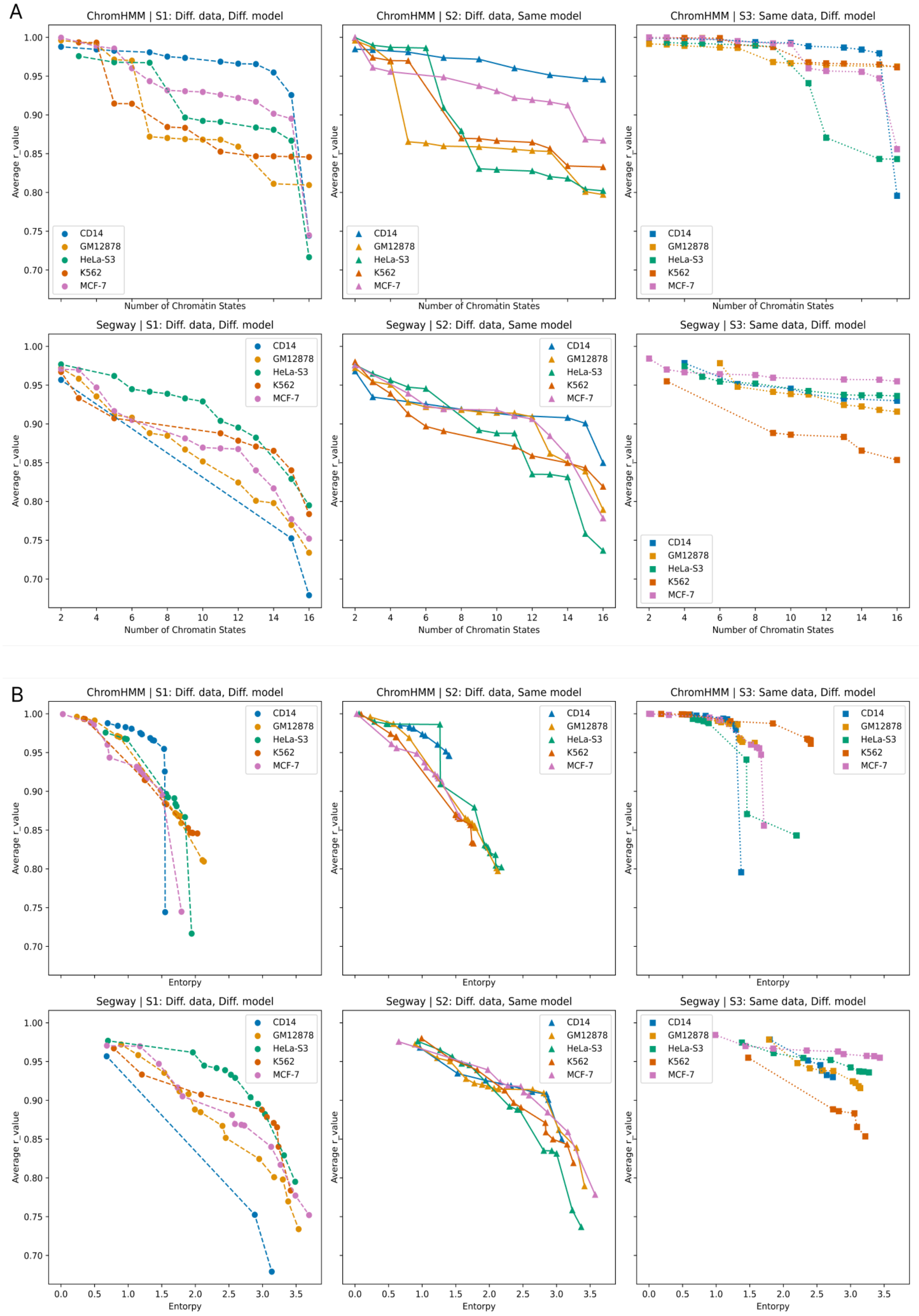


Figure 14: **(A)** Average r-value as a function of number of states of annotations measured during the post-clustering process. **(B)** Average r-value as a function of entropy of annotations measured during the post-clustering process.

##### 3.4.3 Fraction of confident genomic regions

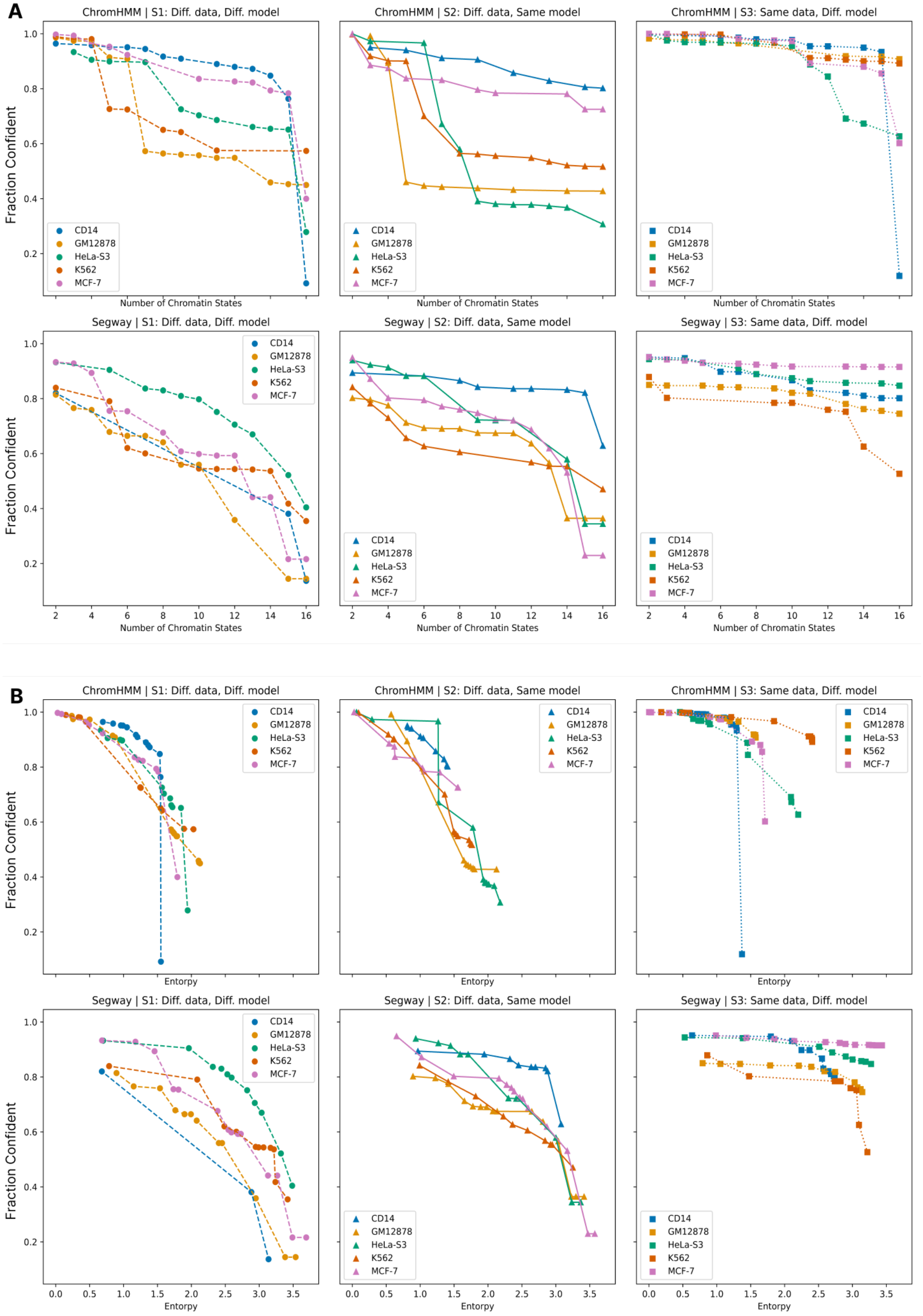


Figure 15: **(A)** Fraction of confident genomic regions as a function of number of states of annotations measured during the post-clustering process. **(B)**Fraction of confident genomic regions as a function of entropy of annotations measured during the post-clustering process.

### 4 Corresponding chromatin states occur at proximity of each other across replicates

#### 4.1 Overlap as a function of $w$

##### 4.1.1 ChromHMM

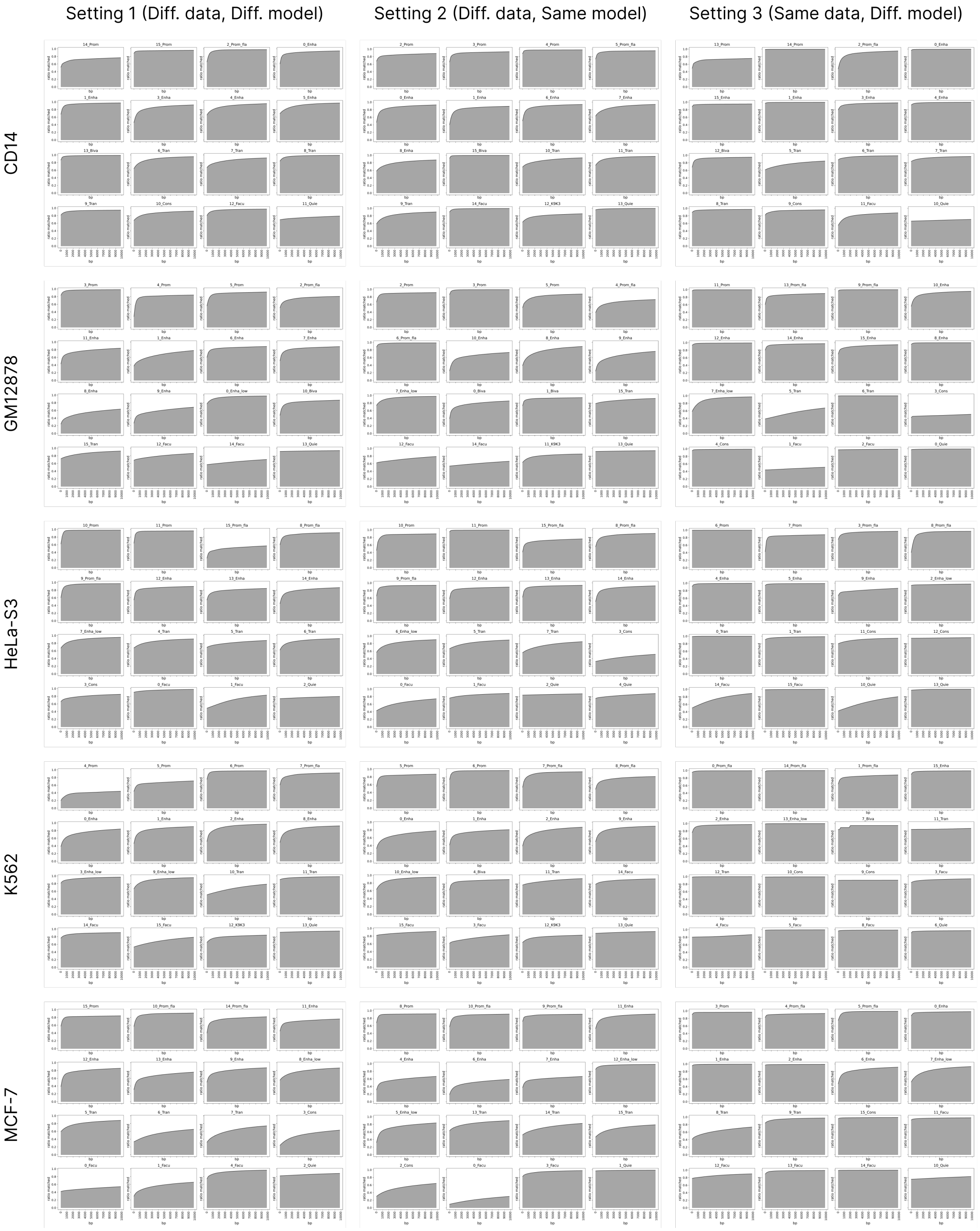


Figure 16: The fraction of each state in base annotation that is overlapped with a corresponding state within a $w$-sized distance upstream or downstream in ChromHMM. Rows correspond to different cell types and columns correspond to settings of variability.

##### 4.1.2 Segway

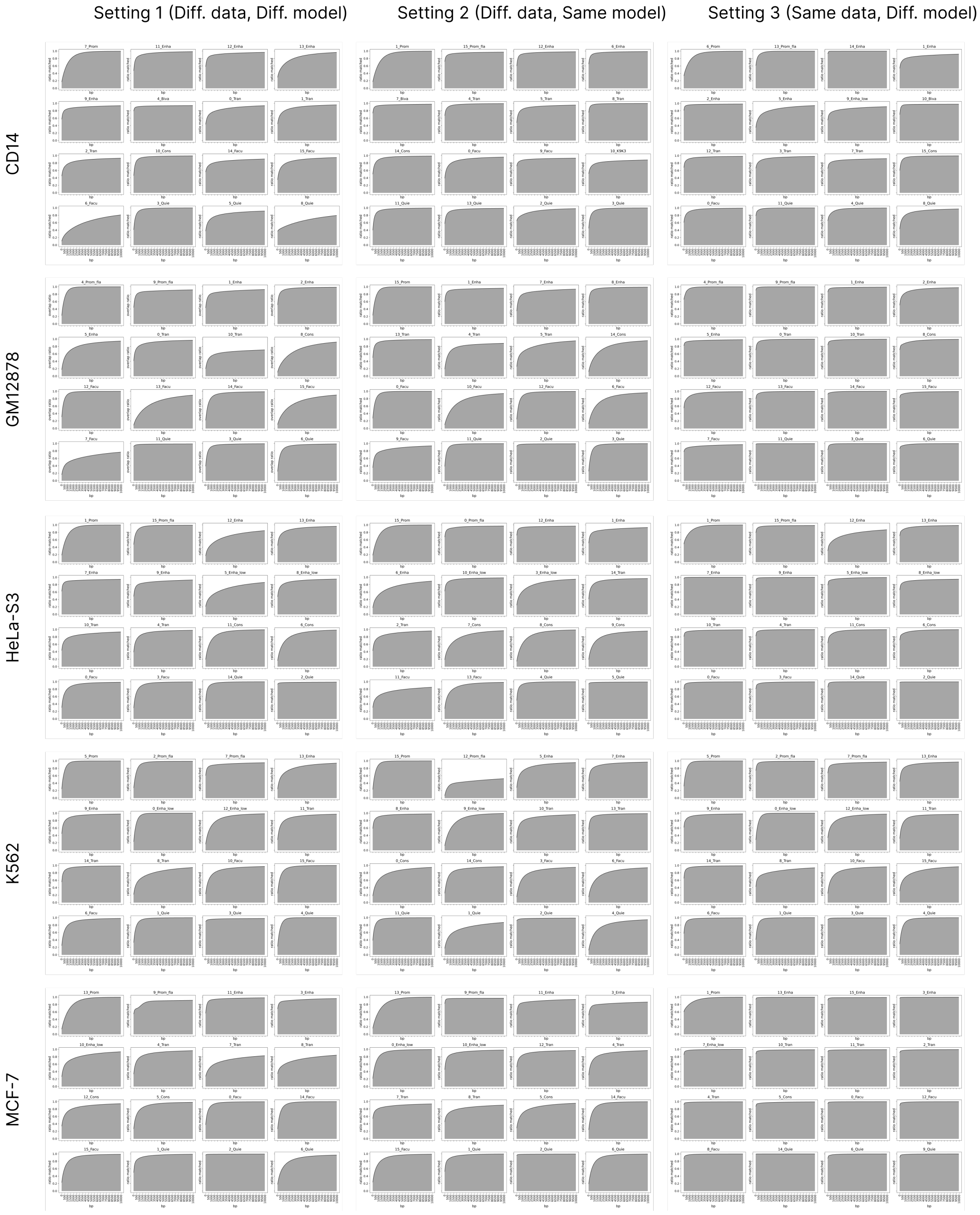


Figure 17: The fraction of each state in base annotation that is overlapped with a corresponding state within a $w$-sized distance upstream or downstream in Segway. Rows correspond to different cell types and columns correspond to settings of variability.

#### 4.2 Correspondence as a function of $w$

##### 4.2.1 ChromHMM

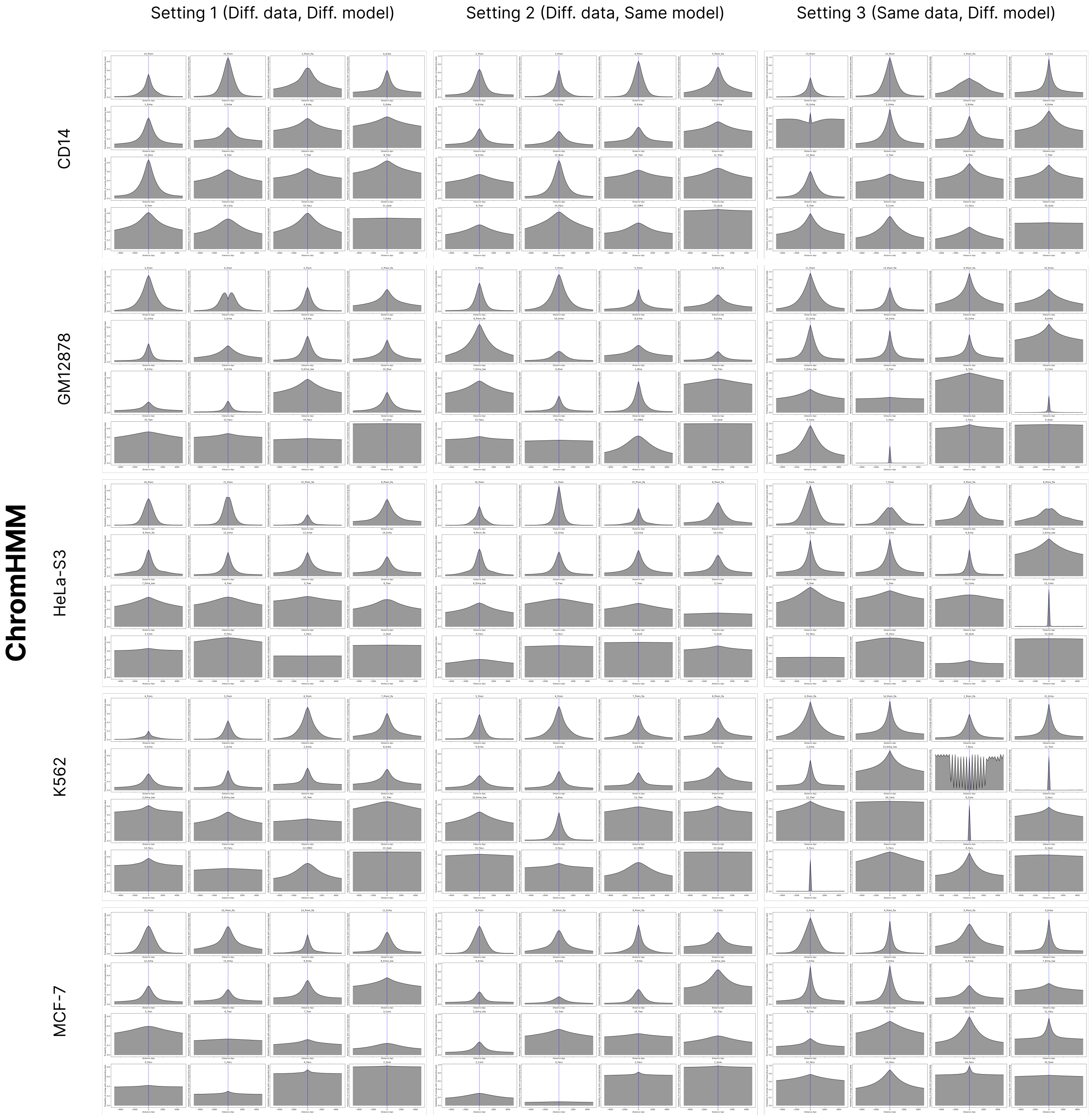


Figure 18: At each position in the genome, the probability of overlap with the corresponding state decreases as a function of distance from the position. Rows correspond to different cell types and columns correspond to settings of variability.

###### 4.2.2 Segway

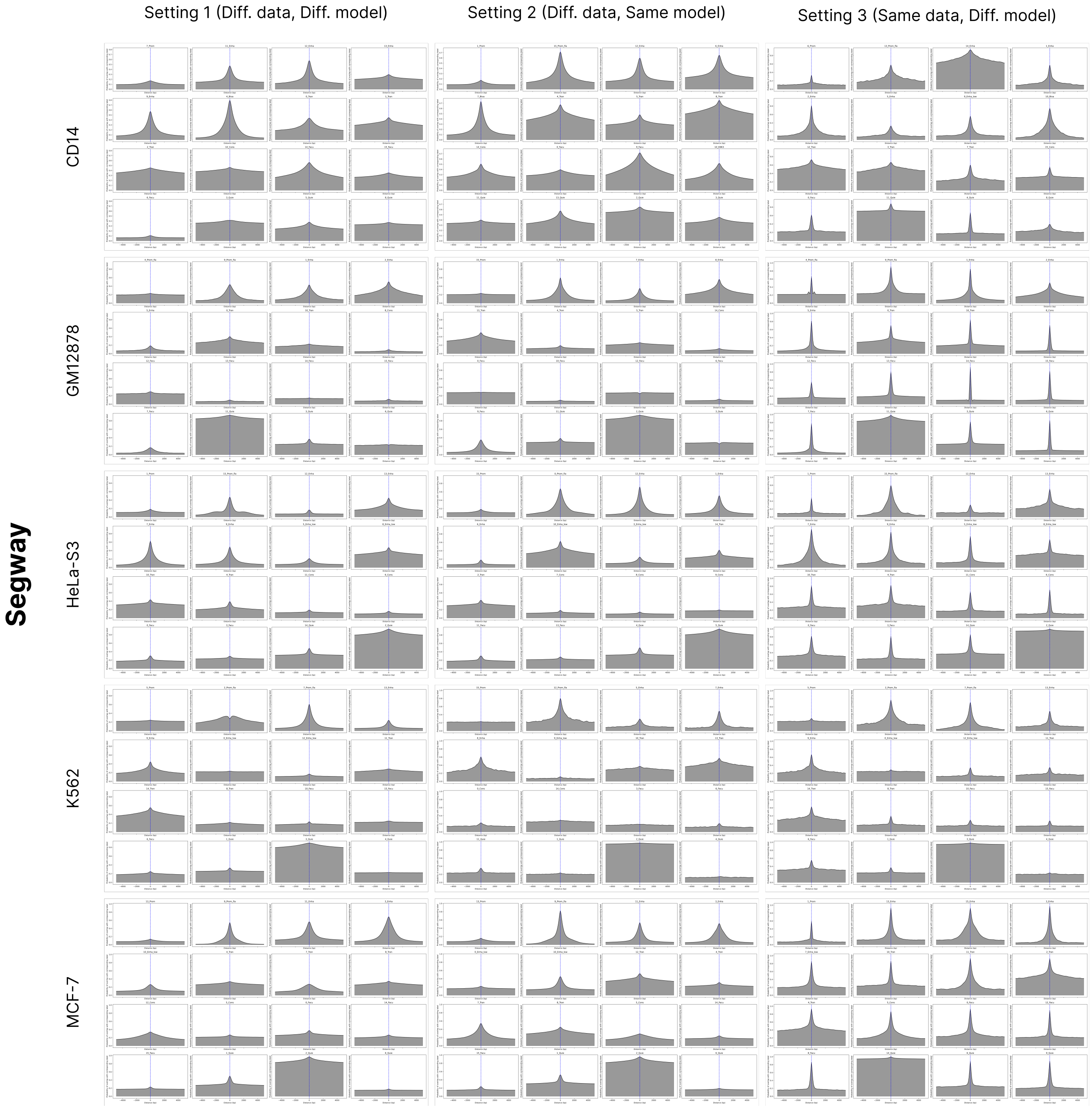


Figure 19: At each position in the genome, the probability of overlap with the corresponding state decreases as a function of distance from the position. Rows correspond to different cell types and columns correspond to settings of variability.

### 5 Posterior probability is associated with overlap

#### 5.1 Calibration of posterior probability

##### 5.1.1 ChromHMM

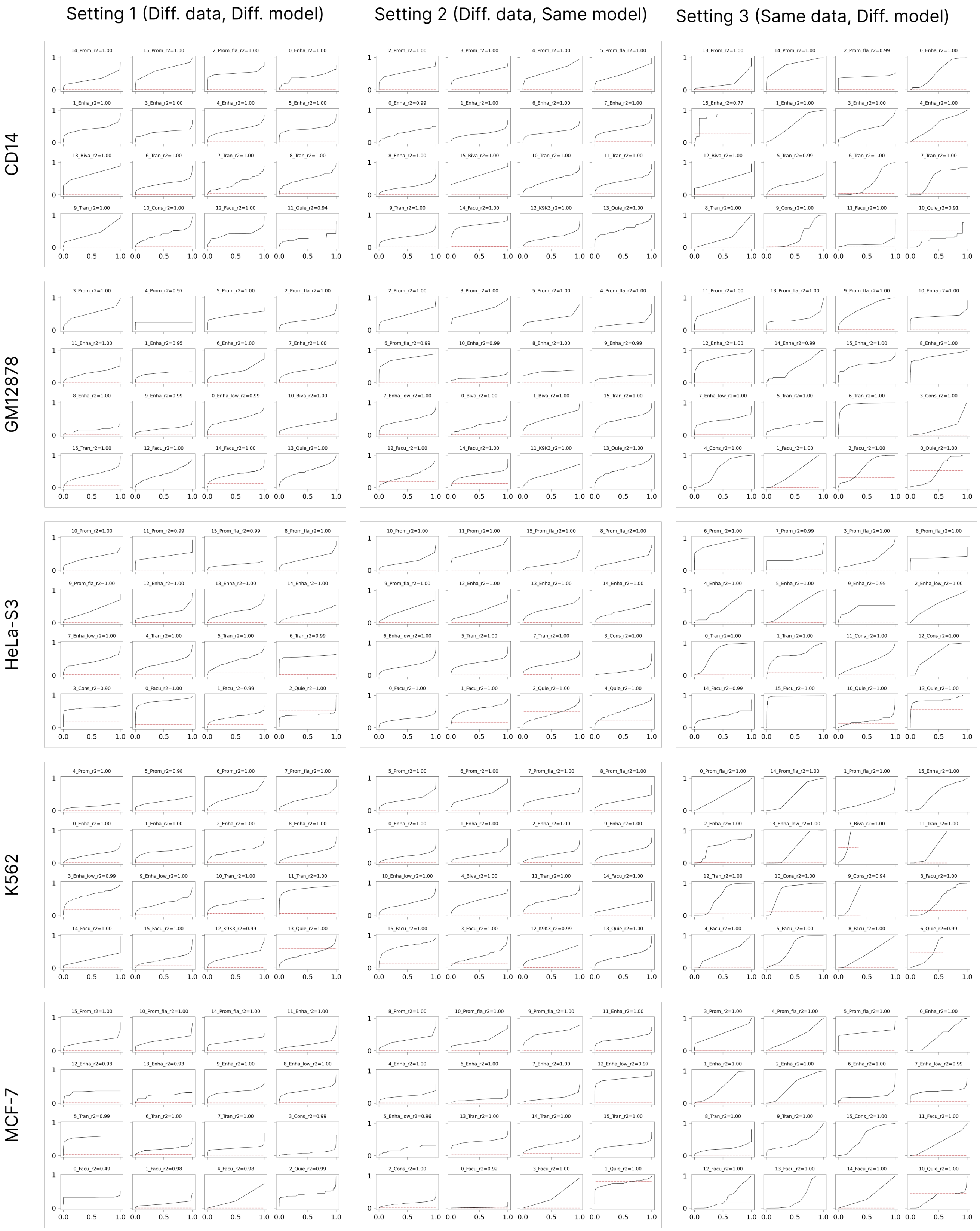


Figure 20: For each chromatin state in the base annotation, we measure the frequency of overlap with a corresponding state as a function of their posterior probability. We observe a clear association between the overlap and the posterior probability. Rows correspond to different cell types and columns correspond to settings of variability.

##### 5.1.2 Segway

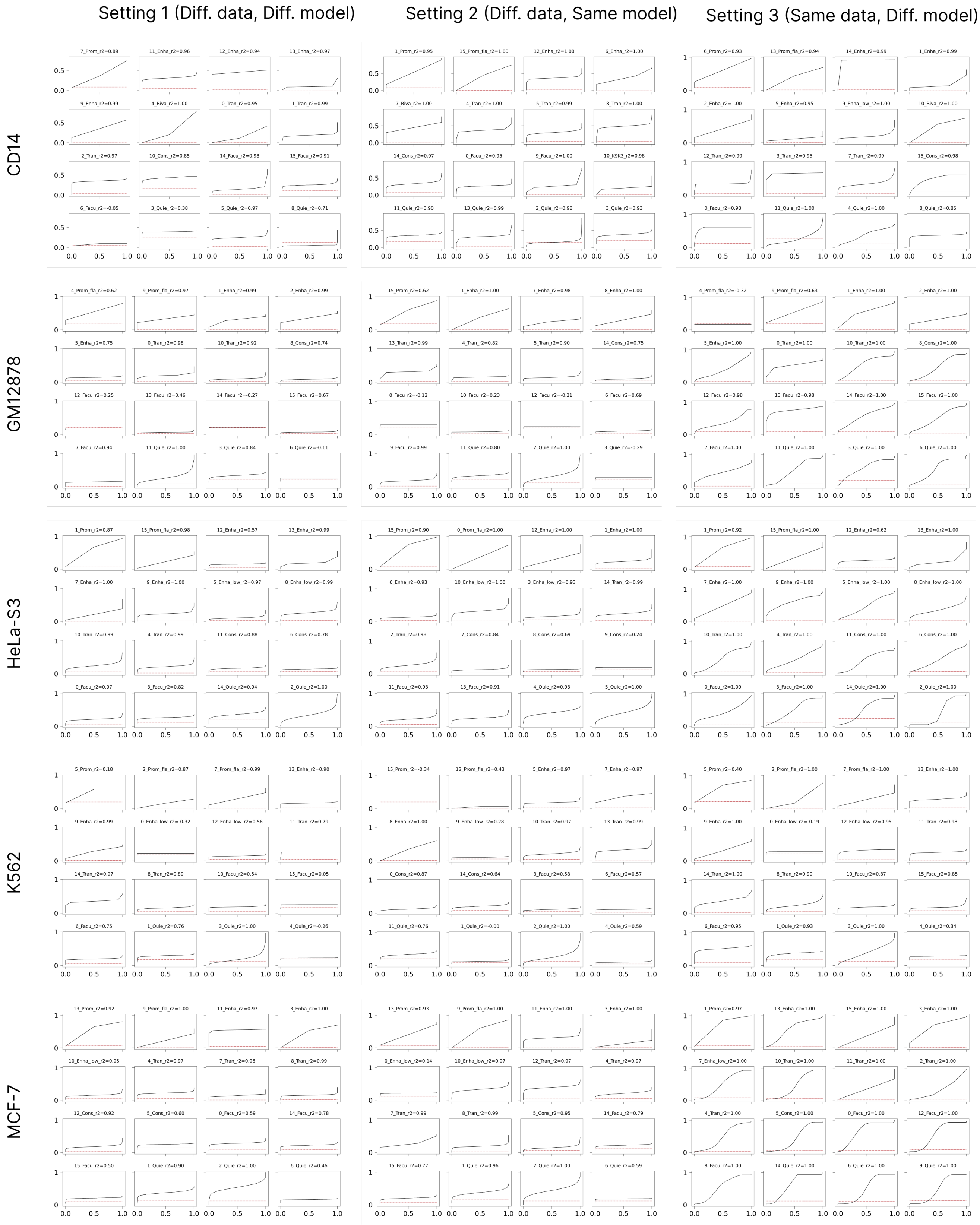


Figure 21: For each chromatin state in the base annotation, we measure the frequency of overlap with a corresponding state as a function of their posterior probability. We observe a clear association between the overlap and the posterior probability. Rows correspond to different cell types and columns correspond to settings of variability.

#### 5.2 Transcription (gene body) as a function of posterior probability

##### 5.2.1 ChromHMM

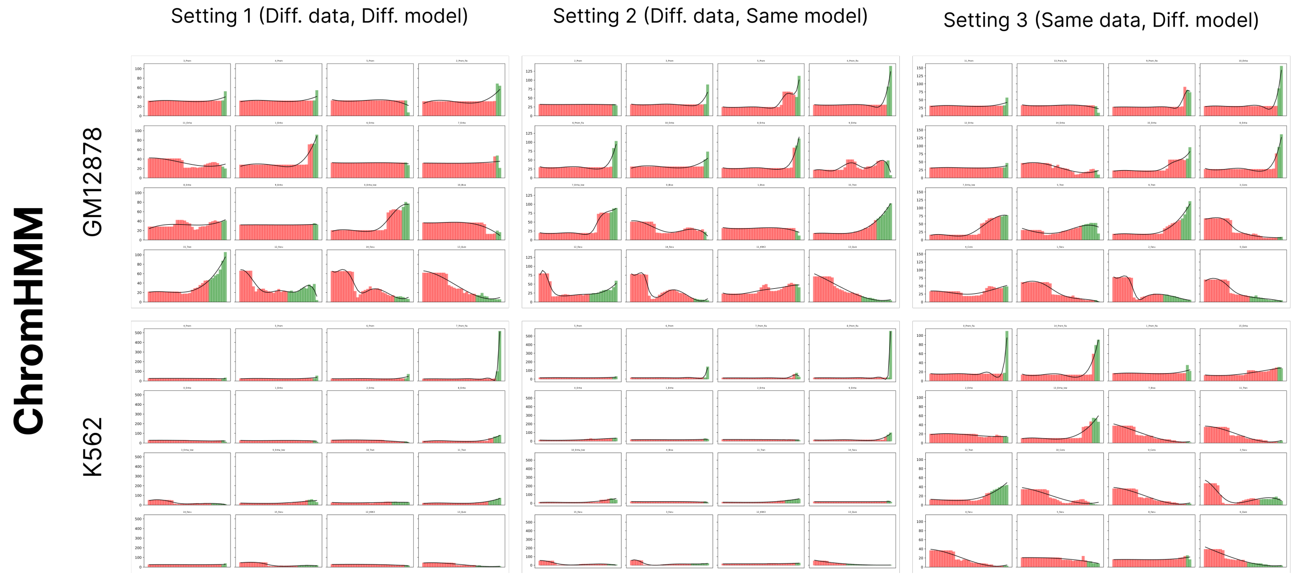


Figure 22: Mean expression level (TPM) in the gene-body as a function of posterior probability for ChromHMM runs in cell types GM12878 and K562. The horizontal axis represents rank of posterior probability, while the vertical axis shows TPM.

##### 5.2.2 Segway

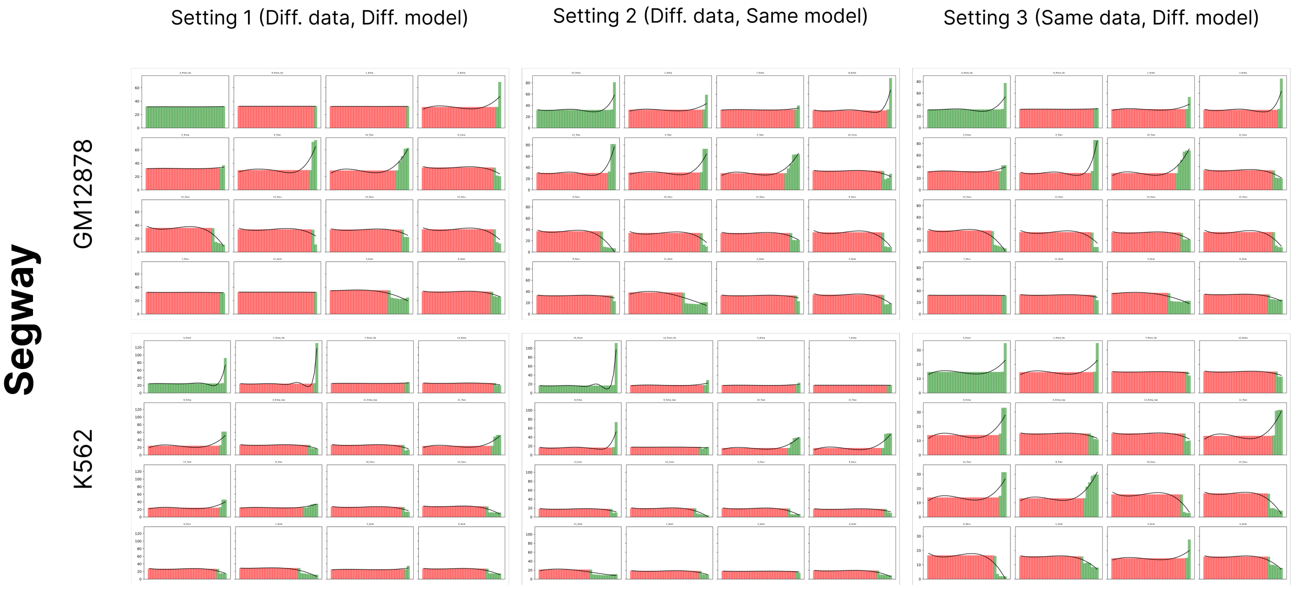


Figure 23: Mean expression level (TPM) in the gene-body as a function of posterior probability for Segway runs in cell types GM12878 and K562. The horizontal axis represents rank of posterior probability, while the vertical axis shows TPM.

#### 5.3 Transcription (around TSS) as a function of posterior probability

##### 5.3.1 ChromHMM

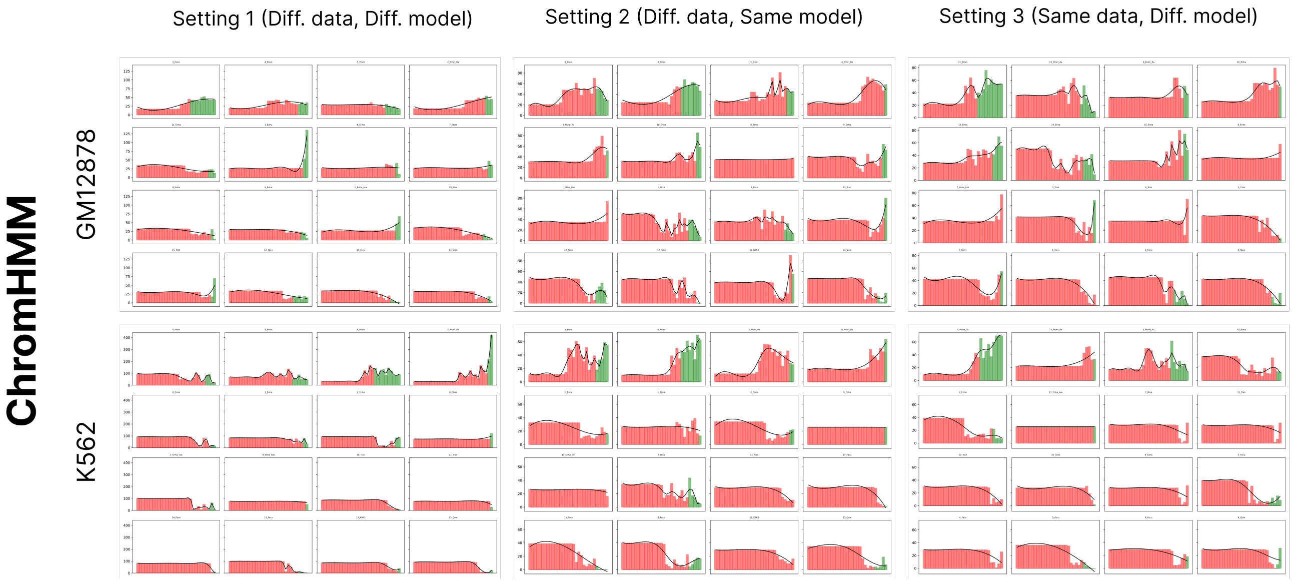


Figure 24: Mean expression level (TPM) around Transcription start sites as a function of posterior probability for ChromHMM runs in cell types GM12878 and K562. The horizontal axis represents rank of posterior probability, while the vertical axis shows TPM.

##### 5.3.2 Segway

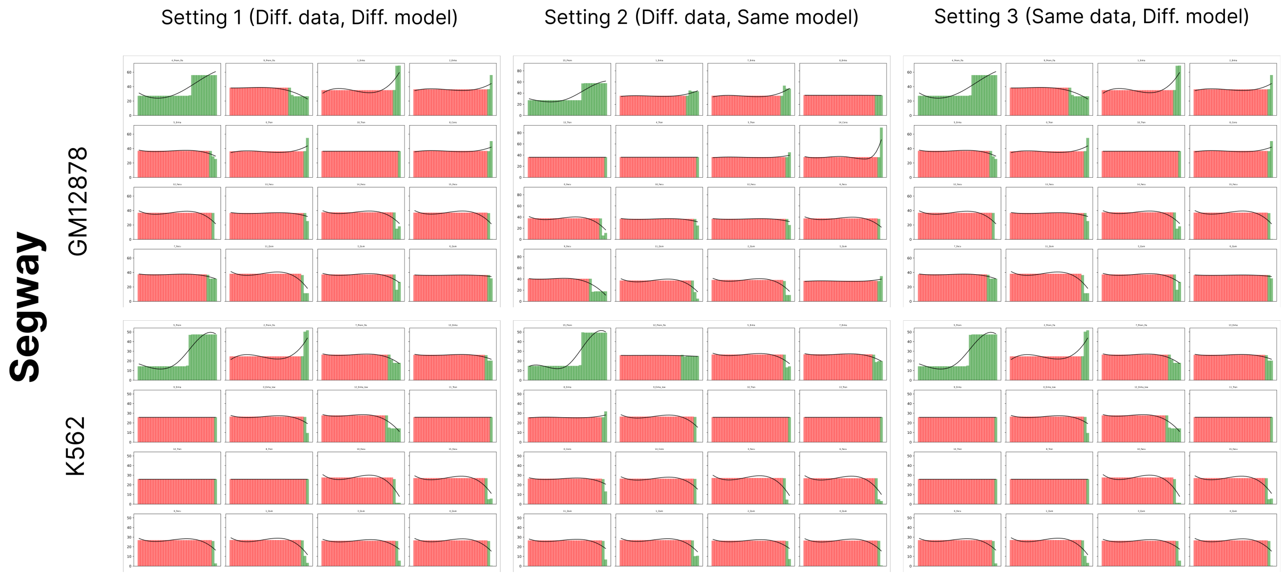


Figure 25: Mean expression level (TPM) around Transcription start sites as a function of posterior probability for Segway runs in cell types GM12878 and K562. The horizontal axis represents rank of posterior probability, while the vertical axis shows TPM.

#### 5.4 TSS enrichment as a function of posterior probability

##### 5.4.1 ChromHMM

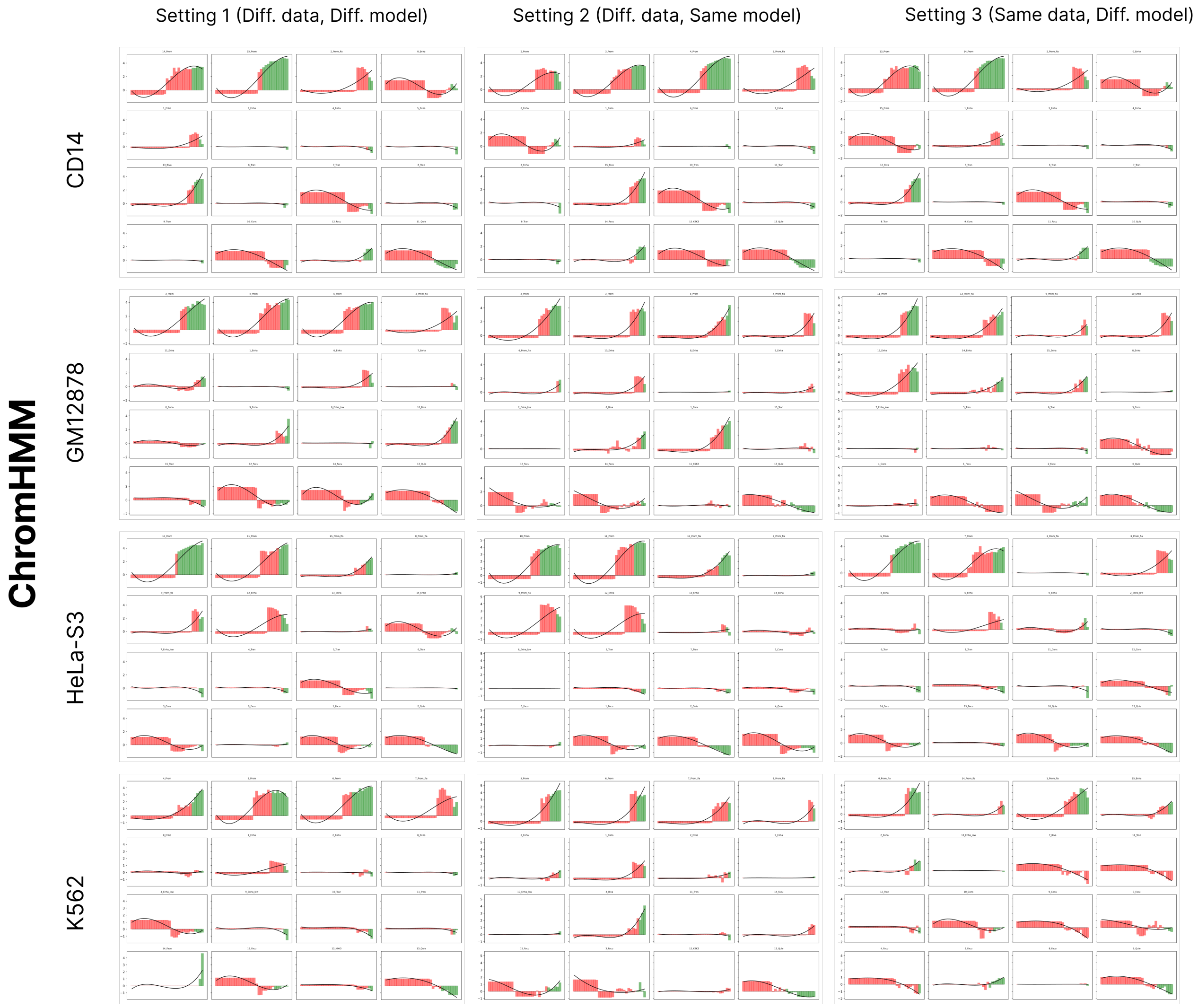


Figure 26: Enrichment of states around Transcription start sites as a function of posterior probability for ChromHMM runs in cell types GM12878 and K562. The horizontal axis represents rank of posterior probability, while the vertical axis shows TPM.

##### 5.4.2 Segway

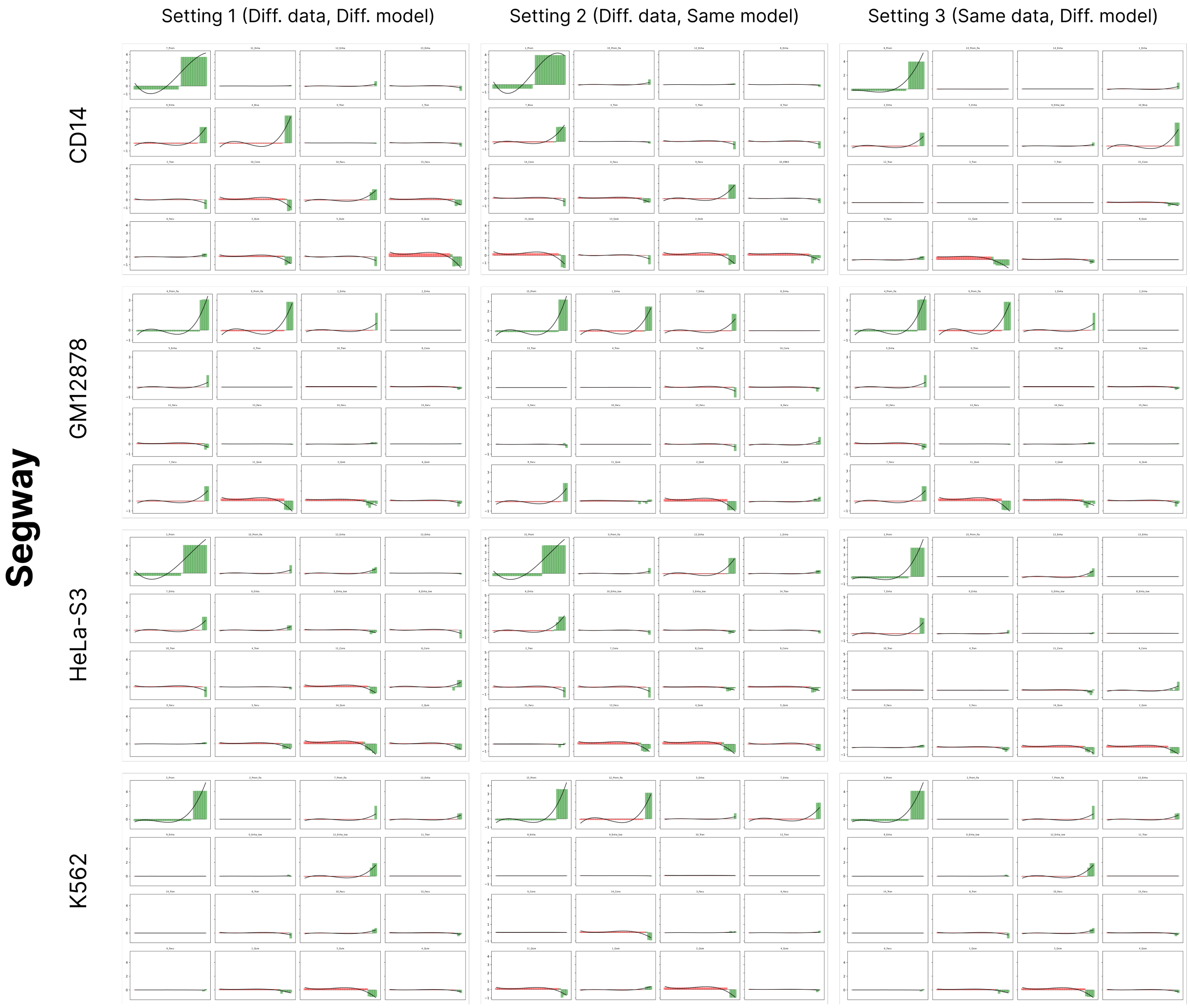


Figure 27: Enrichment of states around Transcription start sites as a function of posterior probability for Segway runs in cell types GM12878 and K562. The horizontal axis represents rank of posterior probability, while the vertical axis shows TPM.

### 6 Reproducibility analysis yields robust annotations

#### 6.1 r-value

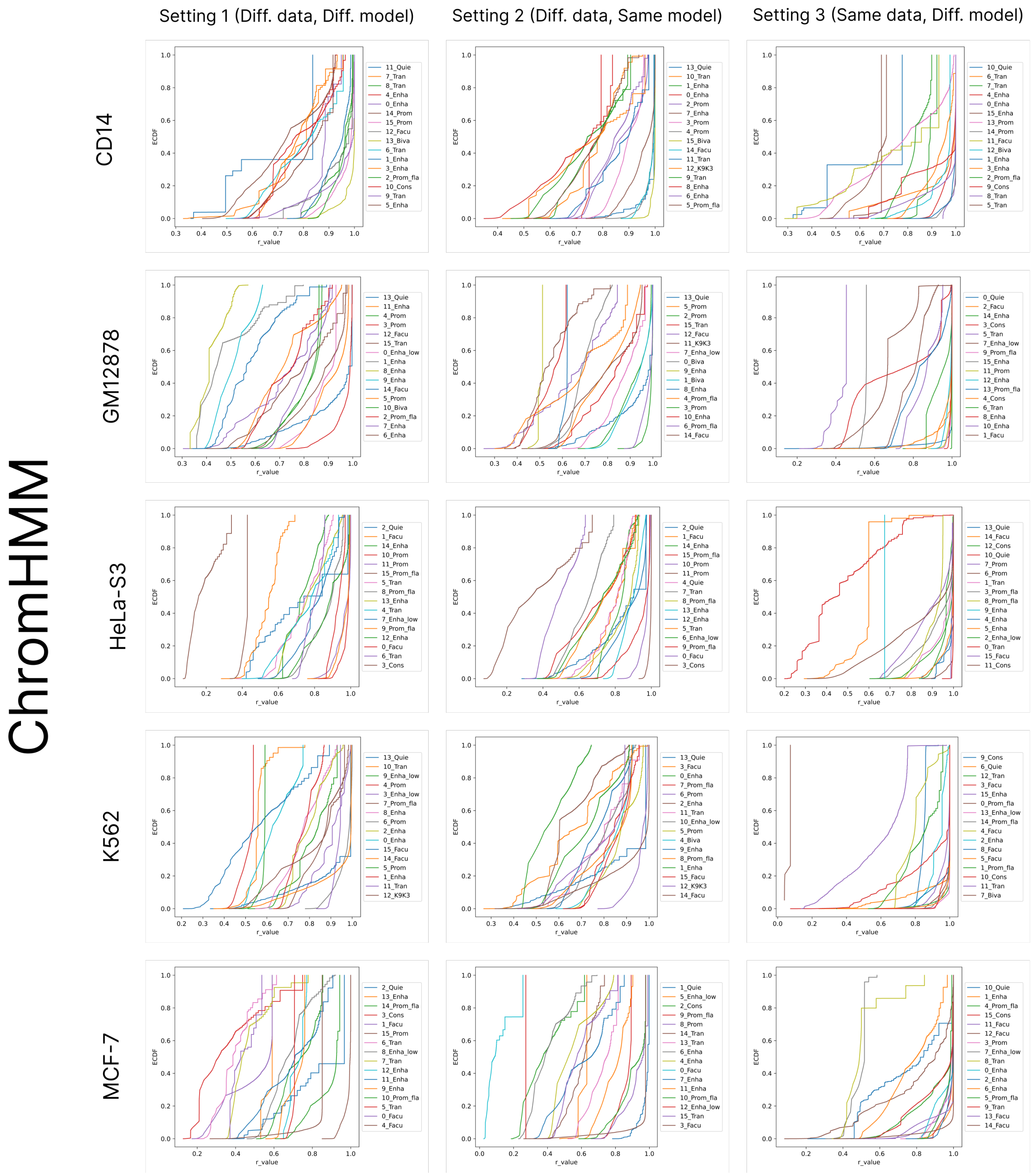


Figure 28: Empirical cumulative distribution function (ECDF) of r-value for all states in ChromHMM runs across five cell types (rows) and three settings of variability (columns).

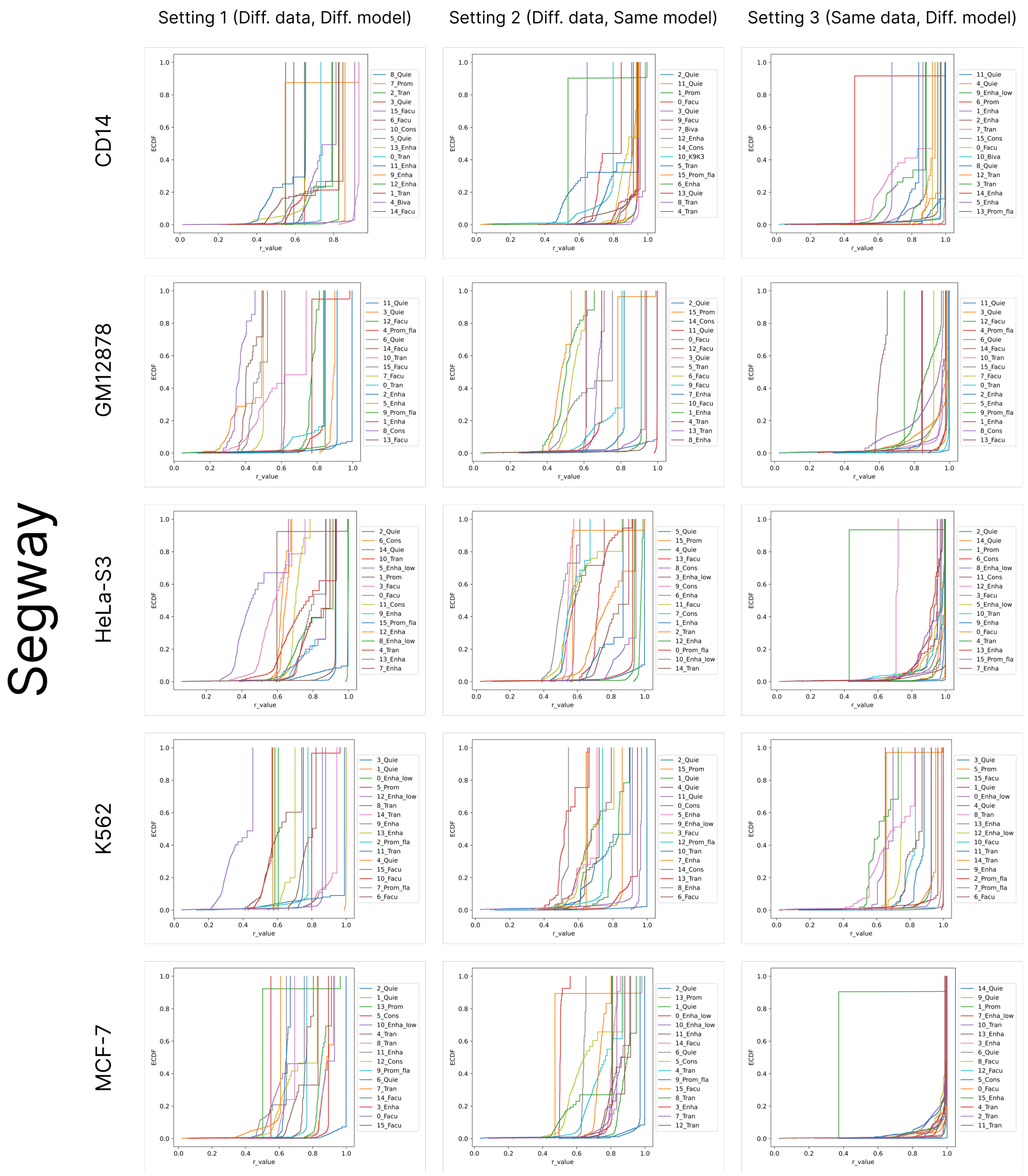


Figure 29: Empirical cumulative distribution function (ECDF) of r-value for all states in Segway runs across five cell types (rows) and three settings of variability (columns).

#### 6.2 Fraction of confident genomic regions

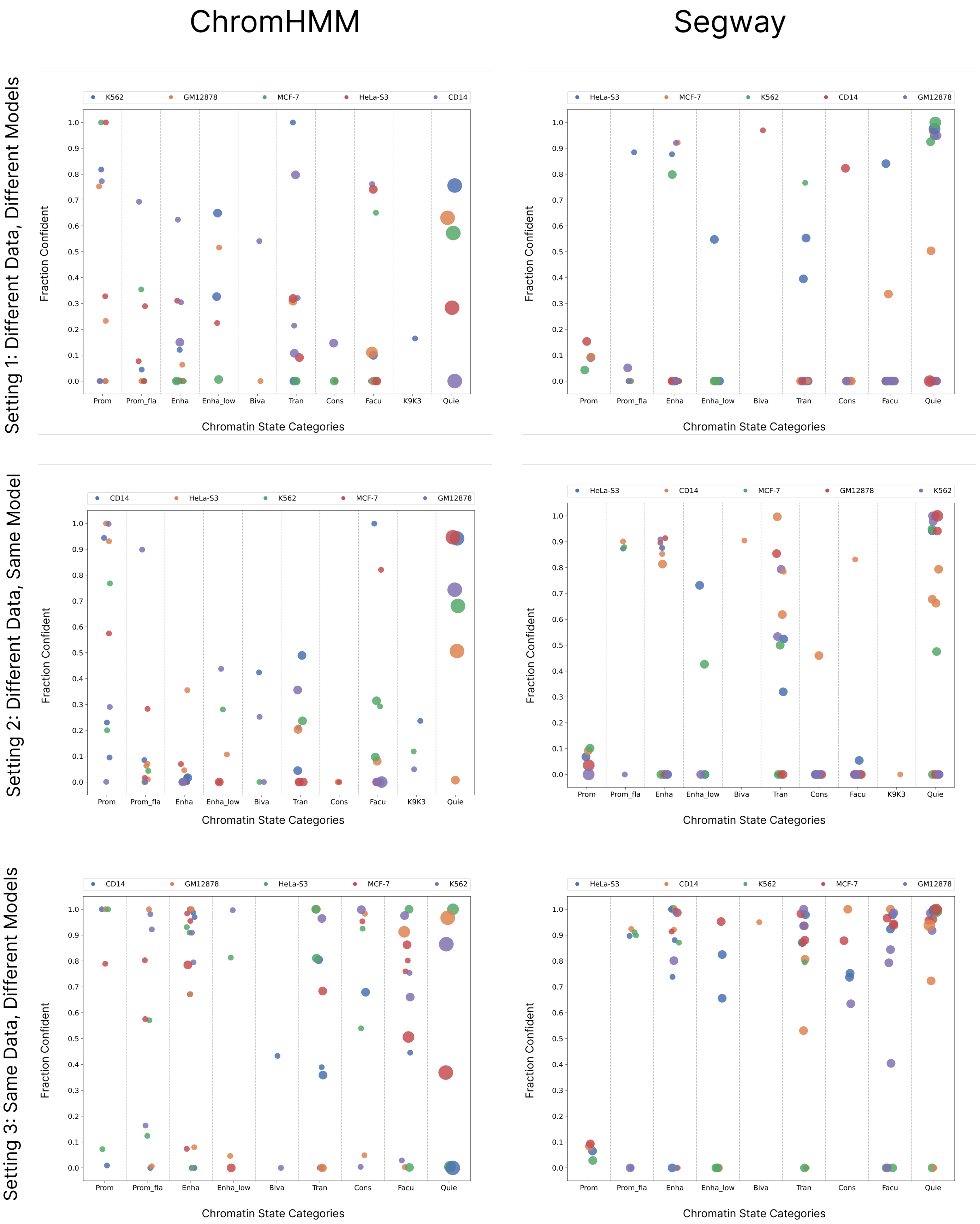


Figure 30: Fraction of confident genomic regions of various chromatin states categories identified in the ChromHMM (left column) and Segway (right column) annotation according to three settings of variability (rows) for five cell types. Each dot represents a chromatin state, with color denoting cell type and size proportional to genome coverage.

### 7 Comparison of Biologically Replicated ChIP-Seq Assays

#### 7.1 CD14-positive-monocyte

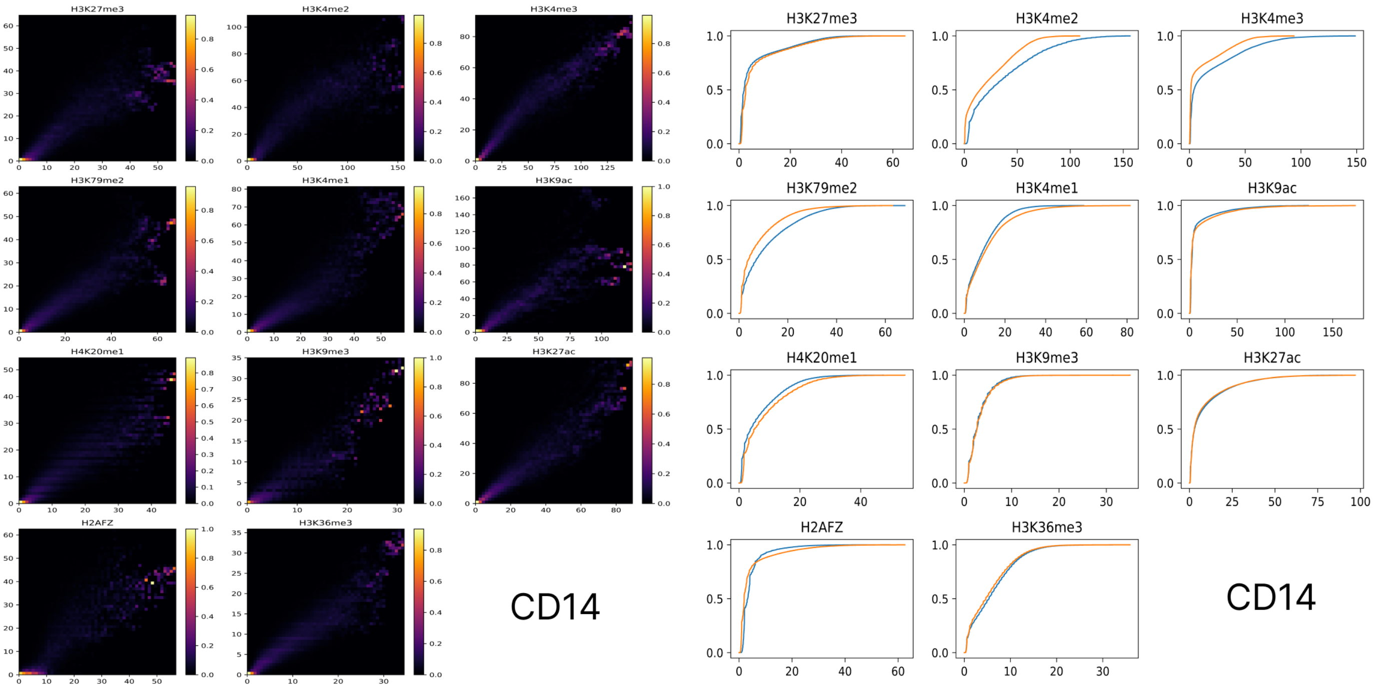


Figure 31: Heatmap of track values across two biological replicates for different histone modification tracks (left). On right, empirical cumulative distribution function (ECDF) of track values across two biological replicates for different histone modification tracks.

## 7.2 GM12878

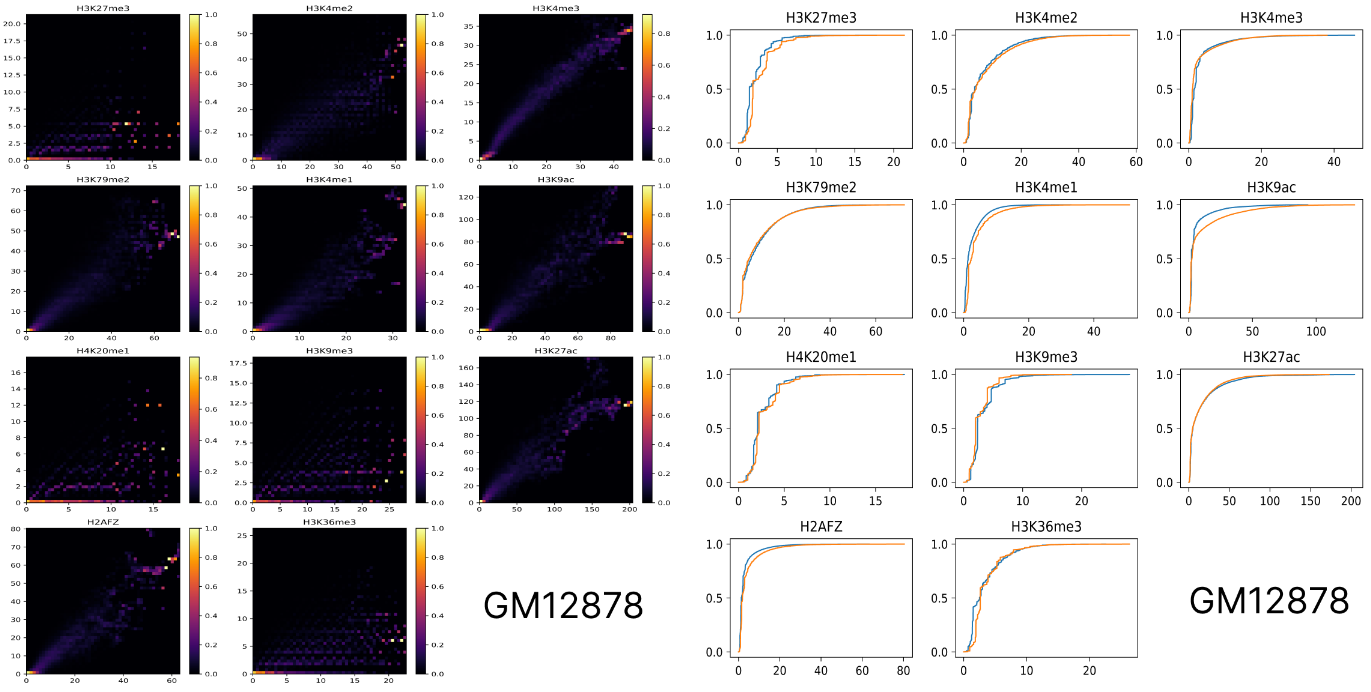


Figure 32: Heatmap of track values across two biological replicates for different histone modification tracks (left). On right, empirical cumulative distribution function (ECDF) of track values across two biological replicates for different histone modification tracks.

#### 7.3 HeLa-S3

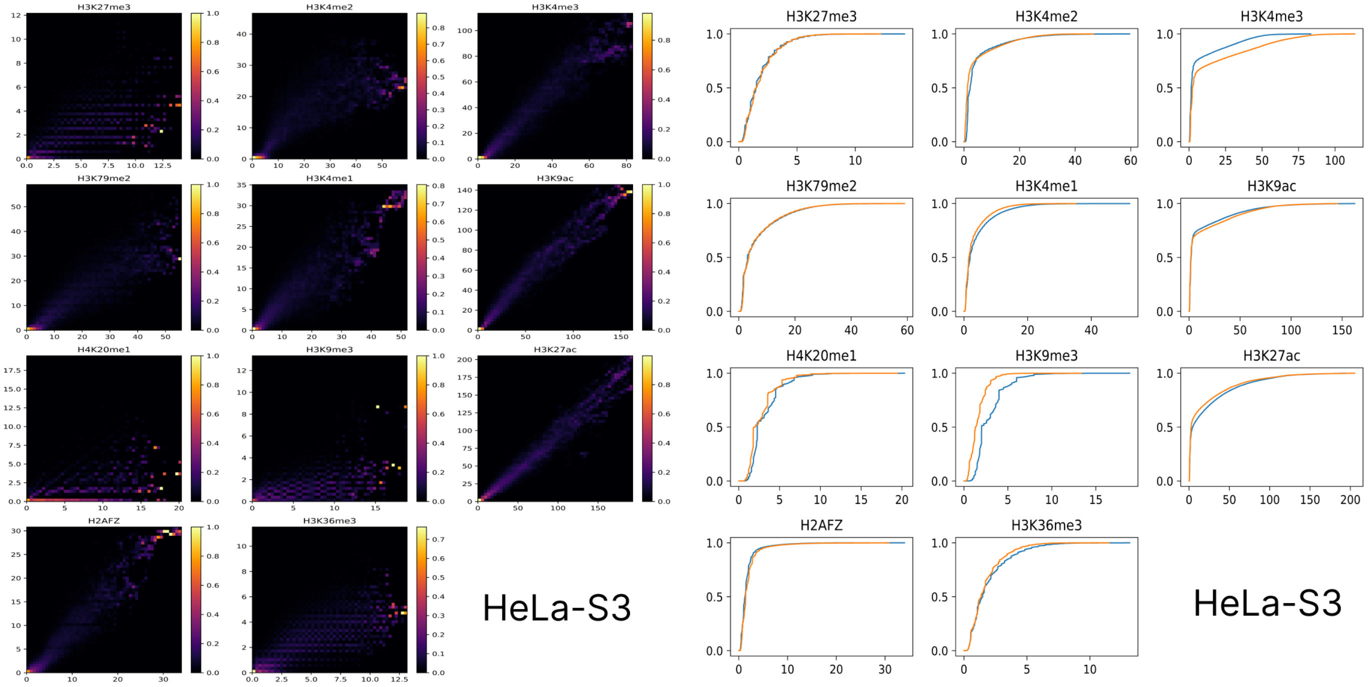


Figure 33: Heatmap of track values across two biological replicates for different histone modification tracks (left). On right, empirical cumulative distribution function (ECDF) of track values across two biological replicates for different histone modification tracks.

## 7.4 K562

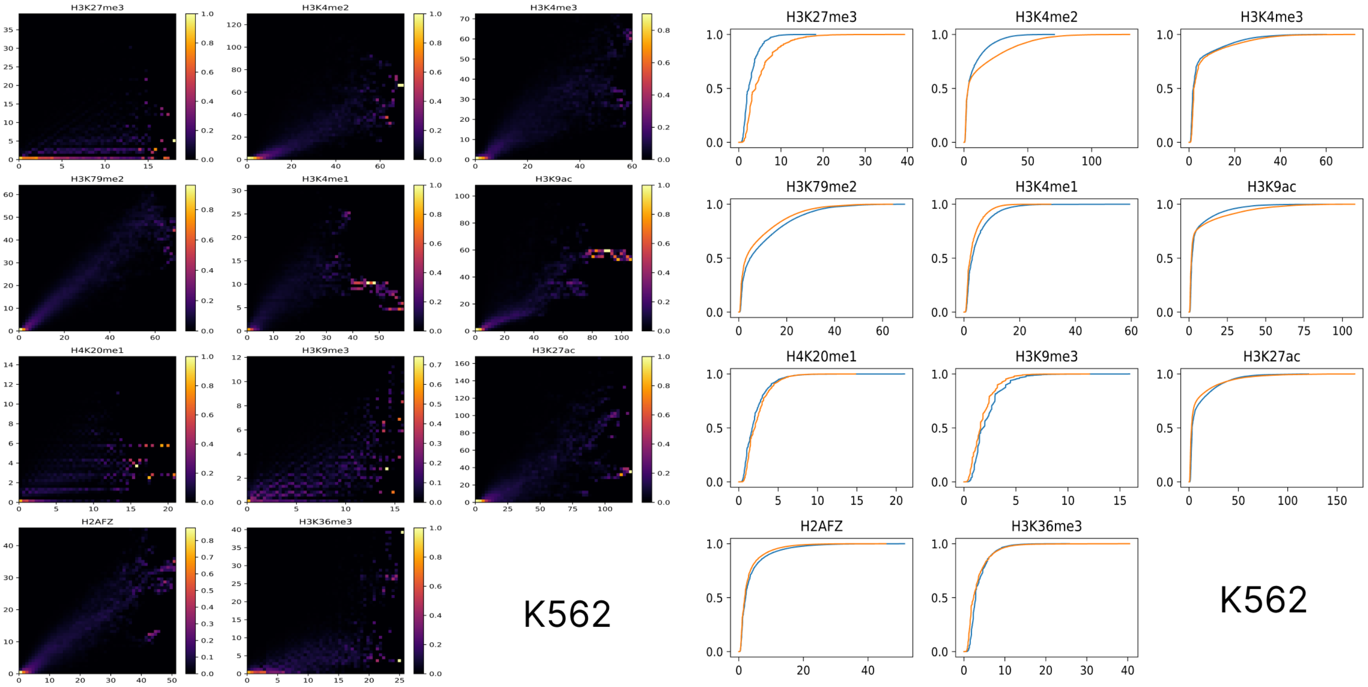


Figure 34: Heatmap of track values across two biological replicates for different histone modification tracks (left). On right, empirical cumulative distribution function (ECDF) of track values across two biological replicates for different histone modification tracks.

#### 7.5 MCF-7

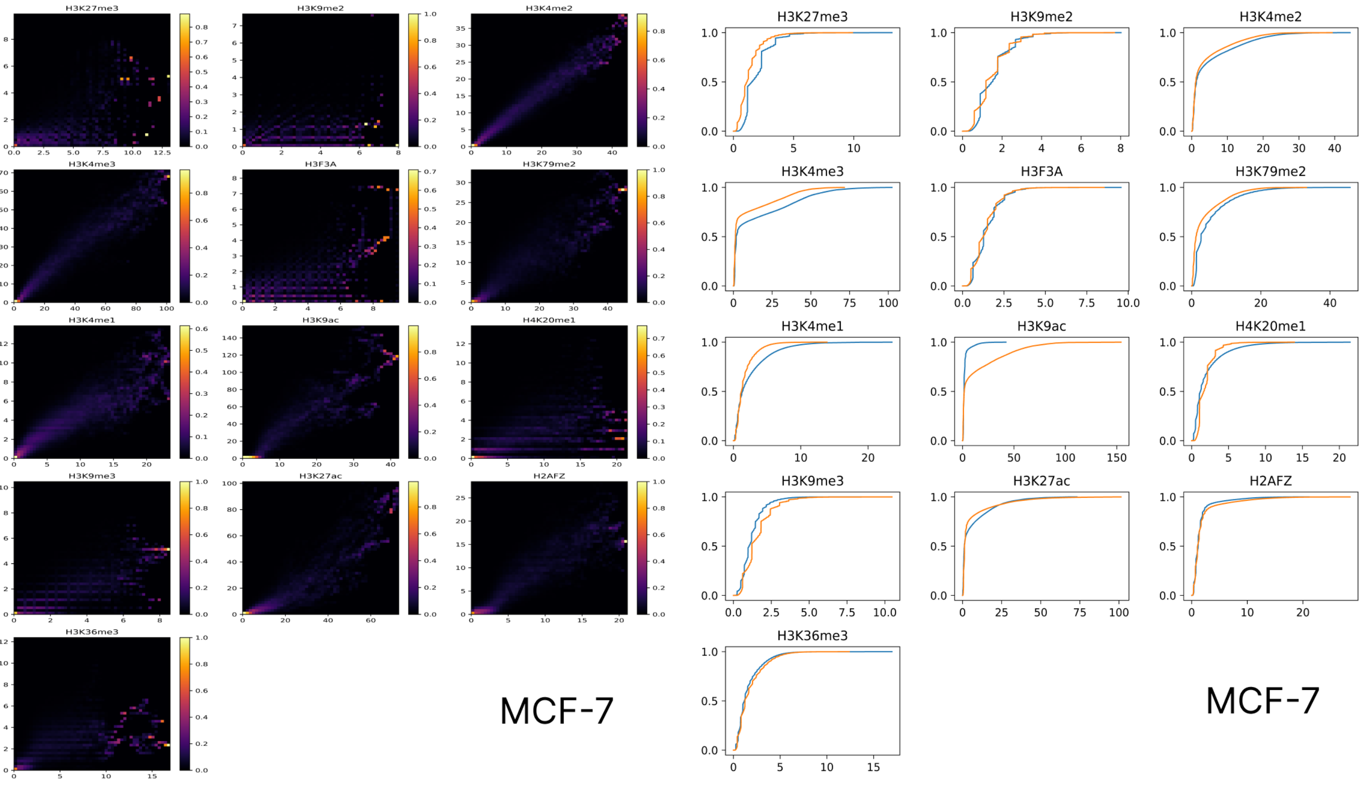


Figure 35: Heatmap of track values across two biological replicates for different histone modification tracks (left). On right, empirical cumulative distribution function (ECDF) of track values across two biological replicates for different histone modification tracks.

### 8 Data Summary

Table 2: This table contains the summary of data retrieved from ENCODE consortium [7] and used to train both ChromHMM and Segway. For each experiment, two biosamples, each corresponding to an isogenic replicate, has been used. For each biosample, two files were retrieved, the aligned reads BAM file and the fold-change over control signal file respectively.

| Cell type | Experiment | Biosample | Files |
|---|---|---|---|
| MCF-7 | H3K27me3 | ENCBS789UPK (rep. 1) | ENCFF413QYQ, ENCFF222KPM |
| MCF-7 | H3K27me3 | ENCBS967MVZ (rep. 2) | ENCFF744JQU, ENCFF070PLQ |
| MCF-7 | H3K9me2 | ENCBS789UPK (rep. 1) | ENCFF249TCH, ENCFF727FOP |
| MCF-7 | H3K9me2 | ENCBS967MVZ (rep. 2) | ENCFF418SFP, ENCFF727JSD |
| MCF-7 | H3K4me2 | ENCBS789UPK (rep. 1) | ENCFF134YPW, ENCFF095LXY |
| MCF-7 | H3K4me2 | ENCBS967MVZ (rep. 2) | ENCFF680ERD, ENCFF930AIB |
| MCF-7 | H3K4me3 | ENCBS789UPK (rep. 1) | ENCFF101TRI, ENCFF559CLH |
| MCF-7 | H3K4me3 | ENCBS967MVZ (rep. 2) | ENCFF374GQR, ENCFF763FZF |
| MCF-7 | H3F3A | ENCBS789UPK (rep. 1) | ENCFF626ORM, ENCFF513VYY |
| MCF-7 | H3F3A | ENCBS967MVZ (rep. 2) | ENCFF183HME, ENCFF820MMV |
| MCF-7 | H3K79me2 | ENCBS789UPK (rep. 1) | ENCFF312WJY, ENCFF921ILS |
| MCF-7 | H3K79me2 | ENCBS967MVZ (rep. 2) | ENCFF654NDM, ENCFF288DKP |
| MCF-7 | H3K4me1 | ENCBS789UPK (rep. 1) | ENCFF592EVS, ENCFF885FSN |
| MCF-7 | H3K4me1 | ENCBS967MVZ (rep. 2) | ENCFF748ISL, ENCFF943MKW |
| MCF-7 | H3K9ac | ENCBS789UPK (rep. 1) | ENCFF571MVG, ENCFF967PEQ |
| MCF-7 | H3K9ac | ENCBS967MVZ (rep. 2) | ENCFF403SOH, ENCFF699NOJ |
| MCF-7 | H4K20me1 | ENCBS789UPK (rep. 1) | ENCFF491EQT, ENCFF665VEZ |
| MCF-7 | H4K20me1 | ENCBS967MVZ (rep. 2) | ENCFF741HOA, ENCFF382BVG |
| MCF-7 | H3K9me3 | ENCBS789UPK (rep. 1) | ENCFF909MGB, ENCFF501WFD |
| MCF-7 | H3K9me3 | ENCBS967MVZ (rep. 2) | ENCFF923NDP, ENCFF587KXV |
| MCF-7 | H3K27ac | ENCBS789UPK (rep. 1) | ENCFF785NUK, ENCFF395YPR |
| MCF-7 | H3K27ac | ENCBS967MVZ (rep. 2) | ENCFF391BOB, ENCFF047PAZ |
| MCF-7 | H2AFZ | ENCBS789UPK (rep. 1) | ENCFF597OEX, ENCFF269EZS |
| MCF-7 | H2AFZ | ENCBS967MVZ (rep. 2) | ENCFF687JRM, ENCFF391YLE |
| MCF-7 | H3K36me3 | ENCBS789UPK (rep. 1) | ENCFF747AWB, ENCFF039TSW |
| MCF-7 | H3K36me3 | ENCBS967MVZ (rep. 2) | ENCFF551PNK, ENCFF781COV |
| GM12878 | H3K27me3 | ENCBS715VCP (rep. 1) | ENCFF633BHN, ENCFF119CAV |
| GM12878 | H3K27me3 | ENCBS830CIQ (rep. 2) | ENCFF565DCK, ENCFF736CNQ |
| GM12878 | H3K4me2 | ENCBS715VCP (rep. 1) | ENCFF008LEY, ENCFF028DRR |
| GM12878 | H3K4me2 | ENCBS830CIQ (rep. 2) | ENCFF081ODV, ENCFF922IYX |
| GM12878 | H3K4me3 | ENCBS715VCP (rep. 1) | ENCFF843BWY, ENCFF346GYG |
| GM12878 | H3K4me3 | ENCBS830CIQ (rep. 2) | ENCFF126HII, ENCFF874RRW |
| GM12878 | H3K79me2 | ENCBS715VCP (rep. 1) | ENCFF465NXQ, ENCFF567RPY |
| GM12878 | H3K79me2 | ENCBS830CIQ (rep. 2) | ENCFF967CHL, ENCFF595CSY |
| GM12878 | H3K4me1 | ENCBS715VCP (rep. 1) | ENCFF047NLO, ENCFF785YET |
| GM12878 | H3K4me1 | ENCBS830CIQ (rep. 2) | ENCFF385FLM, ENCFF721OOX |
| GM12878 | H3K9ac | ENCBS715VCP (rep. 1) | ENCFF729HQN, ENCFF609XKG |
| GM12878 | H3K9ac | ENCBS830CIQ (rep. 2) | ENCFF405REQ, ENCFF404TNY |
| GM12878 | H4K20me1 | ENCBS715VCP (rep. 1) | ENCFF132EDT, ENCFF150LOV |
| GM12878 | H4K20me1 | ENCBS830CIQ (rep. 2) | ENCFF334NVA, ENCFF069ZOX |
| GM12878 | H3K9me3 | ENCBS715VCP (rep. 1) | ENCFF306ENK, ENCFF698SKV |
| GM12878 | H3K9me3 | ENCBS830CIQ (rep. 2) | ENCFF889UPU, ENCFF856REE |
| GM12878 | H3K27ac | ENCBS715VCP (rep. 1) | ENCFF269GKF, ENCFF458CRP |
| GM12878 | H3K27ac | ENCBS830CIQ (rep. 2) | ENCFF201OHW, ENCFF716VWO |
| GM12878 | H2AFZ | ENCBS715VCP (rep. 1) | ENCFF703AIM, ENCFF477ZQM |
| GM12878 | H2AFZ | ENCBS830CIQ (rep. 2) | ENCFF256UUQ, ENCFF553GSY |
| GM12878 | H3K36me3 | ENCBS715VCP (rep. 1) | ENCFF353YPB, ENCFF269OIU |
| GM12878 | H3K36me3 | ENCBS830CIQ (rep. 2) | ENCFF677MAG, ENCFF831NQV |
| K562 | H3K27me3 | ENCBS639AAA (rep. 1) | ENCFF392ZKG, ENCFF585RVP |
| K562 | H3K27me3 | ENCBS674MPN (rep. 2) | ENCFF905CZD, ENCFF679YZC |

| | | | |
|---|---|---|---|
| K562 | H3K4me2 | ENCBS639AAA (rep. 1) | ENCFF446FUS, ENCFF229IJV |
| K562 | H3K4me2 | ENCBS674MPN (rep. 2) | ENCFF583GSH, ENCFF386XDJ |
| K562 | H3K4me3 | ENCBS639AAA (rep. 1) | ENCFF564SVK, ENCFF545JLB |
| K562 | H3K4me3 | ENCBS674MPN (rep. 2) | ENCFF685PPQ, ENCFF617VRS |
| K562 | H3K79me2 | ENCBS639AAA (rep. 1) | ENCFF465UWC, ENCFF952GNK |
| K562 | H3K79me2 | ENCBS674MPN (rep. 2) | ENCFF863HNL, ENCFF080CBF |
| K562 | H3K4me1 | ENCBS639AAA (rep. 1) | ENCFF352HXD, ENCFF490RQD |
| K562 | H3K4me1 | ENCBS674MPN (rep. 2) | ENCFF415GHS, ENCFF149FZG |
| K562 | H3K9ac | ENCBS639AAA (rep. 1) | ENCFF149MXA, ENCFF236VCK |
| K562 | H3K9ac | ENCBS674MPN (rep. 2) | ENCFF505NMT, ENCFF650QTM |
| K562 | H4K20me1 | ENCBS639AAA (rep. 1) | ENCFF255QRL, ENCFF091XLR |
| K562 | H4K20me1 | ENCBS674MPN (rep. 2) | ENCFF923YUN, ENCFF006AYV |
| K562 | H3K9me3 | ENCBS639AAA (rep. 1) | ENCFF104THG, ENCFF103IHG |
| K562 | H3K9me3 | ENCBS674MPN (rep. 2) | ENCFF155UQU, ENCFF600IBI |
| K562 | H3K27ac | ENCBS639AAA (rep. 1) | ENCFF121RHF, ENCFF977KGH |
| K562 | H3K27ac | ENCBS674MPN (rep. 2) | ENCFF907MNY, ENCFF745TSK |
| K562 | H2AFZ | ENCBS639AAA (rep. 1) | ENCFF874SMO, ENCFF242WQK |
| K562 | H2AFZ | ENCBS674MPN (rep. 2) | ENCFF007YZT, ENCFF964BCA |
| K562 | H3K36me3 | ENCBS639AAA (rep. 1) | ENCFF880HKV, ENCFF782UNG |
| K562 | H3K36me3 | ENCBS674MPN (rep. 2) | ENCFF272JVI, ENCFF860PQP |
| CD14+ monocyte | H3K27me3 | ENCBS188BKX (rep. 1) | ENCFF630ANH, ENCFF840DRQ |
| CD14+ monocyte | H3K27me3 | ENCBS865RXK (rep. 2) | ENCFF785GAD, ENCFF586PRJ |
| CD14+ monocyte | H3K4me2 | ENCBS188BKX (rep. 1) | ENCFF591CTO, ENCFF572NHK |
| CD14+ monocyte | H3K4me2 | ENCBS865RXK (rep. 2) | ENCFF384IVB, ENCFF031HMX |
| CD14+ monocyte | H3K4me3 | ENCBS188BKX (rep. 1) | ENCFF640GCK, ENCFF367ZHA |
| CD14+ monocyte | H3K4me3 | ENCBS865RXK (rep. 2) | ENCFF144PKE, ENCFF439PJT |
| CD14+ monocyte | H3K79me2 | ENCBS188BKX (rep. 1) | ENCFF463WUU, ENCFF464AAD |
| CD14+ monocyte | H3K79me2 | ENCBS865RXK (rep. 2) | ENCFF159LCH, ENCFF451RBS |
| CD14+ monocyte | H3K4me1 | ENCBS188BKX (rep. 1) | ENCFF573MNS, ENCFF281FSL |
| CD14+ monocyte | H3K4me1 | ENCBS865RXK (rep. 2) | ENCFF141KHX, ENCFF669AVC |
| CD14+ monocyte | H3K9ac | ENCBS188BKX (rep. 1) | ENCFF775XVG, ENCFF835QUH |
| CD14+ monocyte | H3K9ac | ENCBS865RXK (rep. 2) | ENCFF326KCT, ENCFF627PKL |
| CD14+ monocyte | H4K20me1 | ENCBS865RXK (rep. 2) | ENCFF238ZAC, ENCFF938RSM |
| CD14+ monocyte | H4K20me1 | ENCBS188BKX (rep. 1) | ENCFF899MAF, ENCFF657KKL |
| CD14+ monocyte | H3K9me3 | ENCBS188BKX (rep. 1) | ENCFF537BKE, ENCFF264DFY |
| CD14+ monocyte | H3K9me3 | ENCBS865RXK (rep. 2) | ENCFF168UCU, ENCFF726FHO |
| CD14+ monocyte | H3K27ac | ENCBS188BKX (rep. 1) | ENCFF014BCD, ENCFF078DZN |
| CD14+ monocyte | H3K27ac | ENCBS865RXK (rep. 2) | ENCFF872QFO, ENCFF948MET |
| CD14+ monocyte | H2AFZ | ENCBS188BKX (rep. 1) | ENCFF891JRY, ENCFF834IMH |
| CD14+ monocyte | H2AFZ | ENCBS865RXK (rep. 2) | ENCFF992JKA, ENCFF175WGJ |
| CD14+ monocyte | H3K36me3 | ENCBS188BKX (rep. 1) | ENCFF629ODR, ENCFF494ICL |
| CD14+ monocyte | H3K36me3 | ENCBS865RXK (rep. 2) | ENCFF150SGV, ENCFF554VRE |
| HeLa-S3 | H3K27me3 | ENCBS075PNA (rep. 1) | ENCFF299WXR, ENCFF971CNR |
| HeLa-S3 | H3K27me3 | ENCBS655ARO (rep. 2) | ENCFF620CCL, ENCFF837NDL |
| HeLa-S3 | H3K4me2 | ENCBS075PNA (rep. 1) | ENCFF622IIB, ENCFF375VEB |
| HeLa-S3 | H3K4me2 | ENCBS655ARO (rep. 2) | ENCFF706FSX, ENCFF146XSE |
| HeLa-S3 | H3K4me3 | ENCBS075PNA (rep. 1) | ENCFF300IWJ, ENCFF532TVQ |
| HeLa-S3 | H3K4me3 | ENCBS655ARO (rep. 2) | ENCFF766LEF, ENCFF933NWW |
| HeLa-S3 | H3K79me2 | ENCBS075PNA (rep. 1) | ENCFF616FAT, ENCFF976XPV |
| HeLa-S3 | H3K79me2 | ENCBS655ARO (rep. 2) | ENCFF656YCM, ENCFF418AXA |
| HeLa-S3 | H3K4me1 | ENCBS075PNA (rep. 1) | ENCFF826OLG, ENCFF554WAD |
| HeLa-S3 | H3K4me1 | ENCBS655ARO (rep. 2) | ENCFF712AAP, ENCFF257YVT |
| HeLa-S3 | H3K9ac | ENCBS075PNA (rep. 1) | ENCFF113FCH, ENCFF753SWH |
| HeLa-S3 | H3K9ac | ENCBS655ARO (rep. 2) | ENCFF596XSW, ENCFF995MCG |
| HeLa-S3 | H4K20me1 | ENCBS075PNA (rep. 1) | ENCFF278FTJ, ENCFF776IYG |
| HeLa-S3 | H4K20me1 | ENCBS655ARO (rep. 2) | ENCFF310NWI, ENCFF002OOC |
| HeLa-S3 | H3K9me3 | ENCBS075PNA (rep. 1) | ENCFF183EDU, ENCFF746UJB |
| HeLa-S3 | H3K9me3 | ENCBS655ARO (rep. 2) | ENCFF520XFF, ENCFF629AAP |
| HeLa-S3 | H3K27ac | ENCBS075PNA (rep. 1) | ENCFF609ZAE, ENCFF010QGR |
| HeLa-S3 | H3K27ac | ENCBS655ARO (rep. 2) | ENCFF711QAI, ENCFF855XCJ |

| | | | |
|---|---|---|---|
| HeLa-S3 | H2AFZ | ENCBS075PNA (rep. 1) | ENCFF650RBB, ENCFF423XMI |
| HeLa-S3 | H2AFZ | ENCBS655ARO (rep. 2) | ENCFF822OCY, ENCFF104TOM |
| HeLa-S3 | H3K36me3 | ENCBS075PNA (rep. 1) | ENCFF317FFS, ENCFF305UYI |
| HeLa-S3 | H3K36me3 | ENCBS655ARO (rep. 2) | ENCFF008RRT, ENCFF870FDA |

#### References


[1] Hoffman, M.M., Buske, O.J., Noble, W.S.: The genomedata format for storing large-scale functional genomics data. Bioinformatics **26**(11), 1458–1459 (2010)

[2] Hoffman, M.M., Ernst, J., Wilder, S.P., Kundaje, A., Harris, R.S., Libbrecht, M., Giardine, B., Ellenbogen, P.M., Bilmes, J.A., Birney, E., *et al.*: Integrative annotation of chromatin elements from encode data. Nucleic acids research **41**(2), 827–841 (2013)

[3] Chan, R.C., Libbrecht, M.W., Roberts, E.G., Bilmes, J.A., Noble, W.S., Hoffman, M.M.: Segway 2.0: Gaussian mixture models and minibatch training. Bioinformatics **34**(4), 669–671 (2018)

[4] Hoffman, M.M., Buske, O.J., Wang, J., Weng, Z., Bilmes, J.A., Noble, W.S.: Unsupervised pattern discovery in human chromatin structure through genomic segmentation. Nature methods **9**(5), 473–476 (2012)

[5] Libbrecht, M.W., Rodriguez, O.L., Weng, Z., Bilmes, J.A., Hoffman, M.M., Noble, W.S.: A unified encyclopedia of human functional dna elements through fully automated annotation of 164 human cell types. Genome biology **20**(1), 1–14 (2019)

[6] Ernst, J., Kellis, M.: Chromhmm: automating chromatin-state discovery and characterization. Nature methods **9**(3), 215–216 (2012)

[7] Consortium, E.P., *et al.*: An integrated encyclopedia of dna elements in the human genome. Nature **489**(7414), 57 (2012)
